# CREB drives acinar cells to ductal reprogramming and promotes pancreatic cancer progression in preclinical models of alcoholic pancreatitis

**DOI:** 10.1101/2024.01.05.574376

**Authors:** Supriya Srinivasan, Siddharth Mehra, Sudhakar Jinka, Anna Bianchi, Samara Singh, Austin R. Dosch, Haleh Amirian, Varunkumar Krishnamoorthy, Iago De Castro Silva, Manan Patel, Edmond Worley Box, Vanessa Garrido, Tulasigeri M. Totiger, Zhiqun Zhou, Yuguang Ban, Jashodeep Datta, Michael VanSaun, Nipun Merchant, Nagaraj S. Nagathihalli

**Affiliations:** Department of Surgery, University of Miami Miller School of Medicine, Miami, Florida; Department of Public Health Sciences, University of Miami Miller School of Medicine, Miami, Florida; Sylvester Comprehensive Cancer Center, University of Miami, Miami, Florida; Department of Cancer Biology, University of Kansas Medical Center, Kansas City, Kansas

**Keywords:** cAMP response element binding protein 1, acinar-to-ductal metaplasia, alcoholic chronic pancreatitis, pancreaticintraepithelial neoplasia, pancreatic ductal adenocarcinoma, pancreaticcancer

## Abstract

**BACKGROUND & AIMS:** Chronic alcoholism often leads to pancreatitis, which exacerbates pancreatic damage through acinar cell injury, fibrotic inflammation and activates AKT/mTOR/cyclic adenosine monophosphate response element binding protein 1 (CREB) signaling axis. However, the molecular interplay between oncogenic *Kras^G12D/+^*(*Kras\**) and CREB in promoting pancreatic cancer progression under chronic inflammation remains poorly understood.

**METHODS:** Experimental alcoholic chronic pancreatitis (ACP) induction was established in multiple mouse models, with euthanasia during the recovery stage to evaluate tumor latency. CREB was selectively deleted (*Creb^fl/fl^*) in *Ptf1a^CreERTM/+^;LSL-Kras^G12D/+^*(*KC*) genetic mouse models (*KCC^-/-^*). Pancreata from *Ptf1a^CreERTM/+^*, *KC*, and *KCC^-/-^* mice were analyzed using histological profiling, western blotting, phosphokinase array, and quantitative PCR. Single-cell RNA sequencing was performed in ACP-induced *KC* mice. Lineage tracing analysis in YFP reporter mice and acinar cell explant cultures analysis were also conducted.

**RESULTS:** ACP induction in *KC* mice significantly impaired pancreas’ repair mechanism. Acinar cell-derived ductal lesions demonstrated sustained CREB hyperactivation in acinar-to-ductal metaplasia (ADM)/pancreatic intraepithelial neoplasia (PanIN) lesions associated with pancreatitis and pancreatic cancer. Persistent CREB activity reprogrammed acinar cells, and increased profibrotic inflammation. Notably, acinar specific *Creb* deletion in ACP induced models suppressed high grade PanIN development, restrained tumor progression, and improved acinar cell function.

**CONCLUSIONS:** Our findings demonstrate that CREB and *Kras\** promote irreversible ADM, accelerating pancreatic cancer progression with ACP. Targeting CREB may present a promising strategy to mitigate inflammation-driven pancreatic tumorigenesis.

## Introduction

Alcoholic pancreatitis, both acute and chronic, is a risk factor for pancreatic cancer.^1, 2^ Prolonged alcohol abuse accounts for 60–90% of chronic pancreatitis (CP) cases. ^3, 4^ Continuous inflammation due to alcoholic CP (ACP) causes pancreatic atrophy and fibrosis within and around the pancreas.^5^

Chronic pancreatic inflammation frequently involves loss of acinar cell homeostasis and ductal phenotype acquisition, termed acinar-to-ductal metaplasia (ADM). This adaptive physiological response to inflammation protects the pancreas from further damage.^6^ Nevertheless, scientific experimental models consistently demonstrate that in pancreatic tissue with oncogene *Kras^G12D/+^* (*Kras\**), inflammation conspired by non genetic environmental factors including excess alcohol consumption, obesity and smoking impedes pancreatic regeneration and expedites ^7–9^ the development of neoplastic precursor lesions, including pancreatic intraepithelial neoplasia (PanIN), and may progress rapidly toward invasive carcinoma.^10, 11^

Previous studies using caerulein and ACP mouse models have linked epithelial cell intrinsic-signaling (MAPKs, PI3K/AKT, and mTOR) to inflammation driven pancreatic disease pathogenesis. ^12–14^ Building on our lab’s long standing efforts in understanding the multifaceted role of transcription factor signaling in pancreatitis and pancreatic cancer, we have previously identified hyperactivation of cyclic adenosine monophosphate response element-binding protein 1 (CREB) via granulocyte-macrophage colony stimulation factor (GM-CSF) as a key mediator of chronic inflammation that is induced by smoking and regulates aggressiveness of pancreatic cancer.^15^ In our recent study using a C57BL/6 mouse model of alcohol-induced chronic inflammation, unbiased screening of specific kinase signaling pathways identified CREB phosphorylation at Ser133 as one of the highest upregulated target along side PI3K/AKT. ^16^ Therapeutic targeting of PI3K/AKT signaling node mitigates the severity and progression of CP, primarily by modulating acinar cell death,^17^ however, the role of transcription factors including CREB in acinar-to-ductal reprogramming in the context of ACP has not been explored before. Despite the well-established relationship between inflammation and cancer, the molecular mechanisms linking chronic inflammation with oncogenic drivers remain poorly understood within the context of pancreatic cancer initiation and progression. Since a majority of patients with chronic pancreatitis (CP) are alcoholics, this study aimed to develop pancreas specific genetically engineered mouse models (GEMMs) to better investigate these mechanisms.

Herein, we demonstrate persistent CREB activation in acinar cells transitioning toward a ductal phenotype in *Kras\** (*KC*) using GEMMs post-ACP induction. *Creb* deletion in *KC* mice with or without ACP reduced ADM reprogramming, hindered tumor growth and prolonged tumor latency. These findings establish CREB as a key oncogenic role in *Kras-*mediated pancreatic cancer progression with chronic inflammation and also underscores its potential as a promising therapeutic target for pancreatic cancer.

## Results

### Establishing ACP in Ptf1a^CreERTM/+^ Mice

An ACP induction experimental mouse model was established in *Ptf1a^CreERTM/+^*mice, as described previously.^18^ The mice were sacrificed after 3 or 21 days (ACP recovery period) (Figure 1*A*). All experimental cohort mice exhibited weight gain post ACP induction (Supplementary Figure 1A), however, stopping exogenous ACP induction restored the mouse’s body weight (identical to the control) within 21 days during the ACP recovery period. Serum levels revealed an increased blood alcohol concentration with A or ACP induction (Supplementary Figure 1B). *Ptf1a^CreERTM/+^*mice with ACP induction demonstrated markedly reduced pancreatic weight (implying pancreatic injury) (Supplementary Figure 1C). ACP withdrawal completely restored pancreatic weight to its normal level after 21 days (Supplementary Figure 1D). Alcohol-treated mice showed elevated serum amylase levels whereas experimental mice cohort with ACP induction exhibited the most substantial decline in serum amylase levels among the control, A, and CP mice (Supplementary Figure 1E), suggesting maximum acinar injury. Histological assessment of mouse pancreas harvested 3 days after ACP induction revealed profound injuries to the pancreatic tissue architecture, including acinar cell loss, duct-like structures (CK-19^+^ positivity), activated pancreatic stellate cells (PSCs) displaying alpha-smooth muscle actin positivity, increased collagen deposition (Sirius Red) [Supplementary Figure 1F], and leukocyte infiltration (CD45^+^ immune cells) compared with the control, A-, or CP-exposed mice (Figure 1*B* and Supplementary Figure 1F). Pancreatic injury in *Ptf1a^CreERTM/+^*mice completely reversed after discontinuing the exogenous ACP stimulus (21-day recovery period) (Figure 1*B*). No histologically detectable PanINs occurred (Figure 1*B* and Supplementary Figure 1F) in any mouse cohorts. Quantitative PCR (qPCR)-based transcriptional profiling of ACP-treated mouse pancreata demonstrated marked downregulation of genes implicated in acinar cell regulation and function (Supplementary Figure 2A). Overall, these results communicate the transient and reversible nature of pancreatic acinar injury after ACP induction in *Ptf1a^CreERTM/+^*mice.

**Figure 1.**
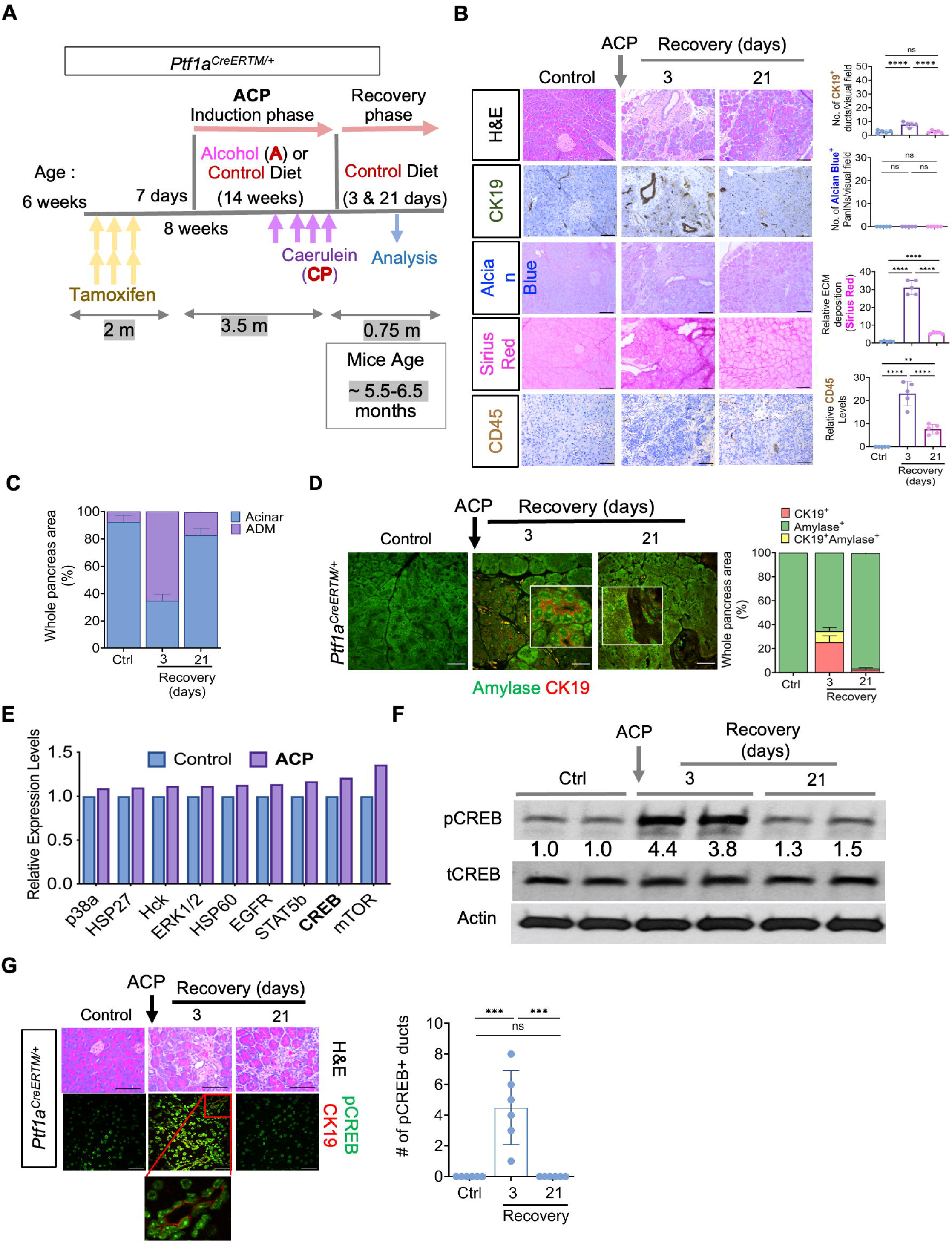
CREB activation in alcoholic chronic pancreatitis (ACP) in *Ptf1a^CreERTM^*^/+^ mice. (A) ACP induction in *Ptf1a^CreERTM^*^/+^ mice exposed to ethanol (alcohol)-enriched liquid diet (A) and caerulein administration (CP). Mice were euthanized after 3- and 21-day recovery periods. (B) Mouse pancreas with H&E and quantification depicting ducts (CK19^+^), PanINs (Alcian Blue), collagen (Sirius red), and immune cells (CD45^+^) (n=5 mice per group). (C) H&E-based quantification of mice pancreata highlighting acinar and ADM regions (n=3 mice per group). (D) Representative images of pancreas depicting amylase (green)/CK19 (red) co-I.F labeling and CK19^+^ amylase^+^ cell corresponding quantification (n=3 mice per group). (E) Mouse kinase array in control (ctrl) and ACP-induced pancreata performed by using pooled tissue lysates from (N=3) biological replicates for each group (Ctrl and ACP), with two membranes used for quantitative estimation of fold change differences. (F) Western blotting showing pCREB expression with relative fold change expression values (normalized to total CREB) in pancreatic tissue lysates in ctrl, 3- and 21-day ACP recovery period (n=2 mice per group) (G) H&E and pCREB (green)/CK19 (red) co-I.F labeling with pCREB^+^ duct quantification in pancreata harvested from ctrl and ACP with recovery (n=6 mice per group). Scale bar, 50 μm. ^ns^=nonsignificant; **p<0.01; ***, p<0.001; ****p<0.0001 by ANOVA.

### Activation of (CREB) is a Hallmark of ADM in Response to ACP

Hematoxylin and eosin (H&E) (Figure 1*C*) and co-immunofluorescence (I.F) (Figure 1*D*) staining for CK19 and amylase in ACP-treated *Ptf1a^CreERTM/+^* mice revealed transient duct-like cells within 3 days of ACP induction. By the end of the 21-day recovery period, these cells redifferentiated and repopulated the acinar compartment, demonstrating the reversibility of ADMs.

We conducted phosphokinase array profiling on *Ptf1a^CreERTM/+^*mice pancreata post-ACP induction. Our analysis revealed mTOR and CREB as the two highest upregulated signaling targets, compared to the control pancreatic lysates (Figure *1E* and Supplementary Figure 2B). This study explores CREB’s role in acinar to ductal reprogramming with *Kras\**. Further validation using immunoblotting and co-I.F staining analysis uncovered elevated pCREB expression in duct-like structures 3 days post-ACP recovery, compared with the controls (Figure 1*F* and *G*). By day 21 of ACP recovery, pCREB expression returned to baseline levels. These findings suggest that ACP induces pancreatic injury, fosters a fibroinflammatory milieu, and enhances CREB activation. Notably, these changes reversed during the recovery period, reducing CREB activation to baseline.

### ACP Induction Accelerates ADM Reprogramming to PanIN and Pancreatic Cancer in Kras^G12D^ Mutant Mice

To enhance the evidence of ACP-mediated acinar cell reprogramming toward the ductal phenotype, we used *KC* GEMM harboring *Kras\** and established ACP induction (described in Figure 1*A*). Mice receiving an alcohol-based diet showed higher blood alcohol concentrations than control mice (Supplementary Figure 3A). Inducing ACP in *KC* mice considerably increased relative pancreatic weight (Figure 2*A*), without significant sex-specific differences (Supplementary Figure 3B).

**Figure 2.**
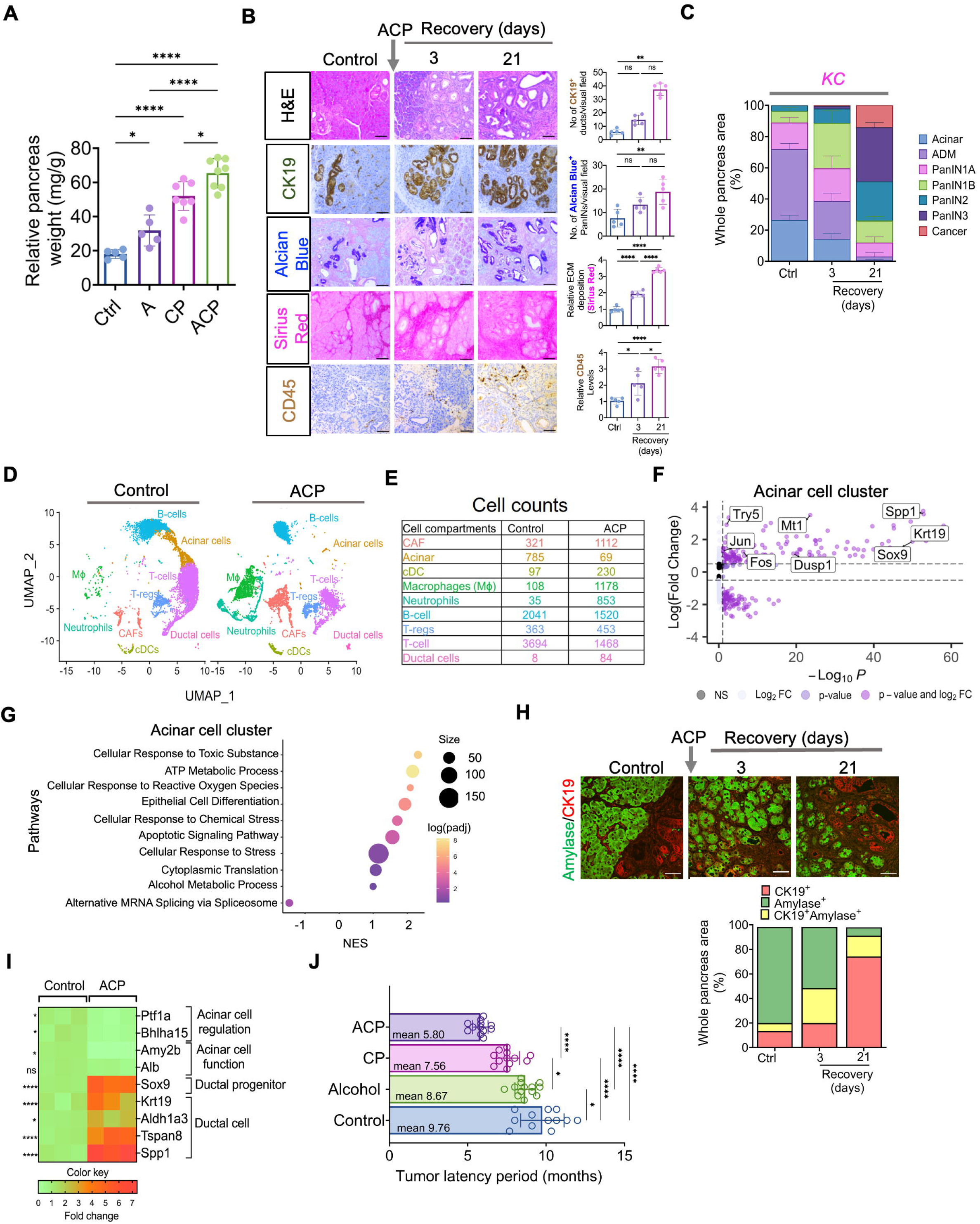
ACP induction accelerates ADM reprogramming toward PanIN and pancreatic cancer in *Kras*^G12D/+^ mutant mice. (A) Relative pancreas weight measurements in respective *KC* mice cohorts (n=5–8 mice per group). (B) Representative pancreatic images depicting H&E with quantification of CK19^+^ ducts, PanINs, collagen, and CD45^+^ in ctrl and ACP-induced *KC* mice with recovery periods (n=5 mice per group). (C) H&E staining within the pancreas depicting acinar, ADMs, PanINs, and cancer regions (n=3 mice per group). (D) UMAP projection displaying cell clusters and (E) cell count table, (F) volcano plot and (G) bubble plot within the acinar cells in the scRNA sequencing analysis of pancreata from ctrl and ACP-*KC* mice. (H) Representative pancreas images depicting amylase (green)/CK19 (red) co-I.F (*top*) CK19^+^ amylase^+^ cell (*bottom*) quantification (within the epithelial cell compartment only) in ctrl and ACP-induced *KC* mice with 3- and 21-day recovery periods (n=3 mice per group). (I) qPCR analysis in the pancreas of ctrl and ACP-induced *KC* mice (n=3 mice per group). (J) Measurement of tumor latency period in ctrl, A, CP, and ACP-induced *KC* mice (n=12–13 mice per group). Scale bar, 50 μm. ^ns^=nonsignificant; *p<0.05; **p<0.01; ****p<0.0001 by ANOVA or unpaired t-test.

The pancreas in control *KC* mice exhibited well-structured acinar compartments and low-grade PanIN lesions (Figure 2*B* and *C*). In contrast, ACP-induced *KC* mice demonstrated a substantially higher frequency of mucinous ductal lesions (low-grade PanINs) with fewer acinar cells (after a 3-day recovery period) than A- or CP-induced *KC* mouse pancreata (Figure 2*C* and Supplementary Figure 3C and 3D). After ACP induction and 21-day recovery, pancreatic tissue histological analysis revealed that these mucinous PanIN progressed towards high-grade lesions (Figure 2*B* and *C*) and a significant increase in CK-19^+^ ductal positivity, mucin content, increased collagen deposition, and infiltration of CD45^+^ immune cells was noted (Figure 2*B*). This increase along with activated PSCs were higher in the ACP experimental cohort of *KC* mice pancreata than in A- or CP mice (Supplementary Figure 3C). A substantial increase in the deposition of extracellular matrix and increased immune cells infiltration accompanied this progression (Figure 2*B* and Supplementary Figure 3C). Chronic inflammatory stimuli induced by ACP resulted in significantly elevated pCREB expression compared to control or alcohol alone, suggesting a potential association with increased pancreatic damage and heightened CREB (Supplementary Figure 3E). Interestingly, chronic inflammatory insult due to CP alone also led to increased CREB activation compared to control *KC* mice (Supplementary Figure 3E). Although, pCREB expression appeared to be higher in ACP induced mice compared to CP (Supplementary Figure 3E). Overall, these findings suggest that the addition of alcohol to caerulein-induced CP amplifies fibroinflammatory milieu - enhanced CREB activation to accelerate tumor progression in *KC* mice pancreata.

Single cell RNA-seq (scRNA-seq) was performed on live single-cell suspensions of *KC* mice pancreata, either control or ACP. Overall, 7903 and 7797 cells from the control and ACP groups were analyzed, respectively. Uniform Manifold Approximation and Projection (UMAP) cell clusters were annotated by examining classic cell type markers.^19, 20^ We distinguished nine distinct clusters representing different cell types (Figure 2*D* and Supplementary Figure 3F). ACP-induced *KC* mice exhibited an increased abundance of cell types, including ductal cells, macrophages, neutrophils, regulatory T cells, and fibroblasts, compared with the control mice (Figure 2*E*). Transcriptional analysis of the acinar cell compartment of *KC* mice pancreas showed upregulation of mRNA transcripts related to inflammation-induced stress, growth, and proliferation with ACP induction compared with the control mice (Figure 2*F*). Gene Ontology Biological Process (GOBP) analysis revealed that genes in the acinar subcluster exhibited differential expression related to cellular stress, apoptosis, alcohol metabolism response, and epithelial cell differentiation, consistent with ACP induction in *KC* GEMM mice (Figure 2*G*). The uninjured control *KC* pancreata showed more amylase^+^ acinar cells with CK19^+^ expression associated with the ductal structures, as revealed using co-I.F analysis (Figure 2*H*). In ACP-treated *KC* mice, the acinar compartment progressively transformed and was replaced by a substantial proportion of ductal structures of acinar origin (dual CK19^+^amylase^+^ cells) after 3 days of recovery. Pancreatic tissue analysis at the end of the 21-day recovery period revealed a ductal phenotype with the highest proportion of CK19^+^ cells (Figure 2*H*). Notably, in ACP induced *KC* mice, pancreata amylase distribution was predominantly localized to the basolateral membrane, contrary to the usual apical distribution, which is in line with previous reports suggesting pancreatic enzymes are released into interstitial space via this pathway. ^21^

qPCR validation of transcriptional alterations in ACP-induced *KC* mice pancreata confirmed the reprogramming of acinar cells toward ductal metaplasia (Figure 2*I*). Advanced-grade histological lesions progressing to pathologically evident cancer in *KC* mice pancreata with ACP induction correlated with a significantly shortened tumor latency period (Figure 2*J*, mean=5.80 months) compared with control (mean=9.76 months), A (mean=8.67 months), or CP (mean=7.56 months)-exposed *KC* mice, confirming a higher tumor burden induced by ACP than damage from A or CP induction alone.

### CREB is Persistently Activated During PanIN Progression in Kras^G12D^ Mutant Mice and in Human CP

Histological assessment using co-I.F analysis of pancreatic tissues harvested after ACP induction in *KC* mice substantially increased pCREB-positive staining within CK19^+^ ducts during the recovery periods (3 and 21 days) compared with the control pancreata (Figure 3*A*). Western blotting (Figure 3*B*) confirmed heightened and sustained CREB activation in ACP-induced *KC* mice at 3-, 21-, and 42-days post-ACP recovery, suggesting a positive feed-forward loop in *Kras\** in driving CREB expression.

**Figure 3.**
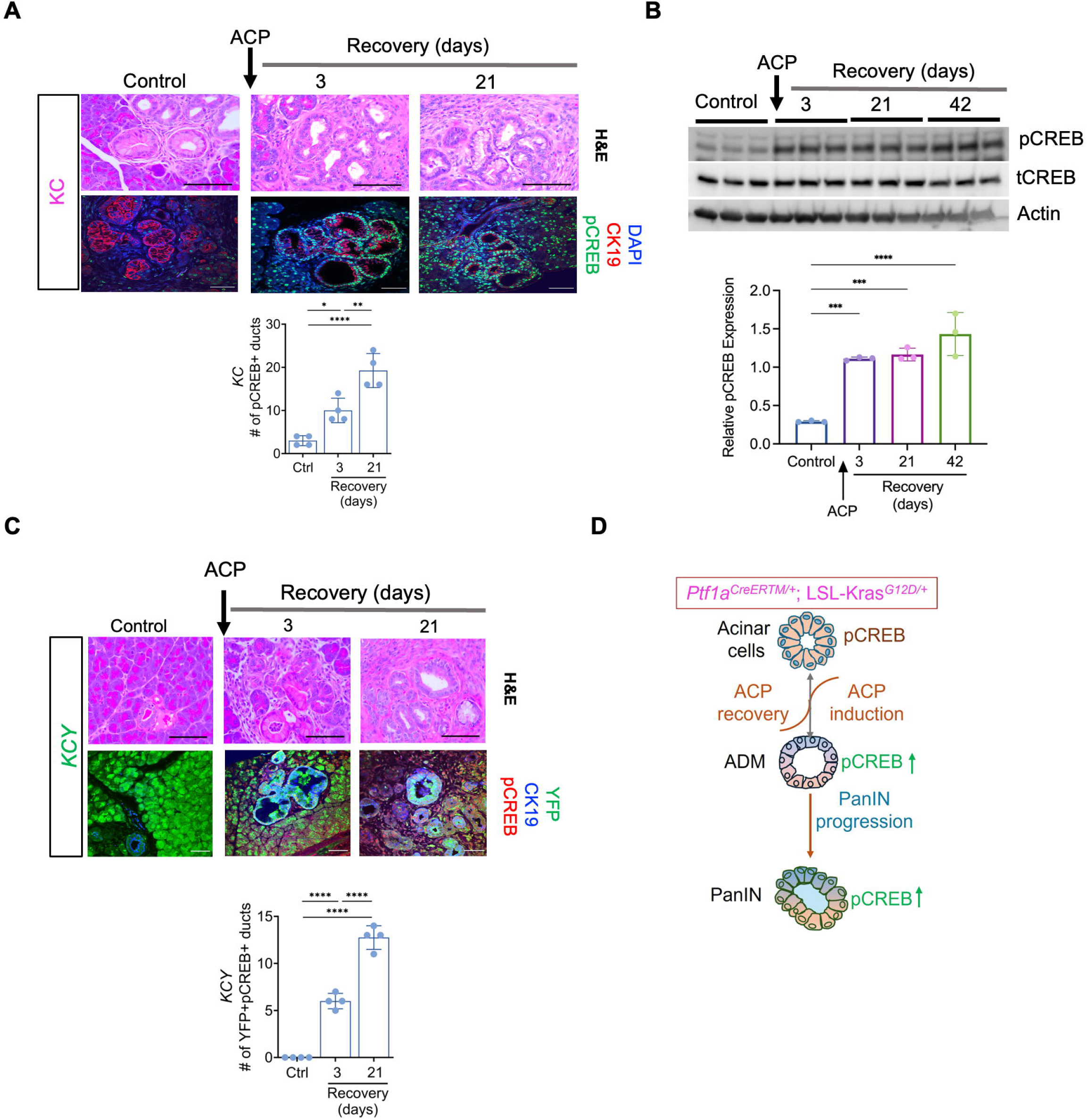
Sustained hyperactivation of CREB in ACP *Kras^G12D/+^* driven ADM and PanIN progression. (A) H&E and co-I.F labeling of pCREB (green) and CK-19 (red) expression (*top*) with pCREB^+^ duct quantification within the ctrl and *KC* mice pancreata at 3- and 21-day ACP recovery period (*bottom*) (n=4 mice per group). (B) Western blotting of pCREB in pancreatic tissue lysates of control and *KC* mice at 3-, 21- and 42-day ACP recovery period (n=3 mice per group). (C) H&E images of the pancreas along with co-I.F labeling of pCREB (red)/CK19 (blue)/YFP (green), and YFP^+^ pCREB^+^ duct quantification in ctrl and *Ptf1a^CreERTM/+^*; *LSL-Kras*^G12D/+^; *R26R-EYFP* (*KCY*) mice at 3- and 21-day recovery periods (n=4 mice per group). (D) Schematic depicting sustained hyperactivation of CREB driving acinar cell neoplastic reprogramming in *KC* mice upon ACP induction. Scale bar, 50 μm. *p<0.05; **p<0.01; ***, p<0.001; ****p<0.0001 by ANOVA for three groups comparison and unpaired T-test for two groups.

We combined the acinar-targeted *KC* GEMM with *R26R-EYFP* to generate the *Ptf1a-^CreERTM/+^;LSL-Kras^G12D/+^;R26R-EYFP (KCY)* mouse model for lineage tracing analysis (Figure 3*C*). The *KCY* mice underwent ACP induction (described in Figure 1*A*). Tamoxifen administration to *KCY* mice achieved approximately 95% recombination efficiency in acinar cells for 24 h.^22^ In control *KCY* mice, YFP expression was mainly in acinar cells and excluded from CK19^+^ ducts. ACP-induced pancreatic injury in *KCY* mice accelerated PanIN progression (similar to *KC*-ACP), with many ductal structures originating from acinar cells (YFP^+^CK19^+^ducts) (Figure 3*C*). We observed sustained CREB activation within acinar-derived YFP/CK-19 dual-positive ductal lesions at 3- and 21-day recovery periods post-ACP induction. These results suggest CREB’s role in reprogramming acinar cells into preneoplastic precursors with ACP and *Kras\** (Figure 3*D*).

### Acinar-specific Ablation of CREB Attenuates Spontaneous Kras^G12D^-mediated PanIN Progression

We generated acinar-specific conditional *Creb* knockout mice with *Kras\** (Figure 4*A*) (*Ptf1a^CreERTM/+^; LSL-Kras^G12D/+^*^;^ *Creb^fl/fl^* or *KCC^-/-^*), whereas *KC* mice with wild-type *Creb* were called *KC*. The pancreata were collected from these two mouse cohorts at 5 and 10 months of age. The relative pancreatic weight of *KC* mice with *Creb* deletion (*KCC^-/-^*) markedly reduced compared with that of wild-type *KC* mice (Figure 4*B* and *C*). Histological examination showed that the pancreata of 5-month-old *KC* mice displayed a higher prevalence of ADM and low-grade PanIN lesions (Figure 4*D* and *E*), whereas that of 5-month-old *KCC^-/-^* mice showed no precursor lesions and demonstrated a normal architecture. Pancreata harvested from 10-month-old *KCC^-/-^*mice displayed infrequent ADM or PanIN1 lesions. Conversely, 10-month-old *KC* mice pancreata exhibited marked high-grade PanIN lesions and cancer (Figure 4*D* and *E*).

**Figure 4.**
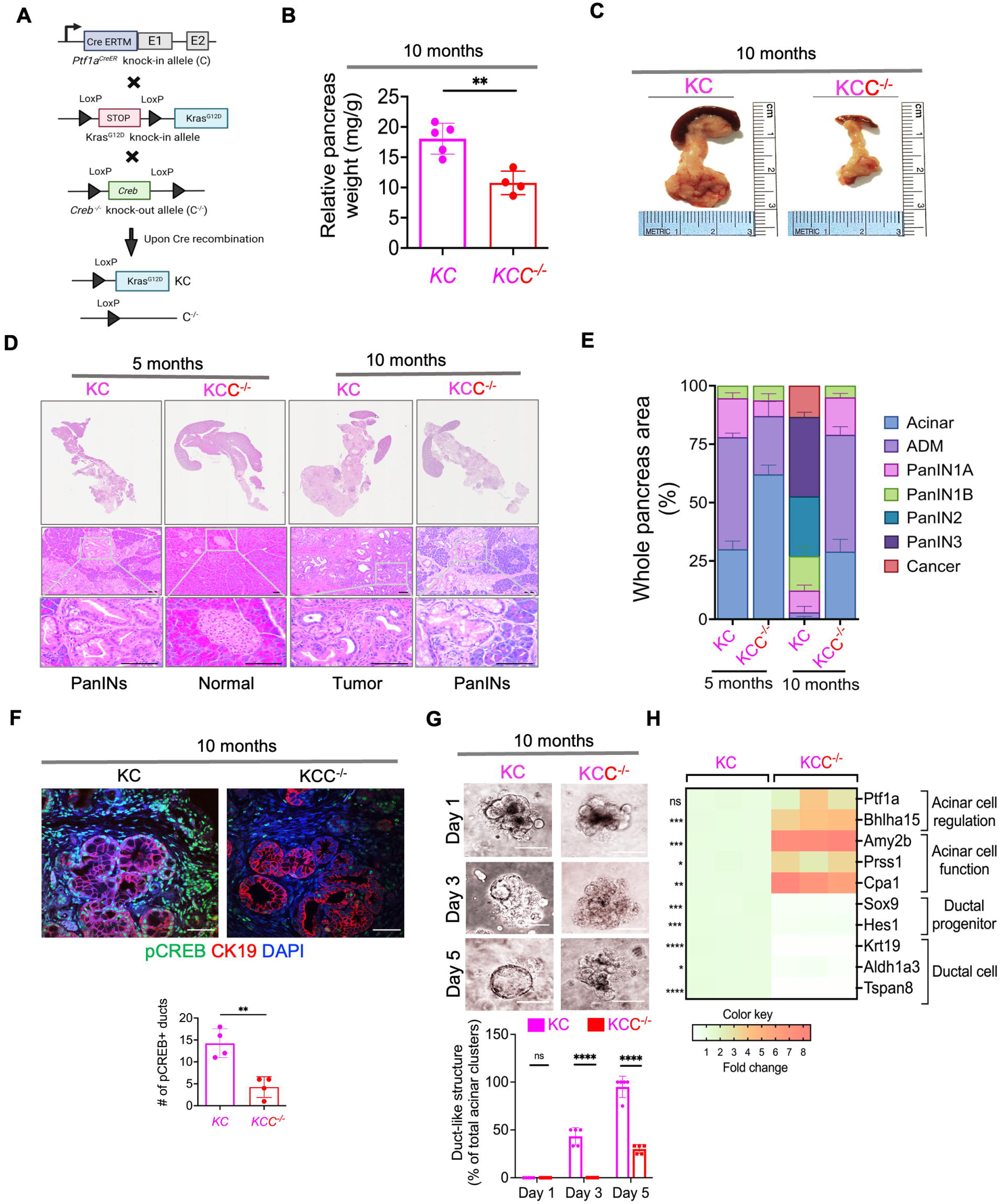
Acinar-specific ablation of *Creb* attenuates spontaneous *Kras^G12D/+^* mediated PanIN progression. (A) Breeding strategy for generating acinar-specific *Creb* deficient *Kras\** mutant mice (*Ptf1a^CreERTM/+^; LSL-Kras^G12D/+^*; *Creb^fl/fl^*or *KC*C^-/-^). (B) Comparative measurement of relative pancreas weight in 10-month-old *Creb* wild type (*KC*) and *KC*C^-/-^ mice (n=4–5 mice per group). (C) Representative photomicrographs of whole pancreas depicting significantly less tumor burden in *KC*C^-/-^ mice than *KC* mice at 10 months of age. (D) Representative H&E images of the pancreata harvested from 5- and 10-month-old *KC* and *KCC^-/-^* mice. Scale bar, 10 and 50 μm. (E) Comparative assessment using H&E-based histology to examine the entire pancreas, illustrating acinar cells, ADMs, PanINs, and cancerous regions in 5- and 10-month-old *KC* and *KCC^-/-^* mice (n=3 mice per group). (F) Co-IF analysis of pCREB (green), CK-19 (red), and DAPI (blue) with quantification in pancreatic tissue sections harvested from 10-month-old *KC* and *KCC^-/-^*mice (n=4 mice per group). Scale bar, 50 μm. (G) Representative bright-field photomicrographs of primary 3D acinar cell cultures established from 10-month-old *KC* and *KCC^-/-^* mice (scale bar, 20 µm) (n=5 mice per group). (H) qPCR of pancreatic tissue harvested from *KC* and *KCC^-/-^* mice (n=3 mice per group). Scale bar, 10 and 50 μm. ^ns^=nonsignificant; *p<0.05; **p<0.01; ***p<0.001; ****p<0.0001 one way ANOVA or unpaired t-test.

Co-I.F analysis in *KC* and *KCC^-/-^* mice (10 months old) confirmed a significant reduction in pCREB expression in the pancreas ductal regions in *KCC^-/-^* mice (Figure 4*F*). Pancreatic acinar cells were isolated from *KC* and *KCC^-/-^* mouse pancreata for explant culture. Acinar-specific *Creb* ablation significantly blocked ductal-like structure formation compared with acinar cells with intact CREB (Figure 4*G*). qPCR analysis of 10-month-old *KCC^-/-^* mice pancreata revealed reduced ductal cell phenotype genes, with a concomitant upregulation in genes linked to acinar cell regulation, function, proliferation, and differentiation compared with wild-type *KC* (Figure 4*H*). Taken together, these results suggest that CREB ablation reduces acinar cells ability to undergo ADM reprogramming.

### Acinar-specific Ablation of Creb Attenuates Kras*-mediated PanINs to Pancreatic Cancer Progression with ACP

In *Ptf1a^CreERTM/+^* mice with *Creb* deletion (CC^-/-^) without *Kras\** (Supplementary Figure 4A), histological assessment confirmed the loss of CREB expression within *Ptf1a^CreERTM/+^;Creb*^fl/fl^ (CC^-/-^) mice pancreata compared with wild-type *Ptf1a^CreERTM/+^*(C) mice (Supplementary Figure 4B). *CC^-/-^* mice exhibited enhanced acinar cell regenerative potential post-ACP induction (Supplementary Figure 4C). The improved recovery in *Creb*-deleted mice was associated with reduced CK19^+^ ducts and attenuated fibroinflammatory milieu.

To assess acinar-specific *Creb* ablation’s capacity to counteract *Kras\****-**driven ADM/PanIN originating from the acini, we histologically compared *KC* and *KCC^-/-^* mouse pancreata (both at 3- and 21-day recovery) (Figure 5*A*). *KC* and *KCC^-/-^* mice maintained on a standard liquid diet were used as controls. ACP induction in *KC* mice considerably increased the relative pancreatic weight (21-day recovery period), confirming a higher tumor burden than that in the control *KC* mice (Supplementary Figure 5A). This was less prominent in *KCC^-/-^* mice with ACP induction, where CREB was deleted (Supplementary Figure 5A and B), suggesting an association between attenuated tumor burden and CREB loss.

**Figure 5.**
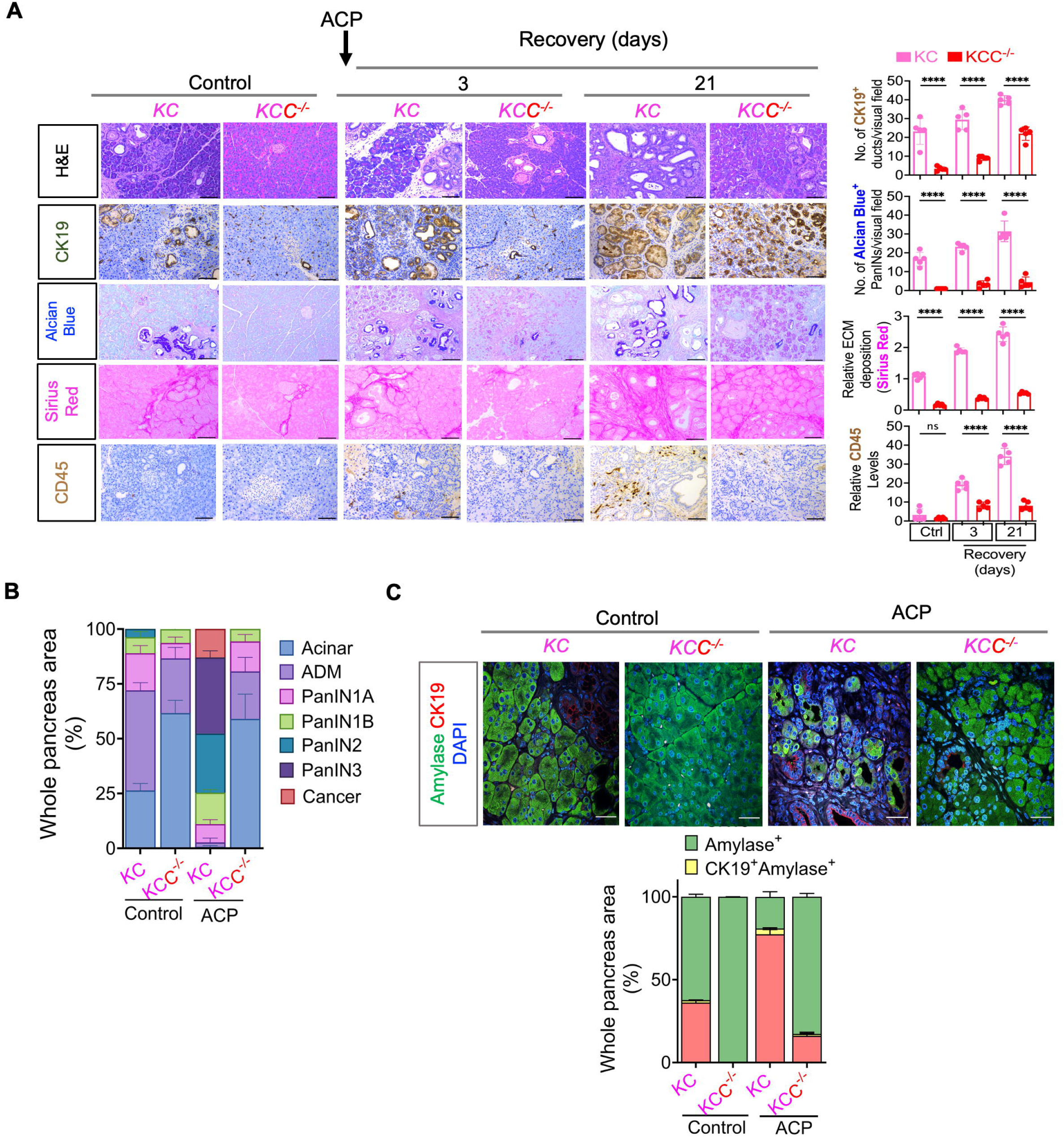
Acinar-specific ablation of *Creb* attenuates *Kras\** induced progression of ADM/PanINs toward pancreatic cancer with ACP. (A) Comparative histological evaluation of mouse pancreas, accompanied by images showcasing H&E, CK-19, Alcian Blue, Sirius Red, and CD45^+^ staining within *KC* and *KCC^-/-^* mice pancreata, ctrl and *KC* mice at 3- and 21-day ACP recovery period (n=5 mice per group). (B) H&E-based histological examination illustrating acinar cells, ADMs, PanINs, and cancerous regions in ctrl or within *KC* and *KCC^-/-^* mice pancreata harvested after 21 days of ACP recovery period (n=3 mice per group). (C) Pancreas images depicting amylase (green)/CK19 (red) and DAPI (blue) immunofluorescent labeling (*top*) and CK19^+^ amylase^+^ cell corresponding quantification (*bottom*)[within the epithelial cell compartment only] in ctrl (*KC* and *KCC^-/-^*) or with ACP induction (n=3 mice per group). Scale bar, 50 μm. ^ns^=nonsignificant; ****p<0.0001 by ANOVA.

*Creb* deletion in *KCC^-/-^* mice pancreata remarkably protected against ACP-triggered ADM and PanIN progression as evidenced by H&E, CK19, and Alcian blue staining (Figure 5*A* and *B*). Post-ACP induction, *KC* mice pancreata displayed a higher prevalence of high-grade PanIN lesions along with pancreatic cancer. Conversely, pancreatic tissue architecture in *KCC^-/-^* mice displayed mostly normal organization of acinar cells, with fewer ADMs and low-grade PanIN lesions (Figure 5*A* and *B*). A notable reduction in the fibroinflammatory milieu (Sirius red and CD45^+^ cells) was observed (3- and 21-day ACP recovery periods) compared with that in *KC* GEMM. Similar beneficial histological trends occurred in control *KCC^-/-^*mice. Co-I.F staining within *KC* and *KCC^-/-^* mice pancreatic tissues showed a notably heightened proportion of amylase-positive acinar cells in *Creb-*depleted KCC^-/-^ mice (Figure 5*C*). Conversely, *KC* mice pancreata exhibited substantial damage resulting from ACP induction, with high CK-19^+^ cell abundance and loss of amylase^+^ acini.

### Acinar-specific Ablation of CREB Attenuates ACP-Induced Acinar Cells to Ductal Reprogramming and Increases Tumor Latency Period

We studied ACP-induced pancreatic injury in a lineage tracing mouse model of *KC (KCY; Ptf1a-^CreERTM/+^;LSL-Kras^G12D/+^;R26R-EYFP)* and *KCYC^-/-^* (*Creb* deletion in *KCY)* mice (Figure 6*A*). Co-I.F staining displayed more YFP^+^/CK19^+^ ductal lesions concurrent with the loss of the acinar cell compartment upon ACP induction in *KCY* mice. We found a substantial reduction in ductal structures, with minimal cells exhibiting dual positivity of YFP^+^/CK19^+^ during ACP-induced *KCYC^-/-^*. qPCR analysis confirmed a notable decrease in the mRNA expression levels of genes associated with ductal cell identity and function (*Sox9*, *Krt19*, *Aldh1a3*, and *Tspan8*) in ACP-treated *KCC^-/-^* mice pancreata compared with *KC* mice (Figure 6*B*).

**Figure 6.**
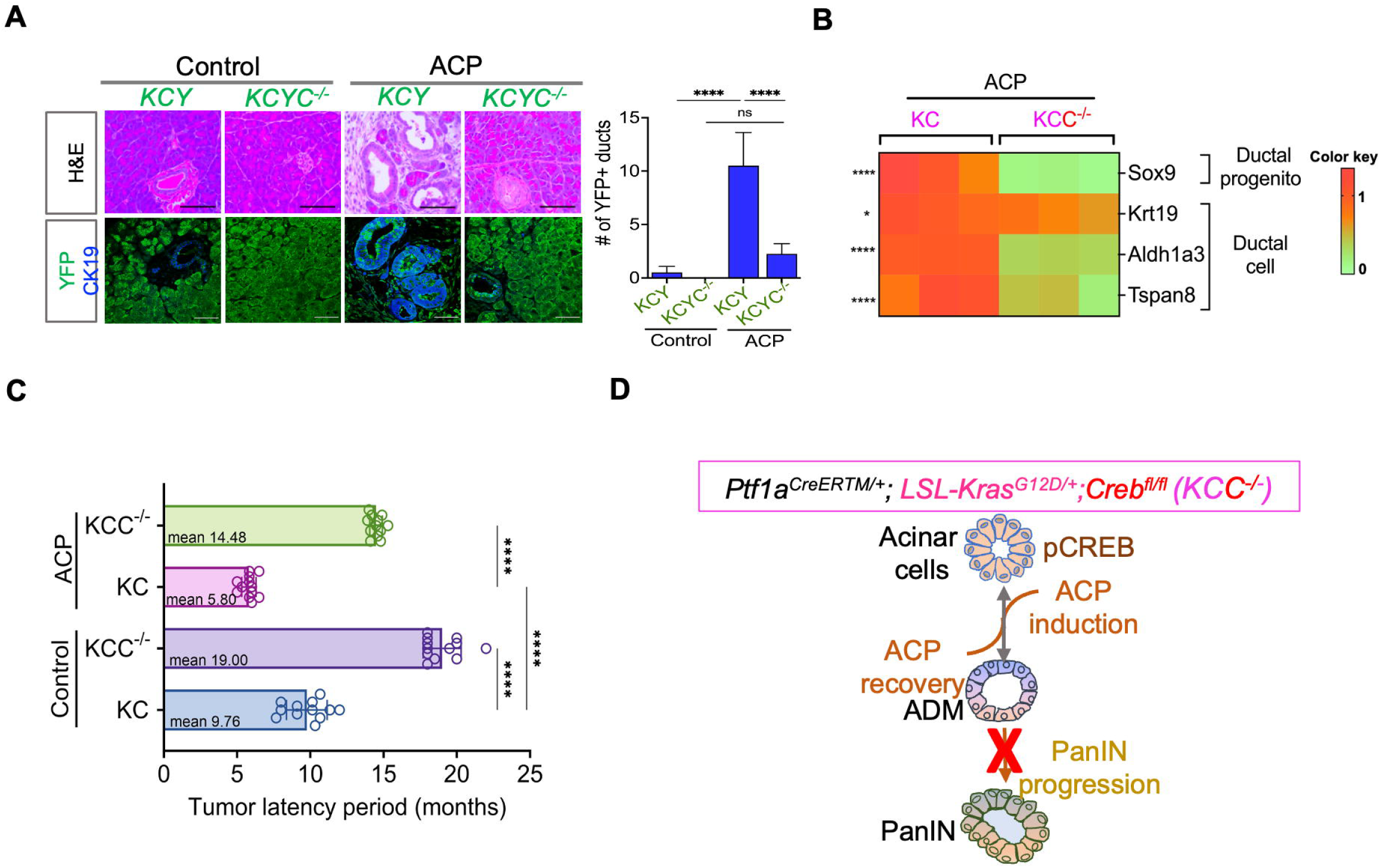
Acinar-specific ablation of *Creb* attenuates ADM reprogramming and increases tumor latency period with ACP induction in *KC* mice. (A) Representative H&E images of the pancreas along with co-immunofluorescent labeling and YFP quantification (green)/CK19 (blue) in *Ptf1a-^CreERTM/+^;LSL-Kras^G12D/+^;R26R-EYFP (KCY)* and *Creb*^fl/fl^ (*KCYC^-/-^*) in control or with ACP induction (n=4 mice per group). (B) qPCR-based analysis of *KC* and *KCC^-/-^* mice pancreatic tissue with ACP induction (n=3 mice per group). (C) Tumor latency period assessment in ctrl or with ACP induction in *KC* and *KCC^-/-^* mice (n=12 mice per group, ctrl mice cohort represented here is similar to that shown in Figure 2I). (D) Schematic representation illustrating that acinar-specific *Creb* ablation diminishes pancreatic cancer progression in ACP induction in *KC* mice. Scale bar, 50 μm. ^ns^=nonsignificant;*p<0.05; ****p<0.0001 by ANOVA or unpaired t-test.

*Creb*-deficient GEMMs (*KCC^-/-^* and *KCYC^-/-^)* displayed low-grade PanIN lesions and a diminished pancreatic ductal phenotype, further reflected in tumor latency (Figure 6*C*). ACP-induced *KCC^-/-^* mice pancreata had a mean pathological malignancy duration of 14.48 months compared with 5.80 months for *KC*-ACP mice. Conversely, control *KC* harboring wild-type *Creb* and *KCC^-/-^* exhibited greater tumor latency (9.76 vs. 19 months) (Figure 6*C*). Overall, these results suggest that acinar-specific *Creb* ablation reduces ACP-induced acinar-to-ductal reprogramming and *Kras\**-induced neoplastic progression, delaying tumor burden (Figure 6*D*).

## Discussion

A consensus exist that alcohol-dependent mouse models replicate the clinical presentation of human CP. ^23, 24^ Chronic alcohol exposure induces membrane instability within the zymogen and lysosomal compartments of acinar cells, increasing the risk of premature enzyme activation and autodigestive injury.^25, 26^ When alcohol exposed mice are subjected to additional stressors such as supraphysiological doses of caerulein, this priming effect leads to an additive and exacerbated pancreatic injury, profoundly damaging the pancreas and affecting other cellular constituents as well (including PSCs and ductal cells). ^27^ Based on these prior findings in animal models-irrespective of mouse sex, ^28^ we demonstrate that GEMMs of pancreatic disease (with or without *Kras\**) recapitulate the clinical progression of chronic pancreatic injury through a combination of ethanol-based diet and caerulein administration, mimicking the pathological features of human ACP. ^18, 29–34^

Pancreatic cancer or PDAC, is characterized by widespread genomic alterations and extensive tumor cell heterogeneity.^35–37^ Previous studies in mouse models have highlighted that both acinar and ductal cells can result in PDAC, with acinar cells more amenable to cellular plasticity and transform into a progenitor-like cell type with ductal characteristics (ADMs) with *Kras\**.^6, 8, 11, 38^ In our current study, utilizing multiple GEMMs and lineage tracing studies, we demonstrated that under chronic inflammatory stimulation by ACP, leads to sustained activation of CREB in ADMs undergoing ductal transition, which cooperates with *Kras\** to accelerate pancreatic cancer progression (Figure 7).

**Figure 7.**
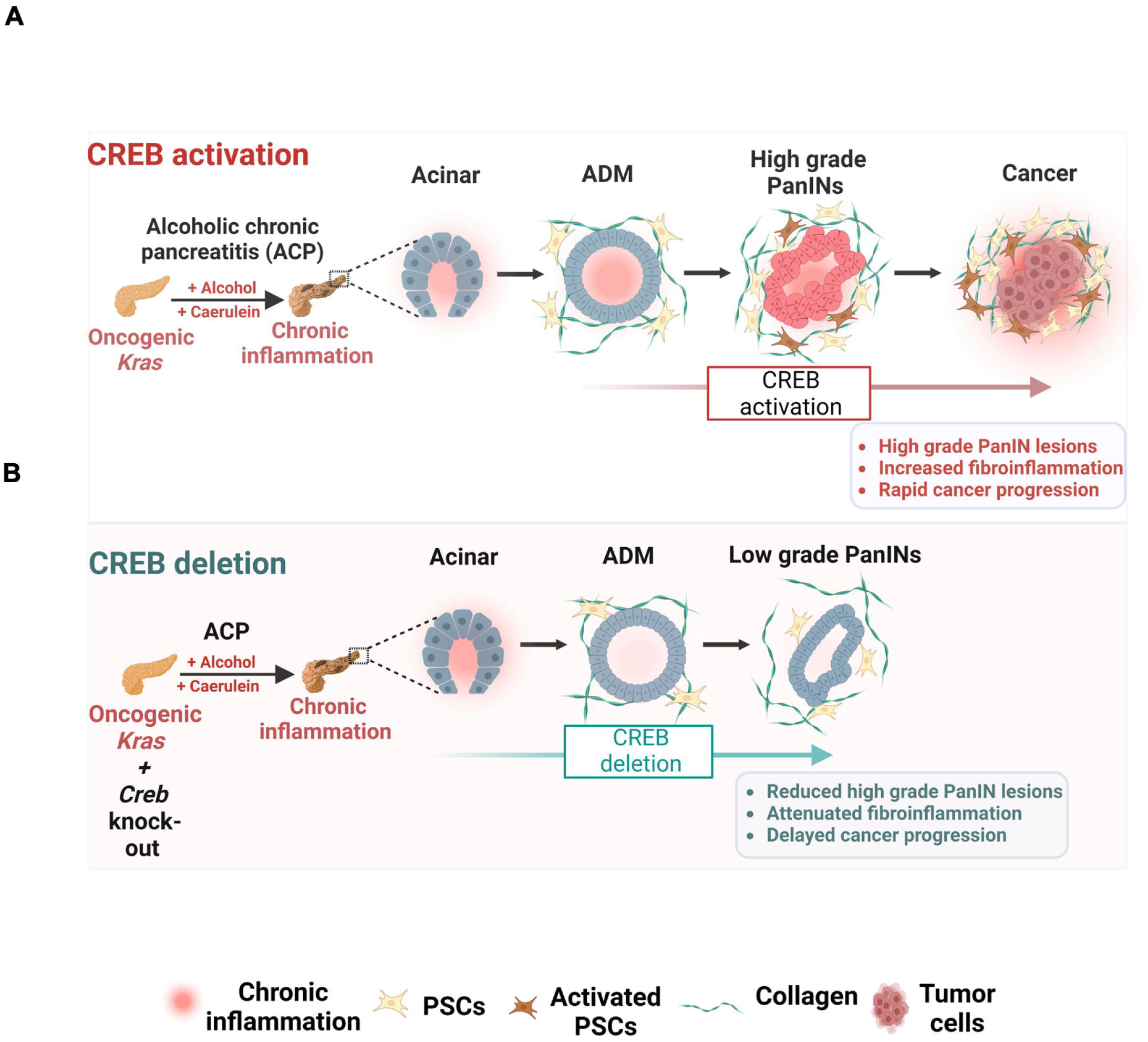
(A) Schematic representation of ACP induction with *Kras\** driving irreversible ADM reprogramming toward accelerated PDAC progression along with sustained CREB activation and increased fibroinflammation. (B) Acinar cell-specific deletion of CREB generates anti-tumor responses and delays pancreatic cancer progression. The schematic was created with BioRender with License agreement numbers OW27SH91EN.

Crosstalk between epithelial cells and inflammation-driven stress response mediators promotes the aggressive biology of pancreatic cancer. We have previously discovered that GM-CSF mediated activation of CREB contributes to PDAC growth and therapeutic resistance within the context of smoking-associated chronic inflammation. ^15^ Simultaneously, other studies have also reported that PDAC epithelial cells sustain their proliferation through obesity-associated stress responses, activating CREB at Ser133 via protein kinase dependent signaling.^15, 39^ Thus, it is plausible, that the additive stimulus of alcohol and caerulein exacerbates epithelial cell intrinsic and extrinsic stress response mediators, converging on CREB, as a central hub of signal integration to accelerate pancreatic cancer progression in GEMM models. This is supported by our scRNA-seq analysis of *KC-*ACP GEMM which revealed acinar cell populations enriched in pathways prominently related to cellular responses to stress and reactive oxygen species (ROS). Simultaneously, we observed a profound accumulation of innate immunosuppressive myeloid subsets–macrophages and granulocytes within the pancreata of ACP-exposed *KC* mice. Our ongoing research is focused on elucidating these complexities of epithelial-myeloid crosstalk, and specifically how it drives CREB activation not only in ductal cells but also across diverse cellular compartments through autocrine and paracrine signaling within the tumor microenvironment.

Recent studies from our group and others have firmly established CREB as a transcription activator and a key cooperative node between *KRAS\**-mediated effector pathways and other genomic drivers, including *TP53*, in pancreatic cancer.^40, 41^ Upregulation of CREB expression has been associated with increased proliferation and migration of pancreatic cancer cells, evasion of apoptosis, and induction of epithelial to mesenchymal transition (EMT).^42–44^ Pharmacological targeting of CREB activation using small molecule inhibitor has been shown to dampen PDAC progression and metastatic spread. ^15, 41, 45^ However, pancreas-specific CREB deleted GEMMs were lacking, which limited our understanding of its oncogenic potential. Therefore, building on a growing body of evidence implicating CREB and its downstream signaling in pancreatic cancer progression, for the first time, our study extensively characterized CREB-deleted GEMMs with *Kras*.* Acinar-specific CREB loss significantly attenuated both spontaneous and ACP-mediated ADM/PanIN formation while simultaneously promoting acinar cells regeneration in *KC* GEMM. Supporting this notion, CREB deletion further enhanced the transcriptional rewiring of gene hubs that promote acinar cell regenerative potential and function.

Pancreatic cancer cell heterogeneity is mainly governed by exocrine compartment plasticity. Acinar cells transition to a duct-like identity in response to exogenous stress, including ACP, making them susceptible to *Kras\**-induced precancerous lesions. Given that multiple upstream signaling pathways can activate CREB, the current study focuses only on defining CREB’s role in the ductal epithelial compartment, both in the presence and absence of ACP. However, comparing the differential regulation of CREB expression in alcohol or CP settings alone with *Kras\** remains an important area for future investigation.

Several upstream kinases phosphorylate CREB at Ser133, which triggers the recruitment of the coactivator CBP. ^46^ This complex then binds to target gene promoters’ cAMP-response element (CRE) sequences and facilitates gene regulation via intrinsic and associated acetylase activities and/or interacting with core transcriptional machinery. ^47^ Previous studies have outlined the cancer cell-autonomous and extrinsic effect of CREB activation promoting PDAC aggressiveness and neutrophil-mediated immunosuppression by engaging with downstream effectors, including (FOXA1, β-catenin, and CXCL1). ^40, 41^ Our ongoing studies are focused on uncovering novel downstream targets by which CREB regulates ADM-PanIN progression with chronic inflammation and *Kras\**. In future studies, we aim to evaluate whether genetic or pharmacological inhibition of these targets can phenocopy CREB loss and suppress pancreatic cancer progression.

In conclusion, ACP induction promotes pancreatic cancer progression via irreversible ADM reprogramming with *Kras\** cooperativity. These ductal lesions originating from acinar cells display continuous CREB hyperactivation. Attenuating high-grade PanIN formation and tumor progression through acinar-specific loss of *Creb* provide foundational insights for future studies on CREB’s role in pancreatic cancer progression in alcoholic pancreatitis with *Kras\**.

## Materials and Methods

### Creb Deletion in a Genetically Engineered Ptf1a^CreERTM/+^;LSL-Kras^G12D/+^ (KC) Mice Model

*Ptf1a^CreERTM/+^* knock-in allele mice were obtained from Jackson Laboratory (Bar Harbor, ME; stock number: 019378)^22^ and crossed with *LSL-Kras*^G12D/+^ to generate acinar-specific *Kras^G12D/+^* mutant mice, referred to as *Ptf1a^CreERTM/+^;LSL-Kras^G12D/+^*(*KC*) mice. To generate acinar-specific *Creb1* (*Creb*) knockout mice, *Ptf1a^CreERTM/+^; Creb^fl/fl^* mice were crossed with *LSL-Kras^G12D/+^; Creb^fl/fl^* mice to generate *Ptf1a^CreERTM/+^;LSL-Kras^G12D/+^; Creb^fl/fl^* (KCC^-/-^) mice. The *Creb^fl/fl^*mice were obtained from Professor Eric Nestler (Cold Spring Harbor Laboratory, Cold Spring Harbor, NY 11724).^48^ *Creb* was selectively excised from acinar cells in *LSL-Kras^G12D/+^* mice using inducible Cre recombinase, expressed from the pancreas-specific *Ptf1a^CreERTM/+^*promoter. Cre recombinase activity was induced in 6-week-old mice through daily tamoxifen administration (Sigma-Aldrich, cat. # T5648) for 6 consecutive days at a dose of 0.15 mg/g body weight, followed by a 1-week rest before starting the ACP induction phase.

### Lineage Tracing in R26R^EYFP^ Reporter Mice

*R26R^EYFP^* mice (obtained from Jackson Laboratory; stock number: 006148) ^49^ were used for lineage tracing. To precisely trace the lineage of recombined acinar cells, *Ptf1a^CreERTM/+^* mice containing the *R26R^EYFP^*reporter were crossed with *LSL-Kras^G12D/+^* mice to generate *Ptf1a^CreERTM/+^;LSL-Kras^G12D/+^; R26R^EYFP^*. These mice were subsequently bred with *Creb^fl/fl^* mice to produce *Ptf1a^CreERTM/+^;LSL-Kras^G12D/+^; R26R^EYFP^; Creb^fl/fl^* (hereafter referred to as *KCYC^-/-^*) mice.

### Mouse Genotyping

Genotyping was conducted using the automated genotyping service provider, Transnetyx (Cordoba, TN, USA). Supplementary Table 1 presents the probe sequences used for genotyping analysis.

### Animal Studies

Here, mice of both sexes weighing 20–25 g were used; they were housed in pathogen-free conditions under a 12-h light-dark diurnal cycle with a controlled temperature of 21°C–23°C and maintained on a standard rodent chow diet (Harlan Laboratories) before the experimental induction of the ACP protocol. The mice were euthanized upon the manifestation of signs of compromised health, including weight loss, accelerated respiration, hunched posture, piloerection, and reduced activity. All animal experiments were approved and performed in compliance with the regulations and ethical guidelines for experimental and animal studies of the Institutional Animal Care and Use Committee and the University of Miami guidlines (Miami, FL; Protocol Nos. 15-057, 15-099, 18-081, and 21-093). All authors had access to the study data and had reviewed and approved the final manuscript.

### Establishing ACP In Vivo

The experimental mice were pair-fed with alcohol for 14 weeks using a Lieber-DeCarli alcohol-based liquid diet (A) (BioServ Inc., cat. #F1259SP) containing 5% v/v ethanol, whereas the control mice received a standard control liquid diet (C) (BioServ Inc., cat. #F1259SP) using 28% carbohydrates instead of ethanol. During the final 4 weeks of alcohol exposure, CP was induced by administering cerulein solubilized in phosphate-buffered saline to achieve a concentration of 10 mg/mL (ACP induction period). This feeding regimen mimics the pancreatic damage from chronic alcohol use in humans.^18, 23, 50^ Cerulein was intraperitoneally delivered at a dose of 50 μg/kg through hourly injections (six times daily, 3 days weekly) over 4 weeks. This combined effect of alcohol and cerulein was observed in the ACP group. The animals were euthanized humanely to harvest the blood and pancreas after ACP induction on days 3 and 21 (ACP recovery period). Supplementary Table 2 presents a comprehensive overview of the treatment groups spanning both the induction and recovery phases.

### Tumor Latency Period Estimation

In the context of genetically engineered murine models (GEMMs), the tumor latency period is the duration between tumor initiation or formation and the point at which these pancreatic tumors become grossly discernible during biweekly abdominal palpation examinations. Mice exhibiting noticeable signs of illness attributed to an increased tumor burden were euthanized. Subsequently, the pancreatic tissues were collected and subjected to hematoxylin and eosin (H&E) staining for pathological analysis.

### Pancreatic Digestion for Single-Cell RNA Sequencing (scRNA-seq) and Library Generation

The mouse pancreatic tissue harvested from control *KC* and *KC* with ACP induction were mechanically dissociated to generate single-cell suspensions, as previously described.^19^ Pancreatic fragments (1–2 mm) were immersed in 0.02% trypsin C-ethylenediamine tetra acetic acid 0.05% (Biological Industries) for 10 min at 37°C with agitation, then washed with 10% fetal calf serum/Dulbecco’s Modified Eagle Medium. For the next dissociation step, the cells were washed with Hanks’ balanced salt solution (HBSS) × 1 containing 1 mg/mL collagenase P, 0.2 mg/mL bovine serum albumin (BSA), and 0.1 mg/mL trypsin inhibitor. After 20–30 min incubation at 37°C with agitation, the samples were pipetted up and down, returned to 37°C, passed through a 70-mm nylon mesh (Corning #431751), and washed two times with 1×HBSS containing 4% BSA and 0.1 mg/mL DNase I. The samples were divided into two equal volumes, with one sample subjected to centrifugation and washed three times at 60 × *g* to isolate large cells containing acinar cells. Simultaneously, the second sample was centrifuged and washed three times at 300 × *g* to collect all cells. If red blood cells were abundant in the second sample, they were treated with red blood cell lysis buffer (Sigma-Aldrich). Live cells were isolated using a MACS Dead Cell Removal Kit (Miltenyi Biotech #130-090-101). Finally, the two samples were combined at a cell ratio of 30% and 70% from the 60 × *g* and 300 × *g* samples, respectively, and processed for scRNA-seq library preparation by the Oncogenomics-Shared Resource Facility (University of Miami). Briefly, the cells were counted, and 10,000 cells were loaded per lane on 10× chromium microfluidic chips. Single-cell capture, barcoding, and library preparation were performed using the Chromium 85 Controller, Chromium Next GEM Single Cell 3’ GEM, Library & Gel Bead Kit v3.1, and Chromium Next GEM Chip G kit (10× Genomics), with a cell recovery target of 100,000 cells as per manufacturer’s guidelines. The cDNA and libraries were sequenced using an Illumina NovaSeq 6000. The Cell Ranger pipeline (version 7.0, 10× Genomics) transformed Illumina base call files into FASTQ files, aligned them to the GRCm38 reference genome, and generated a digital gene-cell count matrix through the Biostatistics and Bioinformatics Shared Resource facility (University of Miami). Subsequently, the count matrices were imported into R version 3.5.0 and analyzed using the R package Seurat version 4.0.13-15.

For additional details on the experimental methodology, please refer to the Supplementary Materials and Methods section.

## Abbreviations

ACP: alcoholic chronic pancreatitis;
ADM: acinar-to-ductal metaplasia;
cAMP: response element binding protein 1 (CREB);
CP: chronic pancreatitis;
CK-19: cytokeratin 19,
CRE: cAMP response element;
EYFP: enhanced yellow fluorescent;
EMT: epithelial to mesenchymal transition;
GEMM: genetically engineered mouse model;
GM-CSF: granulocyte-macrophage colony stimulating factor;
I.F: Immunofluorescence;
KC: *Ptf1a^CreERTM/+^;LSL-Kras^G12D/+^*;
*Kras\**: *Kras ^G12D^*^/+^;
KCC^-/-^: *Ptf1a^CreERTM/+^;LSL-Kras^G12D/+^ Creb^floxed/floxed^*;
PanIN: pancreatic intraepithelial neoplasia;
PDAC: pancreatic ductal adenocarcinoma;
PSC: pancreatic stellate cells;
ROS: reactive oxygen species;
scRNAseq;: single-cell RNA sequencing;
UMAP: uniform manifold approximation and projection for dimension reduction,
αSMA: α smooth muscle actin

## Acknowledgements

The authors thank Dr. Erin Dickey for her assistance in the editing process and Dr. Oliver McDonald for his help in assessing the histopathology sections. Research reported in this publication was performed in part at the Analytical Imaging Shared Resource (AISR), Onco-Genomics Shared Resource (OGSR;RRID:SCR_022502), Cancer Modeling Shared Resource (CMSR;RRID:SCR_022891), and Biostatistics and Bioinformatics Shared Resource (BBSR;RRID:SCR_022890) of the Sylvester Comprehensive Cancer Center at the University of Miami Miller School of Medicine, and in part by the National Cancer Institute of the National Institute of Health under Award Number P30-CA240139. BioRender was used for the creation of Figure 4A (License agreement SC265R8O4S) and Supplementary Figure 4A (License agreement RS265R8Z28). The authors thank Editage (www.editage.com) for English language editing.

## Data Availability

Single cell RNA sequencing data are available at the Sequence Read Archive of NCBI under the BioProject ID: PRJNA109588

---

**Supplementary Figure 1.**
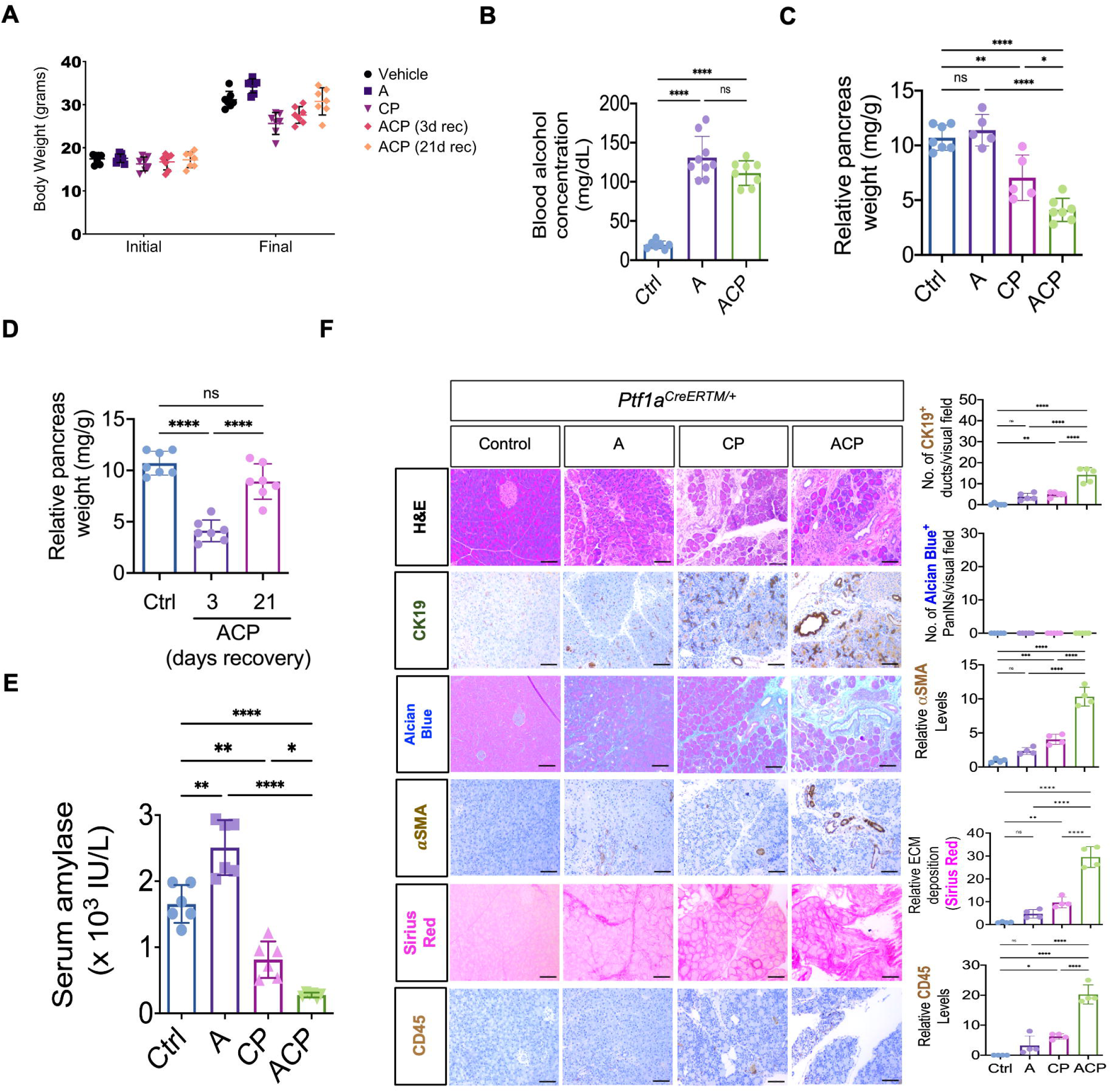

**Supplementary Figure 2.**
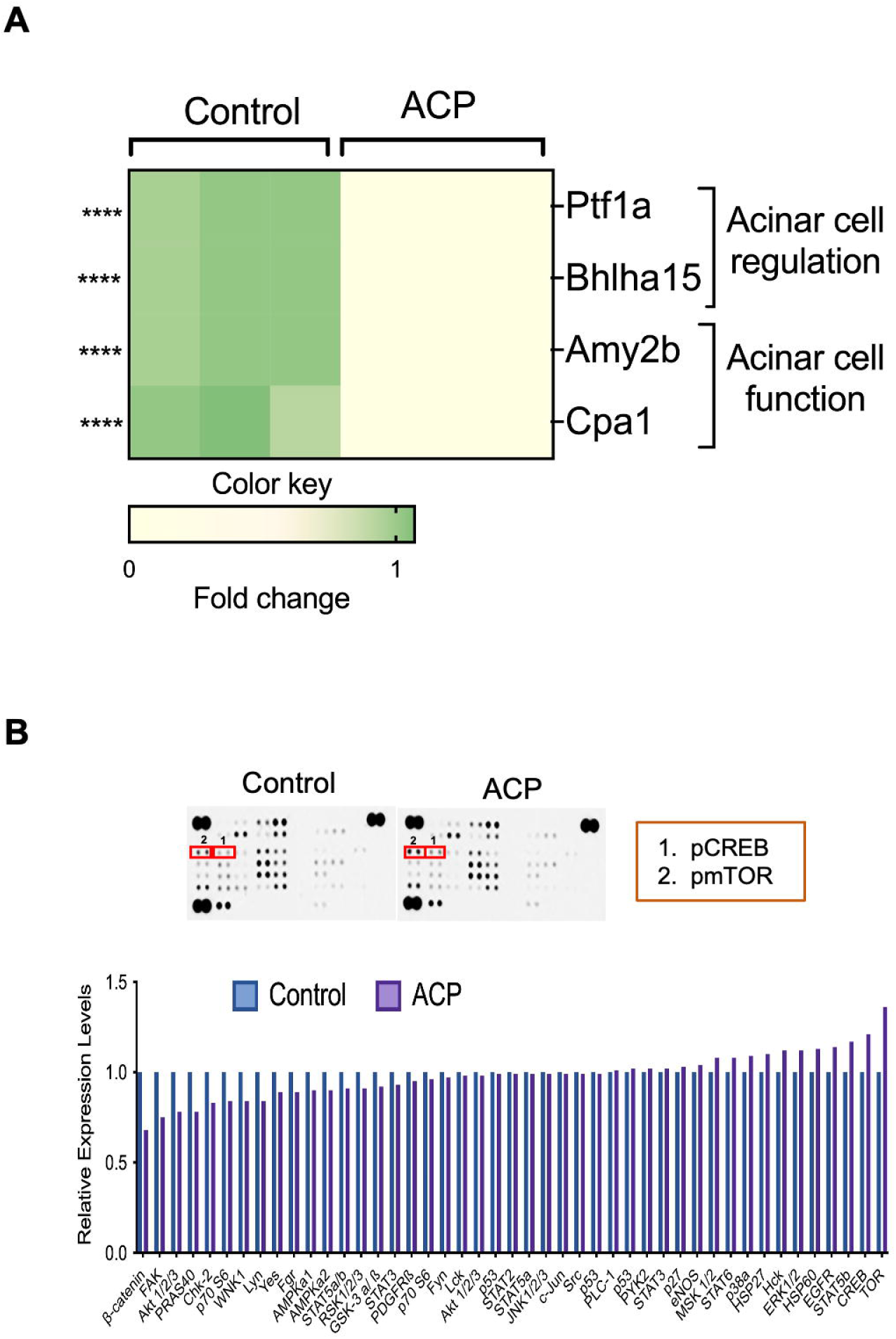

**Supplementary Figure 3.**
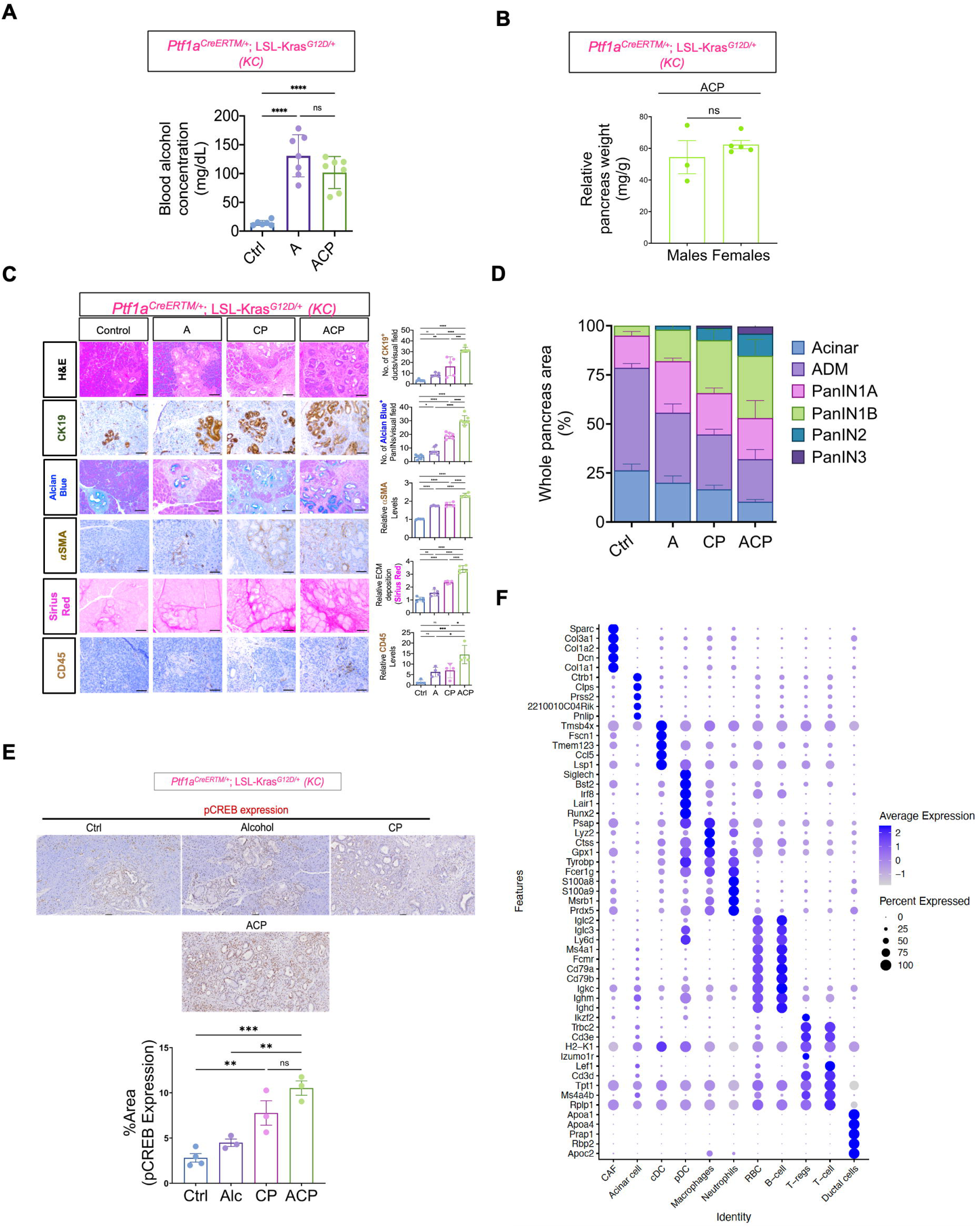

**Supplementary Figure 4.**
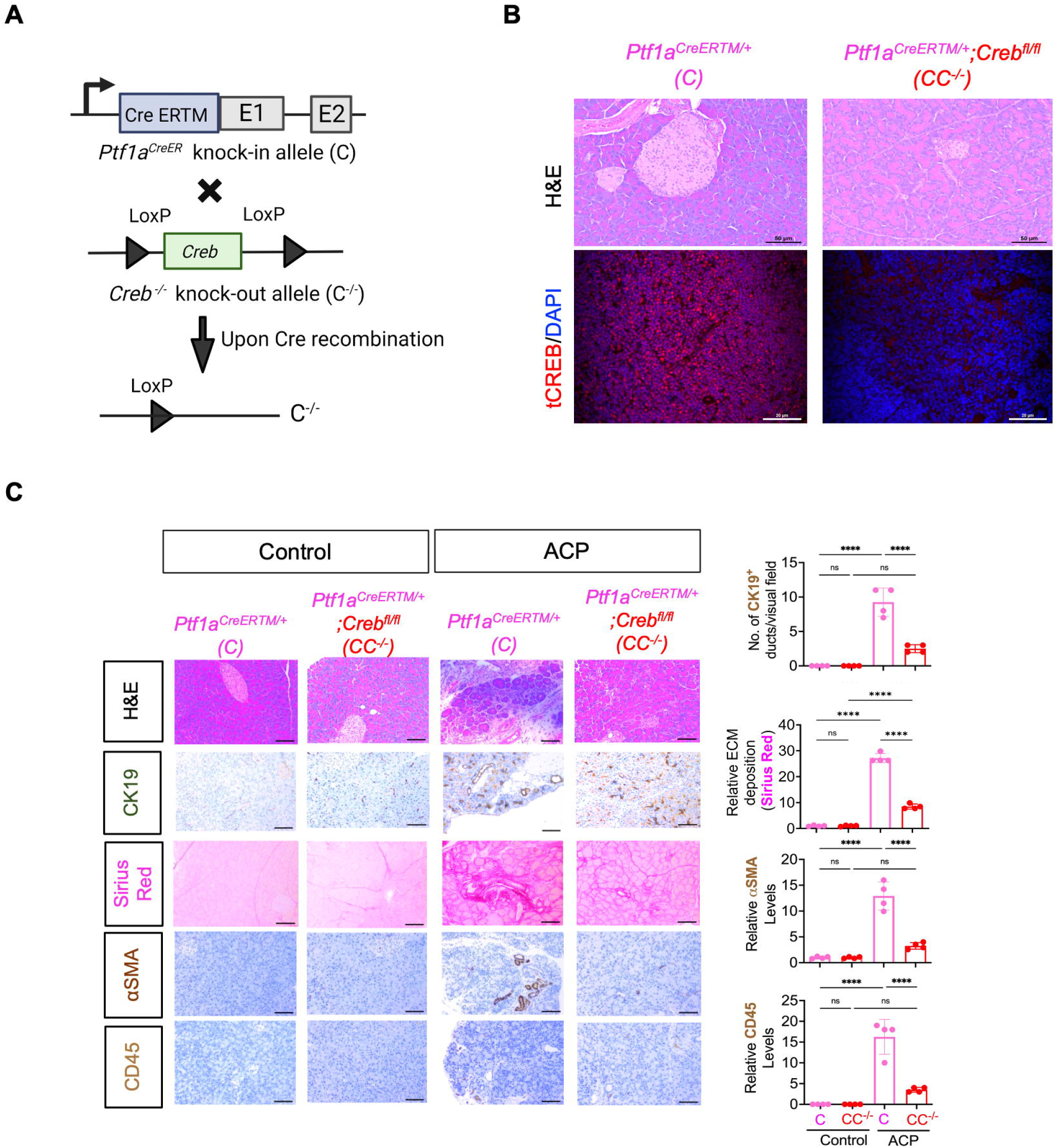

**Supplementary Figure 5.**
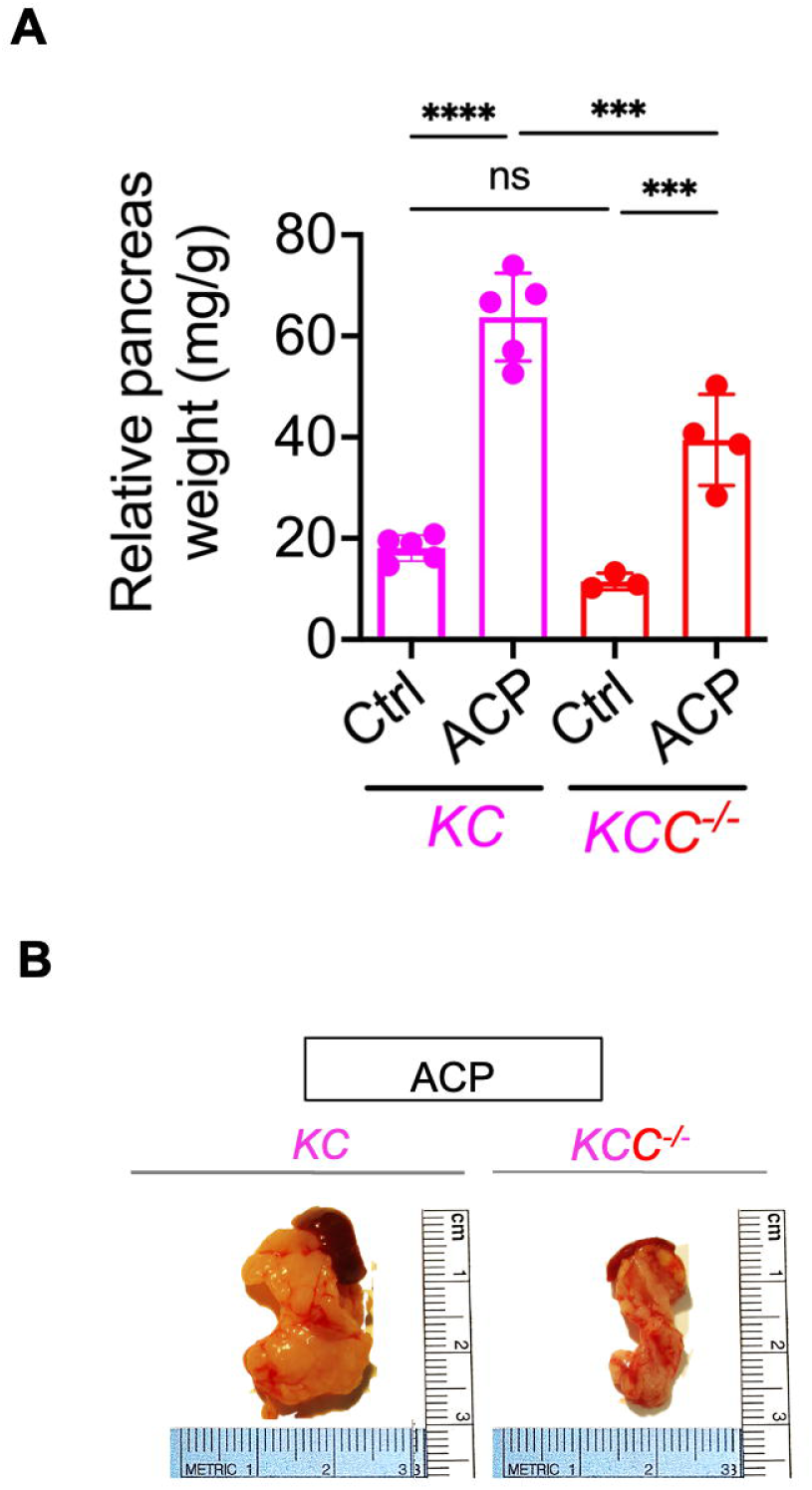

**Supplementary Figure 6.**
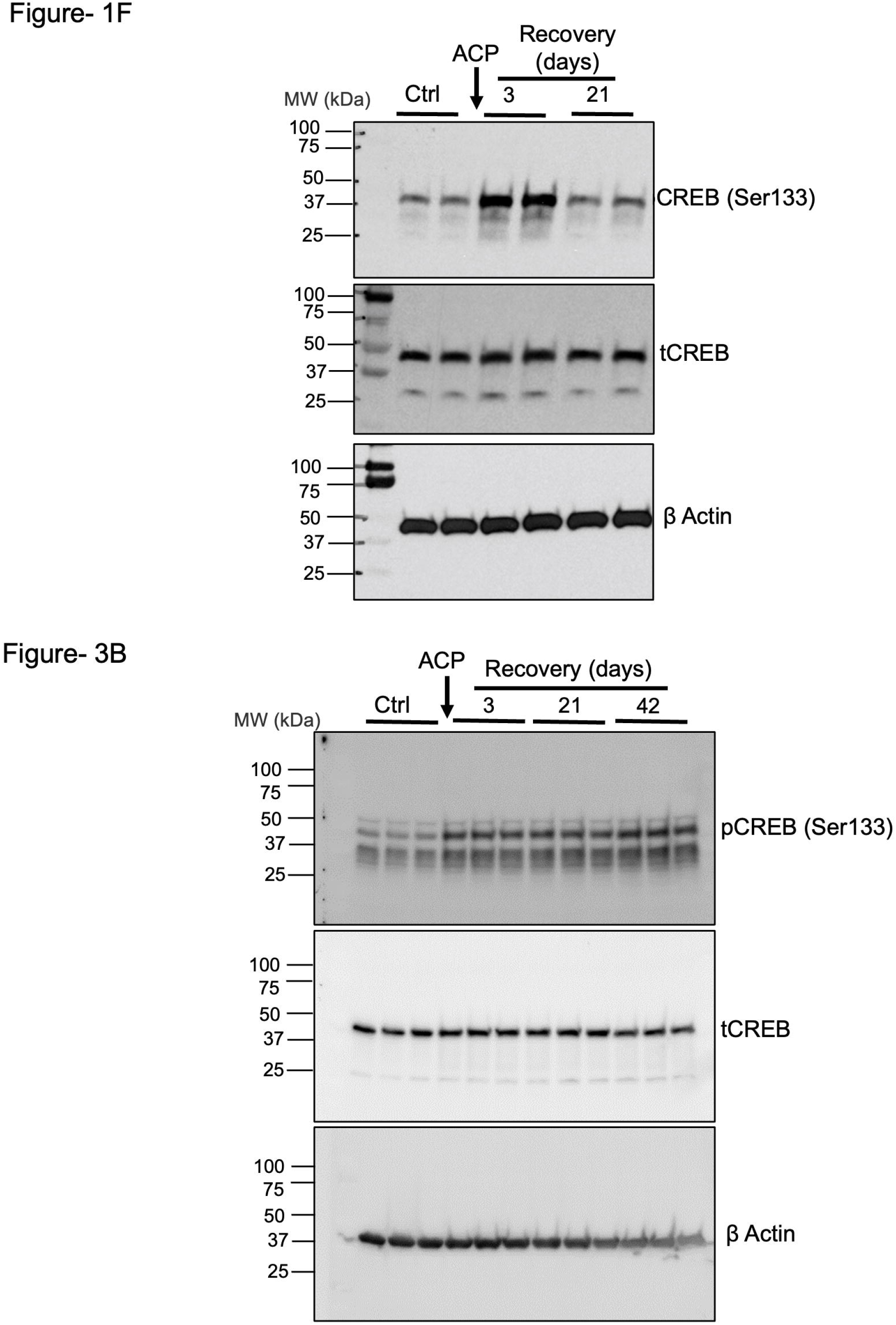

