## Supplementary Figure Legends for "CREB drives acinar cells to ductal reprogramming and promotes pancreatic cancer progression in preclinical models of alcoholic pancreatitis"

**Supplementary Figure 1.** Comparative analysis of severe damage mediated by alcoholic chronic pancreatitis (ACP) as compared to the impact of alcohol (A) or caerulein (CP) alone in *Ptf1a*$^{CreERTM/+}$ mice. (A) Dot plot depicting initial and final body weight measurements in mice treated with vehicle, (A), (CP), or ACP with 3- and 21-day recovery (n=7 mice per group). (B) Blood alcohol concentration levels (mg/dL) shown in control (ctrl), A, and ACP-induced *Ptf1a*$^{CreERTM/+}$ mice (n=7-9 mice per group). (C and D) Relative pancreas weight measurements [(pancreas weight (gm)/body weight(gm) X 1000] of *Ptf1a*$^{CreERTM/+}$ mice in ctrl, A, CP, and ACP-induced groups (C) with 3- and 21-day recovery (D) (n=5-7 mice per group). (E) Measurements of serum amylase activity in ctrl, A, CP, and ACP-induced mice (3-day recovery) (n=6 mice per group). (F) Representative images depicting H&E with histological quantification of CK19$^{+}$ ducts, PanINs (Alcian Blue), αSMA, collagen (Sirius red), and immune cells (CD45$^{+}$) in pancreata of ctrl, A, CP and ACP-induced *Ptf1a*$^{CreERTM/+}$ mice (3-day recovery) (n=4 mice per group). Scale bar, 50µm. $^{ns}$ nonsignificant; *p<0.05; **p< 0.01; ***p<0.001; ****p<0.0001 by ANOVA and unpaired t test.

**Supplementary Figure 2**. Molecular profiling of *Ptf1a*$^{CreERTM/+}$ mice pancreata after experimental induction of ACP. (A) Quantitative (qPCR) analysis in pancreas tissue harvested from ctrl and ACP-induced *Ptf1a*$^{CreERTM/+}$ mice (n=3 mice per group). (B) Representative image of mouse kinase array membranes and its quantification in ctrl and ACP-induced pancreata. Mouse kinase array in control (ctrl) and ACP-induced pancreata performed by using pooled tissue lysates from (N=3) biological replicates for each group (Ctrl and ACP), with two membranes used for quantitative estimation of fold change differences. Scale bar, 50µm. $^{ns}$ nonsignificant; *p<0.05; **p< 0.01; ***p<0.001;****p<0.0001 by ANOVA and unpaired t test.

**Supplementary Figure 3.** Comparative analysis of pancreatic damage mediated by ACP as compared to the impact of alcohol (A) or caerulein (CP) alone in *KC* mice. (A) Blood alcohol concentration measurement (mg/dL) in ctrl, A, and ACP-induced *KC* mice (n=6-7 mice per group). (B) Dot plot depicting comparing relative pancreatic weights between male and female *KC* mice with ACP (n=3-5 mice per group) (C) Representative pancreas images displaying H&E, images, and quantification of $CK19^+$ ducts, PanINs (Alcian Blue), collagen (Sirius red), αSMA, and immune cell positivity ($CD45^+$) in the pancreata of ctrl, A, CP, and ACP-induced *KC* experimental cohorts (3-day recovery) (n=4-7 mice per group). (D) Comparative histological evaluation of mouse pancreas using H&E-based analysis (n=3 mice per group). (E) Representative images illustrating pCREB expression in the pancreata of KC mice, accompanied by corresponding quantification across all experimental mice cohort (n=3-4 mice per group). (F) Bubble plot outlining the expression of canonical lineage cell cluster annotations in *KC* scRNA seq dataset. Scale bar, 50μm. $^{ns}$ nonsignificant**;** *p<0.05; **p< 0.01; ***p<0.001; ****p<0.0001 by ANOVA.

**Supplementary Figure 4.** Histological profiling of $Ptf1a^{CreERTM/+;}Creb^{fl/fl}$ ($CC^{-/-}$) mice pancreata with ACP induction. (A) Mouse breeding strategy to generate a genetic knockout of acinar cell specific *Creb*, under a $Ptf1a^{CreERTM/+}$ promoter. (B) Representative images of the mouse pancreas with H&E staining, along with co-immunofluorescence of total CREB expression (tCREB, red) and DAPI (blue), in control $Ptf1a^{CreERTM/+}$ (*C*) and $CC^{-/-}$ mice. I.F -based histological assessment, confirmed loss of CREB expression within the pancreata of $Ptf1a^{CreERTM/+};Creb^{fl/fl}$ ($CC^{-/-}$) as compared to wild type $Ptf1a^{CreERTM/+}$. (C) Comparative histological evaluation of mouse pancreas, accompanied by representative photomicrographs showcasing H&E, $CK19^+$ ducts, Sirius Red, αSMA, and $CD45^+$ staining within the pancreata of control (*C*) and $CC^{-/-}$ mice, in ctrl or with ACP induction (3-day recovery) (n=4 mice per group). Scale bar, 50μm. $^{ns}$ nonsignificant; ****p<0.0001 by ANOVA.

**Supplementary Figure 5.** Pancreas weight of *KC* and $Ptf1a^{CreERTM/+;}$ $LSL\text{-}Kras^{G12D/+}$; $Creb^{fl/fl}$ ($KCC^{-/-}$) mice with ACP induction. **(**A**)** Comparison of relative pancreas weight measurements in all experimental cohorts (n=3-5 mice per group). (B**)** Representative photomicrographs of whole pancreas depicting significantly less tumor burden in $KCC^{-/-}$ as compared to *KC* mice with ACP induction. Scale bar, 50μm. $^{ns}$ nonsignificant; ***p<0.001; ****p<0.0001 by ANOVA.

**Supplementary Figure 6.** Raw uncropped images of Western blot membranes for Figure *1F* and *3B*.
