## Supplementary Materials and Methods for "CREB drives acinar cells to ductal reprogramming and promotes pancreatic cancer progression in preclinical models of alcoholic pancreatitis"

*Serum Alcohol Analysis*

The alcohol concentration in the serum was estimated using an ethanol assay kit (Abcam, cat. # ab65343) as per the manufacturer's protocol and has been detailed previously.[1]

*Serum Amylase Analysis*

The serum amylase levels in blood samples collected from various study groups of mice were quantified through a colorimetric assay (Abcam, cat. # ab102523). This analysis was conducted in accordance with the manufacturer's provided protocol and described in detail previously.[1]

*Immunohistochemistry (IHC) and Immunofluorescence (IF) Tissue Staining*

Harvested pancreas tissues from mice necropsies were fixed in 10% neutral buffered formalin and embedded in paraffin to carry out histological staining procedures including Hematoxylin and Eosin (H&E), Sirius Red, Masson's trichrome, and Alcian Blue. For immunohistochemistry (IHC)-based detection, antigen retrieval was performed using citrate (pH 6.0) or Tris-EDTA buffer as previously described prior to incubation with BlockAid Blocking Solution (Thermo Fisher). And endogenous peroxidase activity was blocked by incubating with 3% $H_2O_2$. Tissue sections were then stained with primary antibodies at specified concentrations (Supplementary Table 3) overnight at $4^0$C. IHC slides were developed using 3,3' diaminobenzidine (DAB) substrate (Vector) followed by counterstain using Meyer's hematoxylin and imaged using DM750 microscope (Leica Microsystems). For IF-based staining, primary antibody was detected using species-specific Alexa Fluor 594 and/or Alexa Fluor 488 (Thermo Fisher) secondary antibodies incubated on sections for 1 hour at room temperature. Nuclear staining was performed using Hoechst 33342 dye (Thermo Fisher). All IF stained slides were scanned using the Olympus Fluoview1000 confocal microscope. In the present study, we performed digital image conversion to 16-bit grayscale followed by threshold adjustment in ImageJ to distinguish true fluorescent signals from background autofluorescence. This thresholding strategy was validated and we consistently excluded areas with non-specific signal or histological artifacts from quantification. Secondly, fluorescence imaging was performed using an Olympus Fluoview1000 confocal microscope. The use of a confocal system, particularly the adjustable pinhole aperture, allowed for optical sectioning and rejection of out-of-focus background signal, thereby improving signal-to-noise ratio. While paraffin-embedded pancreatic tissues are known to exhibit green-channel autofluorescence, we minimized its impact by optimizing laser power, detector gain, and imaging settings. High resolution multichannel images were exported for quantitative analysis, quantification of double positive cells was performed using ImageJ (NIH), for each mouse pancreas tissue sections, multiple non overlapping field of view per section were selected ensuring representative sampling across acinar and ductal compartments. The $CK19^+$ ducts, Alcian $blue^+$ PanINs, and $CD45^+$ immune cells were counted using Image J by converting the scanned image to 16-bit grayscale and adjusting the thresholds to differentiate the stained cells from the background. After that, the "analyze particles" function was used to automatically count the cells reported as discrete numbers.

For special staining including Sirius Red. The image scale was first calibrated to micrometers, followed by conversion to grayscale. A threshold was then applied to segment the Sirius Red–stained collagen areas, and the stained area was measured. Collagen deposition was reported as relative extracellular matrix (ECM) content normalized to the corresponding group in comparison for the analysis. IHC staining quantification of pCREB expression was done using ImageJ. The color deconvolution tool was applied to separate the true DAB positive signal from hematoxylin-stained purple counterstain. Measurement of percentage (%) area occupied was calculated across 3-4 representative field of view within the pancreas for each biological replicate across all 4 experimental mice cohorts using consistent upper and lower threshold limits to minimize background noise and ensure specificity of DAB accumulation within the tissues.

*H&E-based Assessment of Pancreatic Lesions*

Pancreatic tissue sections from age-matched mice in each group were stained with H&E and subsequently examined for ADMs, pancreatic lesions, and carcinoma in situ. The percentage of acinar area and number of ducts that contained any grade of PanIN lesions were measured by examining 10 H&E-stained high-power fields (40× magnification) per slide. PanINs were graded according to established criteria [2, 3]. In PanIN1 ducts, the normal cuboidal pancreatic epithelial cells transition to columnar architecture and can gain polyploid morphology. PanIN2 lesions are associated with a loss of polarity. PanIN3 lesions (or in situ carcinoma) show cribriform morphology, the budding off cells and luminal necrosis with marked cytological abnormalities, without invasion beyond the basement membrane. [2] The data was expressed as a percentage of total lesions in the whole pancreas.

*Western Blot Analysis*

The freshly harvested pancreas tissue was promptly flash-frozen and preserved at -80°C for long-term storage. To prepare protein lysates for Western blot analysis, the frozen tissues were thawed and homogenized in RIPA buffer (0.1% SDS, 50 mM Tris·HCl, 150 mM NaCl, 1% NP-40, and 0.5% Na deoxycholate) with protease inhibitor cocktail (Sigma, St. Louis, MO) and PhosSTOP phosphatase inhibitor (Roche, Indianapolis, IN, USA). Lysates were sonicated and centrifuged at 10,000 g for 15 minutes at $4^{0}$C to collect supernatant. The protein concentration of the cell and tissue lysate was determined by Bio-Rad protein assay kit (Bio-Rad, Hercules, CA). Next, 35 µg of whole-cell lysate or whole-tissue lysate was separated on NuPAGE Novex 4-12% Bis-Tris Gels and transferred on iBlot transfer stack using iBlot dry blotting transfer system (Life Technologies). For immune-detection, membranes were incubated with antibodies listed in Supplementary Table 3. The membranes were subsequently incubated with secondary anti-mouse or anti-rabbit secondary antibodies conjugated with horseradish peroxidase (Jackson ImmunoResearch). Finally, the immunoreactive bands were developed with Pierce ECL Western Blotting Substrate (Thermo Scientific) and recorded on blue basic autoradiography film (Bioexpress). Uncropped images of the blots are shown in Supplementary Figure 6.

*Phosphokinase Array Analysis*

The phosphorylation profiles of multiple effector kinases were screened using a phospho-kinase antibody array (R&D System, cat. #ARY003B). In brief, 200 mg of equal protein from pancreatic tissue lysates were loaded onto a nitrocellulose membrane with duplicate capture antibody spots. Phosphorylated protein levels were determined with phospho-specific antibodies and detected through chemiluminescent-based reaction chemistry. Spot density on the membrane was quantified using $HL^{++}$ image analysis software.

*Cluster Identification and Annotation of Single-cell RNA Sequencing Dataset*

Gene-cell matrices were analyzed with Seurat package (v4.3.0 RStudio). Cells fewer than 200 transcripts and ≤ 7.5% mitochondrial counts were removed. Feature measurements were normalized using the Normalize Data function with a scale factor of 10,000 and the *LogNormalize* normalization method. Variable genes were identified using the FindVariableFeatures function. Data were scaled and centered using linear regression on the counts and the cell cycle score difference. Principal Component Analysis (PCA) was performed with the *RunPCA* function using the previously defined variable genes to identify necessary dimensions for >90% variance within the data. Violin plots were then used to filter the data according to user-defined criteria. Cell clusters were obtained via the *FindNeighbors* and *FindClusters* functions, using a resolution of 0.7 for all samples, and non-linear dimensional reduction was then performed using Uniform Manifold Approximation and Projection (UMAP) clustering. A *FindAllMarkers* table was created, and clusters were defined by user-defined criteria. Clusters which co-expressed known distinct marker genes were merged for subsequent analysis.

*Differential Gene Expression Analysis of Single-cell RNA Sequencing Dataset*

Prior to differential gene expression analysis, each cluster of interest was subjected to normalization, scaling, and PCA. Next, the function "FindMarkers" from Seurat v4.0 R package was utilized to finds the differentially expressed genes for identity classes. Genes were considered differentially expressed if detected in at least 25% of clusters, with default log fold change of 0.25. Wilcoxon Rank Sum test was used and listed in Supplementary Table 4. Volcano plots were generated based on these output files using the *EnhancedVolcano* package. Gene set enrichment analysis (GSEA) was performed using the "fgsea" R package on differentially expressed genes (log(FC) >0.5 and adjusted p-value<0.05). "MSigDB" R package was utilized to access the following databases: C2 (KEGG, REACTOME, PID, BIOCARTA), C5 (GO:BP) and H (Hallmarks). Among different databases, the gene sets that met the statistical requirements were then curated, visualized via "ggplot2" R package, and ordered by Normalized Enrichment Score (NES) listed in Supplementary Table 4.

*RNA Isolation and qPCR Analysis*

RNA was isolated from flash frozen pancreas tissues using the RNeasy Kit (Qiagen) according to the manufacturer's protocol. cDNA generated after performing reverse transcription of RNA product was subjected to quantitative PCR (qPCR) analysis using gene-specific predesigned primers ($RT^2$ qPCR Primer Assay, Qiagen), listed in Supplementary Table 5. Gene expression was normalized to the housekeeping gene *GAPDH* using the comparative CT ($^{\Delta\Delta}$CT) method and reported as fold change relative to control.

*Isolation of Primary Pancreatic Acinar cells and 3D Explant Culture*

Isolation of primary pancreatic acinar cells and establishment of 3D explant cultures was performed as described previously.[4] Briefly, the acinar cells were incorporated into a mixture of collagen and Waymouth medium. On days 1, 3, and 5, duct-like structures were counted and quantified in a blinded fashion. The area of the ducts was determined using ImageJ software. Quantification of ductal structures involved the manual counting of at least four distinct fields under 10× magnification in triplicates.

*Statistical Analysis*

Descriptive statistics were calculated using Prism software (GraphPad Software Inc). Results are shown as values of means ± SD unless otherwise indicated. Prior to applying parametric tests, data distributions were assessed for normality using Shapiro- Wilk test. To assess multiple comparisons, one way ANOVA was applied followed by Tukey's or Dunnett's post hoc tests when deemed appropriate. A two-tailed Student's t test was used for two group comparisons. Statistical significance was defined using a cutoff of 0.050 unless otherwise specified in the figure legends
