## Supplementary Table 1 for "CREB drives acinar cells to ductal reprogramming and promotes pancreatic cancer progression in preclinical models of alcoholic pancreatitis"

**Supplementary Table 1.** Sequence of probes used for mice genotyping analysis.

| Gene | Forward primer | Reverse primer |
|---|---|---|
| Cre | GAA GGC ATT TGT GTA GGG TCA | GGC TGA GTG AGG GTT GTG AG |
| $Creb^{fl}$ | CTCTTCTTGCATCAAGCTTGGT | AGATCCCTCTAGGCATTTCTTCCT |
| $Creb^{WT}$ | CCAGTTACCTTCTAGGGAGCAGCTTACA | CAGGCCTGAGGTCTGGCTTCA |
| $Kras^{G12D}$ | GGCCTGCTGAAAATGACTGAGTATA | CTGTATCGTCAAGGCGCTCTT |
| $Rosa^{EYFP}$ | AGG GCG AGG AGC TGT TCA | TGA AGT CGA TGC CCT TCA G |
