## Supplementary Table 2 for "CREB drives acinar cells to ductal reprogramming and promotes pancreatic cancer progression in preclinical models of alcoholic pancreatitis"

**Supplementary Table 2.** List of treatment groups from ACP induction and recovery periods.

| *Groups* | *Induction period* | *Recovery period* | *Group Abbreviation* |
|---|---|---|---|
| 1 | Control diet<br>(14 weeks) | Continued control diet | **Ctrl** |
| 2 | Alcohol diet<br>(14 weeks) | Control diet<br>(3 or 21 days) | **A** |
| 3 | Control diet +<br>Caerulein injections<br>(for 4 weeks) | Continued control diet | **CP** |
| 4 | Alcohol diet +<br>Caerulein injections<br>(for 4 weeks) | Control diet<br>(3 or 21 days) | ACP<br>(3- or 21-day recovery)<br>= **ACP** |
| 5 | Alcohol diet +<br>Caerulein injections<br>(for 4 weeks) | Control diet<br>(3 days) | ACP<br>(3-day recovery)<br>= **ACP + 3d** |
| 6 | Alcohol diet +<br>Caerulein injections<br>(for 4 weeks) | Control diet<br>(21 days) | ACP<br>(21-day recovery)<br>= **ACP + 21d** |
