## Supplementary Table 3 for "CREB drives acinar cells to ductal reprogramming and promotes pancreatic cancer progression in preclinical models of alcoholic pancreatitis"

**Supplementary Table 3.** Primary antibodies for Western blot, immunohistochemistry, and immunofluorescence analysis.

| **Primary antibodies** | **Supplier** | **Species** | **Catalogue Number** |
|---|---|---|---|
| Phospho-CREB (Ser 133) | Cell signaling | Rabbit | 9198 |
| Total-CREB | Cell signaling | Rabbit | 9197 |
| Actin | Cell signaling | Mouse | 3700 |
| CK19 | EMD Millipore | Rat | MABT913 |
| GFP | Aves labs | Chicken | GFP-1010 |
| α-amylase | Cell signaling | Rabbit | 3796 |
| CD45 | Cell signaling | Rabbit | 70257 |
| α-SMA | Cell signaling | Rabbit | 19245 |
