## Supplementary Table 4 for "CREB drives acinar cells to ductal reprogramming and promotes pancreatic cancer progression in preclinical models of alcoholic pancreatitis"

| | genes | p_val | avg_log2F | pct.1 | pct.2 | p_val_adj |
|---|---|---|---|---|---|---|
| 11 | Reg3b | 2.29E-52 | 5.67172 | 0.536 | 0.037 | 4.80E-48 |
| 1 | Clu | 1.81E-81 | 5.40206 | 0.681 | 0.031 | 3.79E-77 |
| 31 | Reg2 | 6.19E-36 | 5.20806 | 0.449 | 0.043 | 1.30E-31 |
| 873 | Gm2691 | 0.1750401 | 4.64279 | 0.29 | 0.368 | 1 |
| 34 | S100a6 | 6.11E-34 | 4.24768 | 0.435 | 0.043 | 1.28E-29 |
| 84 | Reg1 | 1.34E-14 | 3.93498 | 0.71 | 0.375 | 2.80E-10 |
| 2 | S100a9 | 2.71E-79 | 3.82623 | 0.551 | 0.011 | 5.68E-75 |
| 7 | Krt8 | 8.20E-58 | 3.76217 | 0.406 | 0.009 | 1.72E-53 |
| 17 | Reg3a | 4.30E-48 | 3.65792 | 0.319 | 0.005 | 9.00E-44 |
| 6 | Spp1 | 6.67E-58 | 3.56157 | 0.377 | 0.005 | 1.40E-53 |
| 45 | Mt1 | 1.22E-28 | 3.50454 | 0.565 | 0.104 | 2.56E-24 |
| 1010 | Clps | 0.5809465 | 3.35757 | 0.58 | 0.729 | 1 |
| 175 | Pnliprp1 | 1.43E-07 | 3.3511 | 0.522 | 0.312 | 0.003 |
| 3 | Krt18 | 5.01E-67 | 3.32218 | 0.478 | 0.011 | 1.05E-62 |
| 58 | Nupr1 | 1.00E-22 | 3.2175 | 0.507 | 0.104 | 2.10E-18 |
| 9 | S100a8 | 1.52E-54 | 3.10616 | 0.464 | 0.019 | 3.18E-50 |
| 451 | Ctrb1 | 0.0007572 | 3.1034 | 0.696 | 0.759 | 1 |
| 69 | Tff2 | 1.78E-18 | 3.01746 | 0.565 | 0.161 | 3.72E-14 |
| 197 | Try5 | 4.81E-07 | 2.90876 | 0.551 | 0.338 | 0.01008 |
| 13 | Epcam | 3.85E-50 | 2.87405 | 0.362 | 0.009 | 8.06E-46 |
| 4 | Krt19 | 4.76E-63 | 2.84983 | 0.377 | 0.003 | 9.97E-59 |
| 74 | Dmbt1 | 4.24E-17 | 2.82317 | 0.42 | 0.093 | 8.87E-13 |
| 21 | Reg3g | 1.14E-44 | 2.81866 | 0.275 | 0.003 | 2.39E-40 |
| 12 | Krt7 | 3.19E-50 | 2.68343 | 0.319 | 0.004 | 6.68E-46 |
| 78 | S100a11 | 1.60E-16 | 2.56557 | 0.58 | 0.201 | 3.35E-12 |
| 35 | Sox4 | 3.94E-33 | 2.54513 | 0.391 | 0.033 | 8.24E-29 |
| 412 | Prss2 | 0.000334 | 2.44678 | 0.696 | 0.638 | 1 |
| 49 | Ctsl | 8.65E-28 | 2.20907 | 0.333 | 0.029 | 1.81E-23 |
| 55 | Dstn | 4.78E-25 | 2.18832 | 0.362 | 0.042 | 1.00E-20 |
| 60 | Mt2 | 1.55E-22 | 2.17233 | 0.319 | 0.034 | 3.24E-18 |
| 199 | Klk1 | 5.23E-07 | 2.16228 | 0.275 | 0.092 | 0.01096 |
| 38 | Anxa2 | 3.84E-31 | 1.97243 | 0.333 | 0.023 | 8.04E-27 |
| 43 | Anxa1 | 6.79E-30 | 1.95263 | 0.304 | 0.019 | 1.42E-25 |
| 319 | Cpa2 | 3.44E-05 | 1.9497 | 0.362 | 0.185 | 0.72044 |
| 41 | Nedd4 | 4.35E-30 | 1.93248 | 0.406 | 0.042 | 9.11E-26 |
| 5 | Col1a2 | 1.60E-58 | 1.91263 | 0.391 | 0.006 | 3.35E-54 |
| 22 | Cdh1 | 2.81E-43 | 1.90689 | 0.333 | 0.01 | 5.88E-39 |
| 16 | Cystm1 | 7.22E-49 | 1.90097 | 0.377 | 0.011 | 1.51E-44 |
| 8 | Sox9 | 5.37E-55 | 1.87755 | 0.348 | 0.004 | 1.12E-50 |
| 59 | Ifitm3 | 1.13E-22 | 1.87687 | 0.362 | 0.047 | 2.37E-18 |
| 29 | Col1a1 | 2.99E-38 | 1.84915 | 0.304 | 0.01 | 6.27E-34 |
| 223 | Spink1 | 1.37E-06 | 1.81359 | 0.333 | 0.135 | 0.02872 |

| | | | | | | |
|---|---|---|---|---|---|---|
| 65 | Cd24a | 2.07E-20 | 1.73658 | 0.406 | 0.068 | 4.34E-16 |
| 42 | Fabp5 | 4.45E-30 | 1.72583 | 0.319 | 0.022 | 9.31E-26 |
| 19 | Cldn7 | 3.45E-47 | 1.72468 | 0.275 | 0.001 | 7.23E-43 |
| 160 | Prdx1 | 4.05E-08 | 1.71942 | 0.536 | 0.28 | 0.00085 |
| 61 | Tceal9 | 2.32E-22 | 1.71818 | 0.391 | 0.056 | 4.86E-18 |
| 94 | Txn1 | 8.52E-13 | 1.68379 | 0.493 | 0.173 | 1.78E-08 |
| 80 | Spint2 | 2.98E-16 | 1.66419 | 0.406 | 0.094 | 6.23E-12 |
| 88 | Pmepa1 | 3.32E-14 | 1.65391 | 0.275 | 0.046 | 6.96E-10 |
| 32 | Gsto1 | 1.52E-35 | 1.55847 | 0.304 | 0.013 | 3.17E-31 |
| 89 | Dap | 7.11E-14 | 1.55102 | 0.391 | 0.092 | 1.49E-09 |
| 15 | Col3a1 | 3.44E-49 | 1.5061 | 0.42 | 0.017 | 7.21E-45 |
| 63 | Marcks | 4.60E-21 | 1.50038 | 0.362 | 0.05 | 9.64E-17 |
| 76 | Atpif1 | 6.42E-17 | 1.49241 | 0.449 | 0.107 | 1.34E-12 |
| 37 | Errfi1 | 1.66E-31 | 1.49066 | 0.319 | 0.019 | 3.48E-27 |
| 971 | Cela3b | 0.4240016 | 1.48255 | 0.42 | 0.583 | 1 |
| 258 | Actn1 | 4.06E-06 | 1.4815 | 0.449 | 0.224 | 0.08495 |
| 535 | Cel | 0.0036955 | 1.4759 | 0.391 | 0.278 | 1 |
| 120 | Rps27l | 5.06E-10 | 1.47028 | 0.464 | 0.185 | 1.06E-05 |
| 251 | Selenop | 3.42E-06 | 1.45568 | 0.333 | 0.135 | 0.07154 |
| 213 | Sec61b | 8.40E-07 | 1.44774 | 0.551 | 0.311 | 0.01759 |
| 27 | Spint1 | 6.77E-40 | 1.44496 | 0.261 | 0.004 | 1.42E-35 |
| 30 | Cldn3 | 7.46E-38 | 1.42536 | 0.261 | 0.005 | 1.56E-33 |
| 52 | Pbx1 | 7.48E-26 | 1.39565 | 0.261 | 0.015 | 1.57E-21 |
| 20 | Prr15l | 5.98E-45 | 1.39082 | 0.29 | 0.004 | 1.25E-40 |
| 25 | Ctnnd1 | 5.28E-41 | 1.38472 | 0.29 | 0.006 | 1.11E-36 |
| 51 | Wars | 4.24E-26 | 1.37956 | 0.304 | 0.024 | 8.88E-22 |
| 23 | Nectin2 | 1.86E-42 | 1.34259 | 0.275 | 0.004 | 3.90E-38 |
| 407 | Ly6a | 0.0003117 | 1.31019 | 0.319 | 0.158 | 1 |
| 305 | Tagln2 | 2.20E-05 | 1.29154 | 0.42 | 0.227 | 0.46084 |
| 159 | Tpm1 | 3.62E-08 | 1.28897 | 0.319 | 0.103 | 0.00076 |
| 257 | Jun | 4.05E-06 | 1.28757 | 0.348 | 0.138 | 0.08483 |
| 18 | Prom1 | 2.55E-47 | 1.26865 | 0.29 | 0.003 | 5.34E-43 |
| 812 | Cpa1 | 0.0951219 | 1.26498 | 0.536 | 0.561 | 1 |
| 128 | Anxa5 | 1.79E-09 | 1.25651 | 0.348 | 0.108 | 3.74E-05 |
| 105 | Fos | 2.38E-11 | 1.24918 | 0.29 | 0.061 | 4.99E-07 |
| 152 | Fth1 | 1.54E-08 | 1.24341 | 0.826 | 0.543 | 0.00032 |
| 46 | Nfib | 1.80E-28 | 1.23873 | 0.261 | 0.013 | 3.77E-24 |
| 83 | Clic4 | 1.08E-14 | 1.2234 | 0.304 | 0.052 | 2.26E-10 |
| 40 | Stard10 | 9.72E-31 | 1.22145 | 0.333 | 0.023 | 2.04E-26 |
| 252 | Lmo4 | 3.52E-06 | 1.22051 | 0.29 | 0.099 | 0.07363 |
| 54 | Rhob | 3.28E-25 | 1.18997 | 0.275 | 0.019 | 6.87E-21 |
| 62 | Serpinb6 | 1.51E-21 | 1.18676 | 0.333 | 0.039 | 3.17E-17 |
| 33 | Serf1 | 8.27E-35 | 1.15293 | 0.275 | 0.009 | 1.73E-30 |

| | | | | | | |
|---|---|---|---|---|---|---|
| 146 | Ftl1 | 1.32E-08 | 1.14212 | 0.681 | 0.427 | 0.00028 |
| 57 | Dusp1 | 2.68E-24 | 1.13073 | 0.275 | 0.02 | 5.60E-20 |
| 260 | Ybx1 | 4.59E-06 | 1.13022 | 0.58 | 0.364 | 0.09621 |
| 226 | Nme2 | 1.64E-06 | 1.12254 | 0.638 | 0.403 | 0.03428 |
| 448 | Cyb5a | 0.0007314 | 1.11621 | 0.362 | 0.189 | 1 |
| 53 | Cd63 | 2.76E-25 | 1.11198 | 0.275 | 0.019 | 5.78E-21 |
| 391 | Gapdh | 0.000219 | 1.11116 | 0.464 | 0.292 | 1 |
| 671 | Try4 | 0.0219013 | 1.11025 | 0.391 | 0.631 | 1 |
| 14 | Gpx2 | 8.56E-50 | 1.11021 | 0.29 | 0.001 | 1.79E-45 |
| 456 | Sec61g | 0.0008548 | 1.10457 | 0.478 | 0.303 | 1 |
| 1053 | Lars2 | 0.8438974 | 1.08696 | 0.623 | 0.559 | 1 |
| 220 | Gpx4 | 1.23E-06 | 1.08656 | 0.493 | 0.248 | 0.0258 |
| 138 | Ppp1r14l | 7.14E-09 | 1.08216 | 0.377 | 0.129 | 0.00015 |
| 10 | Cd14 | 1.58E-52 | 1.08006 | 0.29 | 0 | 3.30E-48 |
| 171 | Cd9 | 1.11E-07 | 1.07338 | 0.261 | 0.075 | 0.00233 |
| 136 | Txndc17 | 5.87E-09 | 1.07294 | 0.362 | 0.115 | 0.00012 |
| 68 | Alcam | 1.05E-18 | 1.0699 | 0.333 | 0.047 | 2.21E-14 |
| 79 | Ifitm2 | 2.09E-16 | 1.06961 | 0.319 | 0.05 | 4.38E-12 |
| 1064 | Cpb1 | 0.9143226 | 1.06723 | 0.406 | 0.532 | 1 |
| 75 | App | 5.18E-17 | 1.06649 | 0.275 | 0.036 | 1.08E-12 |
| 48 | Anxa4 | 2.12E-28 | 1.0659 | 0.261 | 0.013 | 4.45E-24 |
| 26 | Rhou | 3.02E-40 | 1.05765 | 0.275 | 0.005 | 6.33E-36 |
| 279 | Ssr4 | 9.42E-06 | 1.04236 | 0.435 | 0.218 | 0.19726 |
| 333 | Ndufc1 | 4.97E-05 | 1.03947 | 0.333 | 0.141 | 1 |
| 145 | Cd44 | 1.17E-08 | 1.03794 | 0.275 | 0.074 | 0.00025 |
| 137 | Atxn10 | 6.76E-09 | 1.03697 | 0.29 | 0.076 | 0.00014 |
| 341 | Vcp | 6.71E-05 | 1.0313 | 0.391 | 0.196 | 1 |
| 73 | Ctnna1 | 4.16E-17 | 1.02903 | 0.29 | 0.039 | 8.70E-13 |
| 224 | Hint1 | 1.43E-06 | 1.02406 | 0.522 | 0.283 | 0.02993 |
| 36 | St14 | 2.88E-32 | 1.01818 | 0.261 | 0.009 | 6.03E-28 |
| 271 | Eif3b | 6.17E-06 | 1.01643 | 0.333 | 0.126 | 0.12932 |
| 44 | Tmem17 | 4.98E-29 | 1.0142 | 0.304 | 0.019 | 1.04E-24 |
| 284 | Cox5b | 1.10E-05 | 1.00433 | 0.478 | 0.242 | 0.23128 |
| 115 | Prdx2 | 3.23E-10 | 0.99522 | 0.42 | 0.14 | 6.76E-06 |
| 82 | Ptms | 2.13E-15 | 0.99482 | 0.348 | 0.065 | 4.47E-11 |
| 39 | Asns | 4.64E-31 | 0.99352 | 0.261 | 0.01 | 9.71E-27 |
| 72 | Saa3 | 3.19E-17 | 0.99206 | 0.304 | 0.042 | 6.67E-13 |
| 198 | Ctsd | 4.85E-07 | 0.98971 | 0.42 | 0.186 | 0.01015 |
| 103 | Nfic | 1.46E-11 | 0.98761 | 0.275 | 0.054 | 3.06E-07 |
| 232 | Ifi27 | 2.04E-06 | 0.98722 | 0.319 | 0.113 | 0.04272 |
| 368 | Hdgf | 0.0001318 | 0.9843 | 0.348 | 0.175 | 1 |
| 144 | Ifrd1 | 1.13E-08 | 0.98412 | 0.29 | 0.078 | 0.00024 |
| 496 | Romo1 | 0.0021445 | 0.98162 | 0.29 | 0.146 | 1 |

| | | | | | | |
|---|---|---|---|---|---|---|
| 28 | Myo1b | 7.39E-39 | 0.97908 | 0.29 | 0.008 | 1.55E-34 |
| 426 | Mbd3 | 0.0004479 | 0.97621 | 0.275 | 0.118 | 1 |
| 201 | Psmd8 | 5.44E-07 | 0.97482 | 0.348 | 0.129 | 0.01139 |
| 241 | Uqcr11 | 2.37E-06 | 0.97476 | 0.406 | 0.183 | 0.04953 |
| 133 | Tpd52 | 4.14E-09 | 0.97331 | 0.29 | 0.076 | 8.68E-05 |
| 277 | Slc25a3 | 8.15E-06 | 0.9684 | 0.594 | 0.362 | 0.17067 |
| 135 | Lgals1 | 5.43E-09 | 0.96712 | 0.391 | 0.131 | 0.00011 |
| 24 | Cttn | 3.64E-42 | 0.96586 | 0.261 | 0.003 | 7.63E-38 |
| 81 | Sfn | 1.14E-15 | 0.96183 | 0.261 | 0.034 | 2.38E-11 |
| 178 | Sh3bgrl | 1.53E-07 | 0.95753 | 0.275 | 0.079 | 0.0032 |
| 365 | Ndufs6 | 0.0001081 | 0.95153 | 0.348 | 0.158 | 1 |
| 183 | Bri3 | 2.74E-07 | 0.94089 | 0.377 | 0.144 | 0.00573 |
| 338 | Pebp1 | 5.89E-05 | 0.93905 | 0.348 | 0.166 | 1 |
| 404 | Cox7a2 | 0.0002914 | 0.93629 | 0.464 | 0.279 | 1 |
| 395 | Uqcr10 | 0.0002536 | 0.92878 | 0.406 | 0.217 | 1 |
| 325 | Prdx5 | 3.93E-05 | 0.92766 | 0.333 | 0.15 | 0.82404 |
| 500 | Gm4241 | 0.0022893 | 0.92302 | 1 | 1 | 1 |
| 166 | Glud1 | 8.43E-08 | 0.91835 | 0.377 | 0.131 | 0.00176 |
| 216 | Hdlbp | 9.54E-07 | 0.91175 | 0.362 | 0.136 | 0.01998 |
| 164 | Prxl2c | 6.02E-08 | 0.91031 | 0.319 | 0.096 | 0.00126 |
| 273 | Gm1007 | 6.82E-06 | 0.9091 | 0.478 | 0.238 | 0.14283 |
| 434 | Mrpl52 | 0.0005427 | 0.90677 | 0.362 | 0.191 | 1 |
| 608 | Actg1 | 0.010125 | 0.89402 | 0.594 | 0.492 | 1 |
| 127 | Idh2 | 1.73E-09 | 0.89146 | 0.261 | 0.057 | 3.62E-05 |
| 124 | P4hb | 1.35E-09 | 0.89093 | 0.391 | 0.121 | 2.84E-05 |
| 433 | Uqcrq | 0.0005014 | 0.8867 | 0.391 | 0.227 | 1 |
| 561 | Nme1 | 0.0056069 | 0.88591 | 0.391 | 0.239 | 1 |
| 234 | Apoe | 2.06E-06 | 0.88314 | 0.333 | 0.121 | 0.04312 |
| 297 | Rheb | 1.68E-05 | 0.87568 | 0.333 | 0.134 | 0.35202 |
| 337 | Kdm1a | 5.81E-05 | 0.87497 | 0.275 | 0.11 | 1 |
| 262 | Pnlip | 4.69E-06 | 0.8734 | 0.275 | 0.67 | 0.09829 |
| 114 | Rab11a | 3.15E-10 | 0.87213 | 0.304 | 0.074 | 6.60E-06 |
| 480 | Atp5j | 0.0013917 | 0.86698 | 0.435 | 0.276 | 1 |
| 149 | Eny2 | 1.39E-08 | 0.86615 | 0.362 | 0.116 | 0.00029 |
| 635 | Tbca | 0.0150377 | 0.8642 | 0.348 | 0.219 | 1 |
| 436 | Ndufb4 | 0.0005837 | 0.85789 | 0.362 | 0.176 | 1 |
| 476 | Eef2 | 0.0013169 | 0.85286 | 0.667 | 0.52 | 1 |
| 181 | Cstb | 2.04E-07 | 0.85212 | 0.29 | 0.092 | 0.00428 |
| 662 | Psap | 0.020157 | 0.85138 | 0.406 | 0.273 | 1 |
| 402 | Psmb4 | 0.0002819 | 0.8491 | 0.406 | 0.211 | 1 |
| 703 | Ywhaq | 0.0308032 | 0.84698 | 0.304 | 0.205 | 1 |
| 189 | Nars | 3.32E-07 | 0.83891 | 0.362 | 0.131 | 0.00695 |
| 547 | Gpx1 | 0.0046442 | 0.83887 | 0.362 | 0.228 | 1 |

| | | | | | | |
|---|---|---|---|---|---|---|
| 47 | Lmna | 1.83E-28 | 0.83743 | 0.261 | 0.013 | 3.84E-24 |
| 499 | Cox8a | 0.0022748 | 0.83428 | 0.522 | 0.383 | 1 |
| 409 | Srp72 | 0.0003163 | 0.83269 | 0.29 | 0.125 | 1 |
| 85 | Ybx3 | 1.38E-14 | 0.82564 | 0.261 | 0.038 | 2.89E-10 |
| 371 | Sem1 | 0.0001409 | 0.82443 | 0.522 | 0.316 | 1 |
| 610 | Timm13 | 0.0104879 | 0.82377 | 0.348 | 0.219 | 1 |
| 903 | Pitpnc1 | 0.2200874 | 0.82285 | 0.275 | 0.199 | 1 |
| 170 | Atox1 | 1.05E-07 | 0.81591 | 0.391 | 0.154 | 0.0022 |
| 219 | Capns1 | 1.20E-06 | 0.81525 | 0.333 | 0.127 | 0.02508 |
| 542 | Tomm6 | 0.0043638 | 0.80556 | 0.42 | 0.261 | 1 |
| 77 | Ak3 | 1.37E-16 | 0.79564 | 0.261 | 0.032 | 2.87E-12 |
| 112 | Tnip1 | 2.47E-10 | 0.79289 | 0.29 | 0.065 | 5.17E-06 |
| 549 | Arf1 | 0.0048177 | 0.79262 | 0.391 | 0.254 | 1 |
| 156 | Xbp1 | 3.09E-08 | 0.7916 | 0.29 | 0.08 | 0.00065 |
| 632 | Mtpn | 0.0144354 | 0.78918 | 0.275 | 0.154 | 1 |
| 209 | Map1lc3l | 6.81E-07 | 0.78176 | 0.464 | 0.195 | 0.01426 |
| 677 | Ctnnb1 | 0.0235548 | 0.78077 | 0.333 | 0.241 | 1 |
| 111 | Pabpc4 | 2.34E-10 | 0.78022 | 0.275 | 0.059 | 4.90E-06 |
| 126 | Bag1 | 1.62E-09 | 0.77965 | 0.391 | 0.121 | 3.38E-05 |
| 642 | Mrps33 | 0.0159513 | 0.77952 | 0.275 | 0.152 | 1 |
| 598 | Chchd2 | 0.0088187 | 0.77321 | 0.507 | 0.395 | 1 |
| 278 | Coa3 | 8.75E-06 | 0.77306 | 0.319 | 0.122 | 0.18332 |
| 108 | Ormdl2 | 1.20E-10 | 0.76816 | 0.275 | 0.057 | 2.51E-06 |
| 317 | Zyx | 3.26E-05 | 0.76563 | 0.304 | 0.12 | 0.68236 |
| 347 | Gadd45g | 7.66E-05 | 0.76189 | 0.275 | 0.104 | 1 |
| 168 | Rnh1 | 9.39E-08 | 0.75976 | 0.275 | 0.078 | 0.00197 |
| 71 | Tsc22d1 | 1.90E-17 | 0.75654 | 0.275 | 0.033 | 3.97E-13 |
| 270 | Ndufs2 | 6.09E-06 | 0.7557 | 0.319 | 0.117 | 0.12748 |
| 299 | Anapc13 | 1.78E-05 | 0.7551 | 0.29 | 0.104 | 0.37296 |
| 380 | Psmb5 | 0.0001609 | 0.75414 | 0.348 | 0.164 | 1 |
| 445 | Ranbp1 | 0.0007076 | 0.75042 | 0.348 | 0.178 | 1 |
| 221 | Sec31a | 1.32E-06 | 0.74588 | 0.304 | 0.101 | 0.02769 |
| 131 | Hist1h1e | 3.66E-09 | 0.74119 | 0.261 | 0.06 | 7.67E-05 |
| 354 | Rpl41 | 9.77E-05 | 0.74014 | 0.841 | 0.669 | 1 |
| 410 | Rrbp1 | 0.0003177 | 0.73564 | 0.42 | 0.231 | 1 |
| 568 | Cox6c | 0.0061248 | 0.73506 | 0.464 | 0.329 | 1 |
| 785 | Gabarap | 0.073431 | 0.73013 | 0.362 | 0.282 | 1 |
| 265 | Cfdp1 | 5.51E-06 | 0.72971 | 0.29 | 0.102 | 0.11547 |
| 292 | Psmd2 | 1.40E-05 | 0.72946 | 0.275 | 0.097 | 0.29346 |
| 56 | Nckap1 | 5.12E-25 | 0.72892 | 0.261 | 0.017 | 1.07E-20 |
| 345 | Ehmt1 | 7.54E-05 | 0.7256 | 0.275 | 0.106 | 1 |
| 96 | Lyz2 | 1.69E-12 | 0.71696 | 0.319 | 0.064 | 3.55E-08 |
| 501 | Vdac2 | 0.0023533 | 0.70876 | 0.319 | 0.178 | 1 |

| | | | | | | |
|---|---|---|---|---|---|---|
| 458 | Polr2f | 0.0009281 | 0.70326 | 0.29 | 0.139 | 1 |
| 50 | Grn | 5.09E-27 | 0.70243 | 0.261 | 0.014 | 1.07E-22 |
| 576 | Cox7b | 0.0068511 | 0.7021 | 0.377 | 0.238 | 1 |
| 615 | Eprs | 0.0110779 | 0.69647 | 0.275 | 0.153 | 1 |
| 647 | H2afz | 0.0168355 | 0.69611 | 0.435 | 0.317 | 1 |
| 474 | Cox6a1 | 0.0013029 | 0.69572 | 0.391 | 0.231 | 1 |
| 793 | Eif4g2 | 0.0825338 | 0.69551 | 0.522 | 0.404 | 1 |
| 1070 | Sycn | 0.9487775 | 0.69497 | 0.406 | 0.487 | 1 |
| 109 | Rnpep | 1.77E-10 | 0.68849 | 0.261 | 0.054 | 3.70E-06 |
| 421 | Trim28 | 0.0003868 | 0.6884 | 0.275 | 0.116 | 1 |
| 416 | Itpr3 | 0.0003525 | 0.68726 | 0.29 | 0.127 | 1 |
| 122 | Cnpy2 | 7.54E-10 | 0.68516 | 0.275 | 0.061 | 1.58E-05 |
| 893 | Rnase1 | 0.2039495 | 0.68304 | 0.348 | 0.527 | 1 |
| 366 | Cox14 | 0.0001197 | 0.68043 | 0.333 | 0.145 | 1 |
| 459 | Atp5b | 0.0009504 | 0.67316 | 0.551 | 0.367 | 1 |
| 293 | Mrps24 | 1.42E-05 | 0.67217 | 0.391 | 0.171 | 0.29837 |
| 733 | 2210010 | 0.0428377 | 0.66864 | 0.391 | 0.665 | 1 |
| 355 | Arf4 | 9.77E-05 | 0.66524 | 0.348 | 0.163 | 1 |
| 403 | Cdv3 | 0.0002908 | 0.66321 | 0.348 | 0.172 | 1 |
| 616 | Ppib | 0.0115188 | 0.66267 | 0.435 | 0.312 | 1 |
| 344 | Rpn1 | 7.37E-05 | 0.65354 | 0.29 | 0.122 | 1 |
| 764 | Eif2s2 | 0.0631633 | 0.65089 | 0.362 | 0.271 | 1 |
| 646 | Micos10 | 0.0165703 | 0.64771 | 0.333 | 0.208 | 1 |
| 1047 | Cela2a | 0.8084976 | 0.64151 | 0.449 | 0.539 | 1 |
| 359 | Ndufb8 | 0.000101 | 0.64108 | 0.333 | 0.145 | 1 |
| 375 | Ctsb | 0.000146 | 0.64039 | 0.261 | 0.108 | 1 |
| 827 | Atp6v1f | 0.111082 | 0.63954 | 0.319 | 0.224 | 1 |
| 761 | Snrpe | 0.059746 | 0.63684 | 0.42 | 0.308 | 1 |
| 736 | Atxn7l3b | 0.0437722 | 0.63668 | 0.275 | 0.172 | 1 |
| 492 | Junb | 0.0020627 | 0.63532 | 0.377 | 0.22 | 1 |
| 538 | Serp1 | 0.0040116 | 0.63398 | 0.435 | 0.269 | 1 |
| 384 | Pfdn6 | 0.0001769 | 0.6339 | 0.319 | 0.138 | 1 |
| 634 | Ptma | 0.0149121 | 0.62954 | 0.71 | 0.599 | 1 |
| 238 | Dbi | 2.29E-06 | 0.6275 | 0.362 | 0.15 | 0.04791 |
| 530 | Bsg | 0.0034303 | 0.62564 | 0.29 | 0.164 | 1 |
| 621 | H2afj | 0.0119439 | 0.6232 | 0.435 | 0.287 | 1 |
| 435 | Prdx6 | 0.0005474 | 0.62313 | 0.406 | 0.222 | 1 |
| 267 | Eif3l | 5.78E-06 | 0.62222 | 0.304 | 0.107 | 0.12101 |
| 130 | Msrb1 | 2.90E-09 | 0.61976 | 0.261 | 0.059 | 6.07E-05 |
| 360 | Hsd17b1 | 0.0001012 | 0.6193 | 0.261 | 0.097 | 1 |
| 303 | Sqstm1 | 1.89E-05 | 0.61925 | 0.275 | 0.098 | 0.39637 |
| 594 | Scand1 | 0.0082904 | 0.61773 | 0.406 | 0.247 | 1 |
| 475 | Eif4b | 0.0013051 | 0.61551 | 0.391 | 0.21 | 1 |

| | | | | | | |
|---|---|---|---|---|---|---|
| 833 | Ndufa7 | 0.1152838 | 0.61518 | 0.304 | 0.227 | 1 |
| 1043 | S100a1( | 0.7827844 | 0.61513 | 0.333 | 0.335 | 1 |
| 557 | Ndufb9 | 0.0052635 | 0.61327 | 0.333 | 0.19 | 1 |
| 406 | Pgk1 | 0.0003082 | 0.61313 | 0.319 | 0.148 | 1 |
| 98 | Sephs2 | 4.01E-12 | 0.61113 | 0.275 | 0.051 | 8.40E-08 |
| 132 | Glul | 3.89E-09 | 0.60698 | 0.261 | 0.06 | 8.15E-05 |
| 461 | Mif | 0.0009743 | 0.60667 | 0.464 | 0.278 | 1 |
| 563 | Cox6b1 | 0.0057034 | 0.60577 | 0.449 | 0.302 | 1 |
| 362 | Actn4 | 0.0001026 | 0.60367 | 0.319 | 0.145 | 1 |
| 335 | Ppfia1 | 5.58E-05 | 0.6027 | 0.261 | 0.093 | 1 |
| 560 | Ost4 | 0.0055584 | 0.60224 | 0.391 | 0.236 | 1 |
| 513 | Phb2 | 0.0026998 | 0.6017 | 0.319 | 0.166 | 1 |
| 693 | Suclg1 | 0.0282042 | 0.59958 | 0.29 | 0.169 | 1 |
| 741 | Snu13 | 0.0478508 | 0.59179 | 0.319 | 0.204 | 1 |
| 973 | Ctrl | 0.4356505 | 0.58666 | 0.319 | 0.327 | 1 |
| 836 | Tomm20 | 0.1186214 | 0.58594 | 0.377 | 0.285 | 1 |
| 243 | Ube2r2 | 2.41E-06 | 0.5853 | 0.319 | 0.117 | 0.05042 |
| 906 | Neat1 | 0.2246603 | 0.58413 | 0.348 | 0.301 | 1 |
| 715 | Taldo1 | 0.0365521 | 0.58181 | 0.333 | 0.223 | 1 |
| 93 | Atp6v1a | 6.97E-13 | 0.58152 | 0.261 | 0.042 | 1.46E-08 |
| 643 | Ndufb1-p | 0.0163839 | 0.57942 | 0.406 | 0.273 | 1 |
| 153 | Echs1 | 2.17E-08 | 0.57464 | 0.261 | 0.064 | 0.00045 |
| 663 | Mrpl33 | 0.0205186 | 0.57393 | 0.348 | 0.218 | 1 |
| 162 | Sod1 | 4.59E-08 | 0.57307 | 0.348 | 0.11 | 0.00096 |
| 744 | Sat1 | 0.0499613 | 0.57125 | 0.29 | 0.2 | 1 |
| 440 | Rtn4 | 0.0006487 | 0.56851 | 0.319 | 0.162 | 1 |
| 439 | Skp1a | 0.0006377 | 0.56657 | 0.29 | 0.135 | 1 |
| 399 | Gm4998 | 0.000269 | 0.56656 | 0.333 | 0.148 | 1 |
| 653 | Cox7c | 0.0180225 | 0.56378 | 0.522 | 0.395 | 1 |
| 726 | Eif1 | 0.0418766 | 0.56095 | 0.667 | 0.561 | 1 |
| 281 | Ubald2 | 9.71E-06 | 0.56057 | 0.319 | 0.117 | 0.20344 |
| 696 | Mrps14 | 0.0285879 | 0.55985 | 0.319 | 0.191 | 1 |
| 388 | Srsf9 | 0.0001873 | 0.55784 | 0.29 | 0.121 | 1 |
| 819 | Atp5e | 0.1016252 | 0.55459 | 0.536 | 0.422 | 1 |
| 649 | Pfdn2 | 0.017173 | 0.54974 | 0.304 | 0.175 | 1 |
| 674 | Cox4i1 | 0.0228174 | 0.54769 | 0.551 | 0.411 | 1 |
| 463 | Hnrnpab | 0.0010286 | 0.54207 | 0.406 | 0.229 | 1 |
| 933 | Ndufa4 | 0.2986596 | 0.5415 | 0.348 | 0.296 | 1 |
| 644 | Ndufv3 | 0.0164288 | 0.53361 | 0.304 | 0.173 | 1 |
| 601 | Psmd7 | 0.0095508 | 0.53282 | 0.275 | 0.148 | 1 |
| 482 | Grpel1 | 0.0014414 | 0.53042 | 0.275 | 0.126 | 1 |
| 401 | Aurkaip1 | 0.000281 | 0.53017 | 0.304 | 0.13 | 1 |
| 334 | Atp1b1 | 5.49E-05 | 0.52917 | 0.275 | 0.107 | 1 |

| | | | | | | |
|---|---|---|---|---|---|---|
| 669 | Atp5g3 | 0.0216222 | 0.52757 | 0.362 | 0.239 | 1 |
| 483 | Rps12 | 0.0014641 | 0.52481 | 0.87 | 0.676 | 1 |
| 702 | Psmd4 | 0.0306851 | 0.5243 | 0.29 | 0.175 | 1 |
| 452 | Laptm4a | 0.0007664 | 0.52368 | 0.304 | 0.148 | 1 |
| 218 | Copz1 | 1.14E-06 | 0.51979 | 0.261 | 0.076 | 0.02381 |
| 716 | Selenof | 0.0379321 | 0.51941 | 0.333 | 0.223 | 1 |
| 553 | Rtcb | 0.0050083 | 0.51881 | 0.304 | 0.158 | 1 |
| 815 | Atp5g1 | 0.0959806 | 0.51216 | 0.333 | 0.238 | 1 |
| 838 | Psma7 | 0.125007 | 0.51148 | 0.348 | 0.27 | 1 |
| 652 | Imp3 | 0.0176856 | 0.50454 | 0.275 | 0.155 | 1 |
| 308 | Smim15 | 2.30E-05 | 0.5027 | 0.261 | 0.09 | 0.48222 |
| 584 | Mrps21 | 0.0073484 | 0.50243 | 0.29 | 0.158 | 1 |
| 512 | Eif6 | 0.0026784 | 0.49978 | 0.261 | 0.131 | 1 |
| 431 | Psmb6 | 0.0004779 | 0.49717 | 0.304 | 0.136 | 1 |
| 324 | Cst3 | 3.85E-05 | 0.49608 | 0.406 | 0.19 | 0.80595 |
| 686 | Ddx3x | 0.0266607 | 0.49405 | 0.348 | 0.22 | 1 |
| 630 | Rpl7a | 0.0142658 | 0.49281 | 0.696 | 0.508 | 1 |
| 443 | Psmc4 | 0.0006867 | 0.48926 | 0.275 | 0.117 | 1 |
| 1066 | AY03611 | 0.9341519 | 0.48557 | 0.667 | 0.656 | 1 |
| 628 | Gspt1 | 0.0135867 | 0.48483 | 0.319 | 0.182 | 1 |
| 484 | Atp5k | 0.0014704 | 0.48465 | 0.377 | 0.196 | 1 |
| 816 | Aimp1 | 0.0983922 | 0.48246 | 0.275 | 0.18 | 1 |
| 820 | Eif3c | 0.1028063 | 0.48171 | 0.348 | 0.252 | 1 |
| 555 | Prmt1 | 0.0051592 | 0.47816 | 0.261 | 0.124 | 1 |
| 300 | Mydgf | 1.79E-05 | 0.47763 | 0.261 | 0.088 | 0.37565 |
| 141 | Gfpt1 | 8.42E-09 | 0.47646 | 0.275 | 0.069 | 0.00018 |
| 495 | Kcmf1 | 0.0020992 | 0.47457 | 0.275 | 0.129 | 1 |
| 490 | Cycs | 0.001884 | 0.46806 | 0.319 | 0.162 | 1 |
| 581 | Bax | 0.0071852 | 0.46726 | 0.304 | 0.162 | 1 |
| 469 | Esd | 0.0012201 | 0.46507 | 0.29 | 0.141 | 1 |
| 358 | Etf1 | 0.0001006 | 0.46234 | 0.275 | 0.11 | 1 |
| 769 | Pomp | 0.0684758 | 0.45935 | 0.319 | 0.214 | 1 |
| 779 | Myl12a | 0.0714075 | 0.45756 | 0.377 | 0.28 | 1 |
| 698 | Vapa | 0.0295449 | 0.4571 | 0.304 | 0.187 | 1 |
| 636 | Rad23b | 0.0151587 | 0.45544 | 0.319 | 0.183 | 1 |
| 465 | Rab5c | 0.0011456 | 0.45524 | 0.29 | 0.13 | 1 |
| 110 | Aldh2 | 1.92E-10 | 0.45261 | 0.304 | 0.069 | 4.02E-06 |
| 236 | Cat | 2.23E-06 | 0.45224 | 0.275 | 0.085 | 0.04667 |
| 797 | Gas5 | 0.0848662 | 0.45141 | 0.507 | 0.403 | 1 |
| 655 | Lamp1 | 0.0192148 | 0.44738 | 0.275 | 0.166 | 1 |
| 577 | Zfp706 | 0.0068912 | 0.44618 | 0.377 | 0.227 | 1 |
| 466 | Sars | 0.001169 | 0.44533 | 0.29 | 0.132 | 1 |
| 712 | Rpl10 | 0.0349786 | 0.44476 | 0.739 | 0.569 | 1 |

| | | | | | | |
|---|---|---|---|---|---|---|
| 352 | Rab5if | 8.94E-05 | 0.44291 | 0.319 | 0.136 | 1 |
| 468 | Ndufa12 | 0.0012145 | 0.43898 | 0.275 | 0.121 | 1 |
| 1035 | Ywhaz | 0.7228864 | 0.43785 | 0.391 | 0.404 | 1 |
| 66 | Csrp2 | 8.01E-20 | 0.43751 | 0.261 | 0.024 | 1.68E-15 |
| 606 | Ralbp1 | 0.0099312 | 0.43363 | 0.304 | 0.166 | 1 |
| 772 | Eif3k | 0.0691568 | 0.43281 | 0.435 | 0.316 | 1 |
| 233 | Vdac1 | 2.05E-06 | 0.4284 | 0.319 | 0.111 | 0.04295 |
| 600 | Ndufs5 | 0.0090541 | 0.42533 | 0.319 | 0.178 | 1 |
| 622 | Nop58 | 0.012231 | 0.42361 | 0.275 | 0.149 | 1 |
| 485 | Eef1a1 | 0.0016791 | 0.42342 | 0.942 | 0.734 | 1 |
| 494 | Pdcd6 | 0.0020926 | 0.42012 | 0.304 | 0.153 | 1 |
| 554 | Mrfap1 | 0.0050796 | 0.41758 | 0.362 | 0.208 | 1 |
| 1042 | Zg16 | 0.7800821 | 0.41753 | 0.319 | 0.362 | 1 |
| 579 | Akr1a1 | 0.0071589 | 0.41667 | 0.29 | 0.169 | 1 |
| 377 | Adam10 | 0.0001506 | 0.41219 | 0.348 | 0.152 | 1 |
| 582 | Hspa9 | 0.0073057 | 0.40744 | 0.319 | 0.172 | 1 |
| 781 | Polr1d | 0.0716972 | 0.40741 | 0.348 | 0.242 | 1 |
| 666 | G3bp1 | 0.0210149 | 0.40731 | 0.391 | 0.254 | 1 |
| 464 | Mien1 | 0.0010537 | 0.40718 | 0.304 | 0.143 | 1 |
| 864 | Slc25a5 | 0.1603598 | 0.40451 | 0.362 | 0.287 | 1 |
| 670 | Cbfb | 0.0217557 | 0.40368 | 0.261 | 0.139 | 1 |
| 510 | Ssr3 | 0.0026223 | 0.40361 | 0.275 | 0.135 | 1 |
| 692 | Rpl28 | 0.0279349 | 0.40257 | 0.812 | 0.59 | 1 |
| 834 | Tspo | 0.1167147 | 0.40226 | 0.333 | 0.262 | 1 |
| 734 | Atp5d | 0.0431982 | 0.40068 | 0.406 | 0.298 | 1 |
| 619 | Ctsz | 0.011751 | 0.39998 | 0.275 | 0.148 | 1 |
| 414 | Lpp | 0.0003407 | 0.39856 | 0.304 | 0.136 | 1 |
| 875 | Gng5 | 0.1797863 | 0.39636 | 0.42 | 0.364 | 1 |
| 803 | Paip2 | 0.0884078 | 0.3961 | 0.348 | 0.239 | 1 |
| 340 | Gm4330 | 6.15E-05 | 0.39255 | 0.275 | 0.103 | 1 |
| 851 | Arpc5 | 0.1409874 | 0.39044 | 0.406 | 0.317 | 1 |
| 353 | Ubqln1 | 8.95E-05 | 0.39033 | 0.304 | 0.12 | 1 |
| 302 | Ap2m1 | 1.88E-05 | 0.3901 | 0.319 | 0.122 | 0.39292 |
| 880 | Selenok | 0.1852814 | 0.38997 | 0.304 | 0.218 | 1 |
| 511 | Smarca4 | 0.0026349 | 0.38653 | 0.275 | 0.131 | 1 |
| 835 | H3f3b | 0.1173246 | 0.38567 | 0.667 | 0.613 | 1 |
| 453 | Zfand5 | 0.0008027 | 0.38529 | 0.304 | 0.136 | 1 |
| 310 | Nsun2 | 2.51E-05 | 0.38343 | 0.261 | 0.09 | 0.52512 |
| 521 | Psma6 | 0.0030001 | 0.38317 | 0.29 | 0.143 | 1 |
| 607 | Tmem25 | 0.0100363 | 0.38233 | 0.333 | 0.191 | 1 |
| 182 | Bccip | 2.22E-07 | 0.38176 | 0.319 | 0.098 | 0.00464 |
| 597 | Atf4 | 0.0085843 | 0.37987 | 0.304 | 0.164 | 1 |
| 479 | Tkt | 0.0013712 | 0.37924 | 0.275 | 0.131 | 1 |

| | | | | | | |
|---|---|---|---|---|---|---|
| 143 | Lrrc58 | 1.00E-08 | 0.37835 | 0.29 | 0.073 | 0.00021 |
| 558 | Gnb2 | 0.0053091 | 0.37831 | 0.406 | 0.248 | 1 |
| 809 | Snrpd3 | 0.0932424 | 0.37705 | 0.348 | 0.232 | 1 |
| 266 | Cyb5b | 5.65E-06 | 0.37561 | 0.261 | 0.083 | 0.1183 |
| 925 | Cdc37 | 0.2692143 | 0.37498 | 0.304 | 0.229 | 1 |
| 351 | Uri1 | 8.47E-05 | 0.37379 | 0.29 | 0.111 | 1 |
| 364 | Ndufs3 | 0.0001063 | 0.37161 | 0.304 | 0.122 | 1 |
| 722 | Trp53inp | 0.0403957 | 0.37157 | 0.304 | 0.182 | 1 |
| 573 | Ghitm | 0.0067221 | 0.37131 | 0.275 | 0.141 | 1 |
| 1040 | Anp32b | 0.7728953 | 0.37105 | 0.319 | 0.288 | 1 |
| 907 | Atp5j2 | 0.2253182 | 0.37018 | 0.42 | 0.334 | 1 |
| 886 | Jpt1 | 0.1906091 | 0.36869 | 0.348 | 0.264 | 1 |
| 940 | Rpl22l1 | 0.3191634 | 0.36862 | 0.507 | 0.465 | 1 |
| 658 | Snrpd2 | 0.019721 | 0.36815 | 0.348 | 0.218 | 1 |
| 527 | Rnf10 | 0.0033682 | 0.36804 | 0.275 | 0.132 | 1 |
| 832 | Tpm4 | 0.1117443 | 0.36121 | 0.348 | 0.256 | 1 |
| 518 | Sfr1 | 0.0029542 | 0.36069 | 0.319 | 0.162 | 1 |
| 831 | Eif5a | 0.1117112 | 0.36022 | 0.362 | 0.283 | 1 |
| 562 | Smarce1 | 0.0056835 | 0.35858 | 0.319 | 0.167 | 1 |
| 719 | Atp1a1 | 0.0386795 | 0.35836 | 0.319 | 0.21 | 1 |
| 590 | Ndufc2 | 0.0079624 | 0.35772 | 0.261 | 0.136 | 1 |
| 514 | 1110004 | 0.0027382 | 0.35557 | 0.275 | 0.131 | 1 |
| 648 | Ptbp1 | 0.01696 | 0.35501 | 0.29 | 0.162 | 1 |
| 397 | Strap | 0.0002613 | 0.35397 | 0.261 | 0.103 | 1 |
| 749 | Swi5 | 0.0518127 | 0.3537 | 0.304 | 0.203 | 1 |
| 602 | Timm10b | 0.0095699 | 0.35367 | 0.304 | 0.159 | 1 |
| 311 | R3hdm4 | 2.64E-05 | 0.35306 | 0.29 | 0.106 | 0.55234 |
| 488 | Uqcrb | 0.0017942 | 0.35257 | 0.377 | 0.213 | 1 |
| 842 | Hmgn1 | 0.1363333 | 0.35245 | 0.333 | 0.252 | 1 |
| 804 | Ndufa2 | 0.0907642 | 0.35104 | 0.391 | 0.29 | 1 |
| 481 | Rnf7 | 0.0013971 | 0.351 | 0.261 | 0.113 | 1 |
| 724 | Rrp1 | 0.0407347 | 0.35062 | 0.261 | 0.153 | 1 |
| 796 | Plekhj1 | 0.0838938 | 0.34843 | 0.275 | 0.175 | 1 |
| 735 | Caprin1 | 0.0436771 | 0.34803 | 0.391 | 0.268 | 1 |
| 801 | Ywhae | 0.0877262 | 0.34675 | 0.362 | 0.274 | 1 |
| 239 | Cops3 | 2.31E-06 | 0.3441 | 0.261 | 0.078 | 0.04832 |
| 887 | Trir | 0.1913225 | 0.34354 | 0.348 | 0.26 | 1 |
| 580 | Bola2 | 0.0071675 | 0.34347 | 0.377 | 0.213 | 1 |
| 264 | Rala | 5.39E-06 | 0.34174 | 0.275 | 0.092 | 0.11288 |
| 890 | Gabarap | 0.2007636 | 0.33966 | 0.261 | 0.18 | 1 |
| 551 | Psmc6 | 0.0049293 | 0.33919 | 0.29 | 0.145 | 1 |
| 272 | Copb2 | 6.49E-06 | 0.33844 | 0.29 | 0.098 | 0.13582 |
| 844 | Myl6 | 0.1376014 | 0.3383 | 0.551 | 0.465 | 1 |

| | | | | | | |
|---|---|---|---|---|---|---|
| 673 | Eif3d | 0.0221962 | 0.33715 | 0.29 | 0.159 | 1 |
| 507 | mt-Nd3 | 0.0025148 | 0.33713 | 0.638 | 0.908 | 1 |
| 780 | Krtcap2 | 0.0714826 | 0.33672 | 0.333 | 0.238 | 1 |
| 894 | Atp6v1g1 | 0.2060602 | 0.33614 | 0.275 | 0.206 | 1 |
| 1057 | Hnrnpc | 0.8714752 | 0.33488 | 0.333 | 0.322 | 1 |
| 1001 | Ran | 0.5385671 | 0.33405 | 0.333 | 0.297 | 1 |
| 912 | Tes | 0.2356018 | 0.33257 | 0.275 | 0.211 | 1 |
| 748 | Rps26 | 0.0514373 | 0.33236 | 0.826 | 0.597 | 1 |
| 191 | Ergic3 | 4.00E-07 | 0.33175 | 0.261 | 0.07 | 0.00837 |
| 751 | Atp5o | 0.0534752 | 0.33142 | 0.319 | 0.204 | 1 |
| 766 | Calm2 | 0.0645758 | 0.3293 | 0.391 | 0.288 | 1 |
| 565 | Spcs1 | 0.0058662 | 0.32895 | 0.29 | 0.15 | 1 |
| 190 | Cyth2 | 3.45E-07 | 0.3285 | 0.261 | 0.071 | 0.00723 |
| 738 | Rpl3 | 0.0449889 | 0.32788 | 0.783 | 0.568 | 1 |
| 699 | Atp5g2 | 0.029753 | 0.32532 | 0.478 | 0.35 | 1 |
| 747 | Rpl14 | 0.0513176 | 0.32516 | 0.725 | 0.567 | 1 |
| 861 | Cct8 | 0.1541488 | 0.32457 | 0.348 | 0.243 | 1 |
| 672 | Pdia6 | 0.0221125 | 0.32328 | 0.275 | 0.159 | 1 |
| 626 | S100a13 | 0.0126264 | 0.32224 | 0.261 | 0.138 | 1 |
| 575 | Arih1 | 0.006802 | 0.32115 | 0.333 | 0.175 | 1 |
| 707 | Rps2 | 0.0333485 | 0.31982 | 0.739 | 0.586 | 1 |
| 603 | Atp5md | 0.009619 | 0.31898 | 0.348 | 0.21 | 1 |
| 417 | Prr13 | 0.0003626 | 0.31773 | 0.261 | 0.107 | 1 |
| 556 | Hsbp1 | 0.005241 | 0.31687 | 0.275 | 0.14 | 1 |
| 753 | Fkbp4 | 0.0546998 | 0.31648 | 0.319 | 0.21 | 1 |
| 637 | Ndufa10 | 0.0151761 | 0.30991 | 0.275 | 0.148 | 1 |
| 559 | Ak2 | 0.005494 | 0.308 | 0.261 | 0.129 | 1 |
| 574 | Cd164 | 0.0067474 | 0.30663 | 0.362 | 0.204 | 1 |
| 806 | Aldoa | 0.0911397 | 0.30472 | 0.319 | 0.236 | 1 |
| 572 | H13 | 0.0064308 | 0.30381 | 0.319 | 0.173 | 1 |
| 595 | Rab1a | 0.008338 | 0.3035 | 0.261 | 0.132 | 1 |
| 853 | Rwdd1 | 0.1428378 | 0.30288 | 0.261 | 0.173 | 1 |
| 627 | 1810037 | 0.0130511 | 0.30134 | 0.29 | 0.168 | 1 |
| 721 | Hnrnpa0 | 0.0402467 | 0.3012 | 0.362 | 0.241 | 1 |
| 446 | mt-Co1 | 0.0007155 | 0.30044 | 0.812 | 0.992 | 1 |
| 709 | Capzb | 0.0338662 | 0.29865 | 0.391 | 0.261 | 1 |
| 211 | Dazap1 | 8.00E-07 | 0.29811 | 0.319 | 0.103 | 0.01676 |
| 727 | Rpl35 | 0.041886 | 0.29607 | 0.681 | 0.501 | 1 |
| 972 | Nap1l1 | 0.4291128 | 0.29472 | 0.348 | 0.287 | 1 |
| 982 | Psmb1 | 0.4768835 | 0.29437 | 0.377 | 0.311 | 1 |
| 437 | Dynlrb1 | 0.0006014 | 0.29338 | 0.348 | 0.164 | 1 |
| 840 | Atp5l | 0.1324691 | 0.29309 | 0.464 | 0.354 | 1 |
| 296 | Api5 | 1.68E-05 | 0.29251 | 0.275 | 0.096 | 0.35114 |

| | | | | | | |
|---|---|---|---|---|---|---|
| 314 | Snhg6 | 2.94E-05 | 0.2905 | 0.261 | 0.092 | 0.61484 |
| 800 | Eif4g1 | 0.086831 | 0.28656 | 0.348 | 0.255 | 1 |
| 814 | Elavl1 | 0.0955054 | 0.28655 | 0.261 | 0.164 | 1 |
| 537 | Higd1a | 0.0038374 | 0.28529 | 0.29 | 0.145 | 1 |
| 988 | Eif3f | 0.4964284 | 0.28446 | 0.435 | 0.363 | 1 |
| 862 | Btf3 | 0.1546669 | 0.28023 | 0.493 | 0.404 | 1 |
| 789 | Rpl26 | 0.0789403 | 0.27902 | 0.826 | 0.637 | 1 |
| 415 | Denr | 0.0003503 | 0.279 | 0.261 | 0.107 | 1 |
| 723 | Psma4 | 0.0405139 | 0.27881 | 0.29 | 0.173 | 1 |
| 240 | Dtymk | 2.33E-06 | 0.27872 | 0.261 | 0.079 | 0.04883 |
| 369 | Sbds | 0.0001341 | 0.27706 | 0.29 | 0.115 | 1 |
| 657 | Tomm5 | 0.0196459 | 0.27631 | 0.275 | 0.157 | 1 |
| 967 | Tmem59 | 0.4156645 | 0.27394 | 0.319 | 0.269 | 1 |
| 710 | Selenos | 0.0340674 | 0.27285 | 0.275 | 0.161 | 1 |
| 782 | Metap2 | 0.0723867 | 0.27138 | 0.275 | 0.178 | 1 |
| 508 | Eif1ax | 0.0025574 | 0.27098 | 0.261 | 0.126 | 1 |
| 524 | Irf2bpl | 0.0030887 | 0.27074 | 0.29 | 0.144 | 1 |
| 847 | Ostc | 0.1390192 | 0.26937 | 0.261 | 0.172 | 1 |
| 923 | Arpc1b | 0.2654399 | 0.26906 | 0.464 | 0.391 | 1 |
| 526 | Aamp | 0.003268 | 0.26337 | 0.304 | 0.152 | 1 |
| 891 | Ndufb11 | 0.2016364 | 0.26198 | 0.333 | 0.25 | 1 |
| 706 | Tmed2 | 0.032558 | 0.26064 | 0.333 | 0.214 | 1 |
| 720 | Taf10 | 0.0399048 | 0.2587 | 0.377 | 0.245 | 1 |
| 400 | mt-Co3 | 0.0002706 | 0.25869 | 0.87 | 0.991 | 1 |
| 289 | Aph1a | 1.34E-05 | 0.2573 | 0.261 | 0.085 | 0.28165 |
| 742 | Cggbp1 | 0.0479705 | 0.25675 | 0.29 | 0.178 | 1 |
| 505 | Aprt | 0.0024363 | 0.25285 | 0.29 | 0.145 | 1 |
| 650 | Pkm | 0.0172805 | 0.2525 | 0.348 | 0.215 | 1 |
| 865 | Pop5 | 0.1621701 | 0.25154 | 0.261 | 0.169 | 1 |
| 571 | Reep5 | 0.0063302 | 0.25138 | 0.304 | 0.166 | 1 |
| 491 | Pdcd5 | 0.0020522 | 0.25132 | 0.275 | 0.129 | 1 |
| 924 | Selenow | 0.26565 | -0.25001 | 0.304 | 0.334 | 1 |
| 983 | Rpl8 | 0.4780636 | -0.25022 | 0.696 | 0.619 | 1 |
| 981 | Eif3h | 0.4736756 | -0.25026 | 0.348 | 0.345 | 1 |
| 995 | Ppp2ca | 0.5244516 | -0.25245 | 0.261 | 0.273 | 1 |
| 949 | Smc6 | 0.3362515 | -0.25246 | 0.275 | 0.293 | 1 |
| 943 | Smarca2 | 0.3244127 | -0.25256 | 0.261 | 0.187 | 1 |
| 1069 | Psme2 | 0.9484697 | -0.25333 | 0.304 | 0.265 | 1 |
| 897 | Npc2 | 0.2089924 | -0.25601 | 0.304 | 0.343 | 1 |
| 991 | Rps20 | 0.5140228 | -0.25616 | 0.841 | 0.734 | 1 |
| 1045 | Rps18 | 0.7984748 | -0.25627 | 0.812 | 0.671 | 1 |
| 1039 | Map4k4 | 0.7726613 | -0.25842 | 0.304 | 0.301 | 1 |
| 944 | Eef1d | 0.3279371 | -0.26032 | 0.319 | 0.346 | 1 |

| | | | | | | |
|---|---|---|---|---|---|---|
| 955 | Rpl11 | 0.3504815 | -0.26286 | 0.739 | 0.623 | 1 |
| 1032 | Chd8 | 0.7180137 | -0.26346 | 0.275 | 0.22 | 1 |
| 960 | Btg2 | 0.3808796 | -0.26588 | 0.304 | 0.335 | 1 |
| 913 | Pfn1 | 0.2356646 | -0.2711 | 0.507 | 0.524 | 1 |
| 994 | Psma3 | 0.5210466 | -0.27358 | 0.304 | 0.318 | 1 |
| 255 | mt-Nd2 | 3.82E-06 | -0.27782 | 0.725 | 0.971 | 0.07996 |
| 1031 | Cct7 | 0.7016918 | -0.27939 | 0.29 | 0.229 | 1 |
| 998 | Ywhab | 0.5321012 | -0.28208 | 0.261 | 0.273 | 1 |
| 948 | Atp6v0b | 0.3361887 | -0.28312 | 0.304 | 0.225 | 1 |
| 966 | Snrpg | 0.4006192 | -0.28426 | 0.406 | 0.4 | 1 |
| 1038 | Tomm22 | 0.7690565 | -0.28876 | 0.261 | 0.214 | 1 |
| 539 | Srgn | 0.0040502 | -0.29067 | 0.174 | 0.357 | 1 |
| 896 | Hspe1 | 0.2076293 | -0.29314 | 0.377 | 0.42 | 1 |
| 937 | Gna13 | 0.3102288 | -0.29363 | 0.275 | 0.192 | 1 |
| 1023 | Dusp11 | 0.6446827 | -0.29396 | 0.319 | 0.302 | 1 |
| 958 | Sf3b5 | 0.3684248 | -0.29812 | 0.261 | 0.189 | 1 |
| 977 | Csnk1a1 | 0.46468 | -0.29906 | 0.362 | 0.387 | 1 |
| 1071 | Psmb3 | 0.9573707 | -0.29909 | 0.275 | 0.245 | 1 |
| 1016 | Papola | 0.6175109 | -0.30028 | 0.29 | 0.284 | 1 |
| 883 | Smim14 | 0.1882558 | -0.30289 | 0.362 | 0.245 | 1 |
| 935 | Nol7 | 0.3033854 | -0.30417 | 0.232 | 0.251 | 1 |
| 783 | Gnai2 | 0.0726406 | -0.30461 | 0.319 | 0.42 | 1 |
| 1033 | Arfgef1 | 0.7211212 | -0.30711 | 0.261 | 0.219 | 1 |
| 910 | Rpl27a | 0.2266434 | -0.31431 | 0.783 | 0.711 | 1 |
| 1027 | Cbx3 | 0.6729016 | -0.31525 | 0.304 | 0.294 | 1 |
| 386 | mt-Nd5 | 0.0001849 | -0.31773 | 0.58 | 0.834 | 1 |
| 1029 | Rps29 | 0.6821843 | -0.31949 | 0.841 | 0.683 | 1 |
| 985 | Rpl30 | 0.4881745 | -0.31989 | 0.841 | 0.692 | 1 |
| 1009 | Tnrc6a | 0.5744541 | -0.32482 | 0.275 | 0.275 | 1 |
| 927 | Stk24 | 0.2736788 | -0.3256 | 0.261 | 0.297 | 1 |
| 1048 | Ddx21 | 0.8218164 | -0.32819 | 0.261 | 0.223 | 1 |
| 877 | Sypl | 0.1816592 | -0.32939 | 0.261 | 0.171 | 1 |
| 1063 | Ppp4r3a | 0.9118469 | -0.33172 | 0.261 | 0.241 | 1 |
| 936 | H3f3a | 0.309207 | -0.33321 | 0.594 | 0.61 | 1 |
| 1055 | Canx | 0.8678956 | -0.33376 | 0.29 | 0.264 | 1 |
| 843 | Cfl1 | 0.1373256 | -0.33442 | 0.391 | 0.483 | 1 |
| 956 | Rpl17 | 0.353474 | -0.33819 | 0.754 | 0.65 | 1 |
| 817 | Ctss | 0.0994338 | -0.34044 | 0.188 | 0.273 | 1 |
| 892 | Arid4b | 0.2037521 | -0.3406 | 0.232 | 0.282 | 1 |
| 1002 | Rbm6 | 0.5458729 | -0.34178 | 0.29 | 0.28 | 1 |
| 1044 | Gm1180 | 0.7981685 | -0.34505 | 0.261 | 0.213 | 1 |
| 1024 | Trip11 | 0.6549292 | -0.34578 | 0.246 | 0.25 | 1 |
| 855 | Ssb | 0.144276 | -0.34792 | 0.217 | 0.274 | 1 |

| | | | | | | |
|---|---|---|---|---|---|---|
| 1025 | Ube2i | 0.6643863 | -0.34966 | 0.261 | 0.265 | 1 |
| 1022 | Cct4 | 0.6365263 | -0.35292 | 0.275 | 0.269 | 1 |
| 984 | Rap1a | 0.4829629 | -0.35918 | 0.246 | 0.25 | 1 |
| 919 | Csde1 | 0.2469464 | -0.35943 | 0.348 | 0.377 | 1 |
| 975 | Sf3b2 | 0.4486519 | -0.35978 | 0.29 | 0.308 | 1 |
| 1065 | Ppp1cb | 0.9179089 | -0.36016 | 0.261 | 0.231 | 1 |
| 926 | Rpl19 | 0.2715052 | -0.3612 | 0.739 | 0.662 | 1 |
| 964 | Rpl37a | 0.3973781 | -0.36795 | 0.797 | 0.689 | 1 |
| 874 | Mcl1 | 0.1760134 | -0.36904 | 0.377 | 0.411 | 1 |
| 1000 | Clint1 | 0.5332711 | -0.37079 | 0.29 | 0.227 | 1 |
| 950 | Snx5 | 0.3378085 | -0.37156 | 0.275 | 0.194 | 1 |
| 1008 | Rabac1 | 0.5644145 | -0.37552 | 0.304 | 0.297 | 1 |
| 1017 | Spcs2 | 0.61911 | -0.37578 | 0.275 | 0.271 | 1 |
| 1004 | Anp32a | 0.5486065 | -0.3791 | 0.304 | 0.29 | 1 |
| 1030 | Trip12 | 0.6828779 | -0.38058 | 0.29 | 0.233 | 1 |
| 1073 | Srrm1 | 0.9765731 | -0.3817 | 0.319 | 0.293 | 1 |
| 1012 | Zfp207 | 0.5874698 | -0.38356 | 0.304 | 0.303 | 1 |
| 939 | Atp1b3 | 0.3184059 | -0.38699 | 0.304 | 0.326 | 1 |
| 1013 | Cast | 0.6063149 | -0.39043 | 0.29 | 0.228 | 1 |
| 990 | Fam104a | 0.511764 | -0.39123 | 0.261 | 0.2 | 1 |
| 1054 | Ddit4 | 0.845051 | -0.39236 | 0.261 | 0.219 | 1 |
| 1018 | Tmem23 | 0.6195358 | -0.39494 | 0.319 | 0.313 | 1 |
| 895 | Actb | 0.2064089 | -0.39818 | 0.855 | 0.783 | 1 |
| 993 | Eif3m | 0.5204775 | -0.39826 | 0.29 | 0.284 | 1 |
| 946 | Kif5b | 0.3322861 | -0.39836 | 0.362 | 0.378 | 1 |
| 1058 | Tuba1b | 0.8720229 | -0.39885 | 0.275 | 0.246 | 1 |
| 830 | Vim | 0.111676 | -0.39944 | 0.188 | 0.255 | 1 |
| 899 | Srsf2 | 0.2101743 | -0.40087 | 0.304 | 0.352 | 1 |
| 1052 | Tgfbr2 | 0.8422819 | -0.40169 | 0.319 | 0.285 | 1 |
| 1036 | Smarca5 | 0.7294128 | -0.40216 | 0.261 | 0.251 | 1 |
| 1021 | Rpl39 | 0.6300416 | -0.40338 | 0.812 | 0.634 | 1 |
| 1046 | Sec11c | 0.8003338 | -0.40415 | 0.261 | 0.245 | 1 |
| 980 | Vasp | 0.4684039 | -0.4042 | 0.246 | 0.264 | 1 |
| 697 | Prkcd | 0.0289622 | -0.40493 | 0.275 | 0.146 | 1 |
| 845 | Csk | 0.1376288 | -0.40817 | 0.246 | 0.29 | 1 |
| 918 | Rpl35a | 0.2437826 | -0.41849 | 0.725 | 0.634 | 1 |
| 544 | Fermt3 | 0.0046008 | -0.42258 | 0.116 | 0.278 | 1 |
| 860 | Zfp652 | 0.1531756 | -0.42412 | 0.261 | 0.163 | 1 |
| 1014 | Cltc | 0.611616 | -0.4272 | 0.246 | 0.26 | 1 |
| 942 | Thrap3 | 0.3229378 | -0.42769 | 0.261 | 0.282 | 1 |
| 1015 | Zfp36l1 | 0.614977 | -0.42846 | 0.464 | 0.448 | 1 |
| 978 | Chd4 | 0.4682289 | -0.4295 | 0.348 | 0.352 | 1 |
| 951 | Rps7 | 0.3383399 | -0.43013 | 0.739 | 0.636 | 1 |

| | | | | | | |
|---|---|---|---|---|---|---|
| 776 | Tpm3 | 0.0709011 | -0.43099 | 0.348 | 0.438 | 1 |
| 1011 | Aff4 | 0.5811682 | -0.4314 | 0.304 | 0.242 | 1 |
| 1019 | Cdk12 | 0.6279675 | -0.43185 | 0.275 | 0.265 | 1 |
| 1049 | Tardbp | 0.8250349 | -0.43338 | 0.333 | 0.308 | 1 |
| 745 | Itm2b | 0.0501671 | -0.43448 | 0.391 | 0.483 | 1 |
| 969 | Senp6 | 0.4223741 | -0.43904 | 0.246 | 0.27 | 1 |
| 881 | Ctsh | 0.1860703 | -0.44439 | 0.261 | 0.167 | 1 |
| 954 | Zfp91 | 0.3420548 | -0.44789 | 0.29 | 0.21 | 1 |
| 968 | Gatad2b | 0.4212551 | -0.44792 | 0.275 | 0.284 | 1 |
| 934 | Nsa2 | 0.2991949 | -0.44796 | 0.333 | 0.346 | 1 |
| 1062 | Pabpn1 | 0.897266 | -0.44834 | 0.275 | 0.237 | 1 |
| 957 | H2afy | 0.3544384 | -0.44928 | 0.333 | 0.345 | 1 |
| 911 | Ppp1r12a | 0.2293455 | -0.45368 | 0.232 | 0.274 | 1 |
| 1006 | 2410006 | 0.557664 | -0.45522 | 0.275 | 0.274 | 1 |
| 1034 | Hnrnpdl | 0.7225589 | -0.46081 | 0.319 | 0.254 | 1 |
| 786 | Pak2 | 0.0747492 | -0.46411 | 0.275 | 0.346 | 1 |
| 1068 | G3bp2 | 0.9423228 | -0.46611 | 0.304 | 0.266 | 1 |
| 805 | 7-Sep | 0.0908462 | -0.46646 | 0.217 | 0.288 | 1 |
| 900 | Lbh | 0.2115678 | -0.46955 | 0.261 | 0.307 | 1 |
| 1037 | Mga | 0.761252 | -0.47647 | 0.261 | 0.251 | 1 |
| 825 | Cotl1 | 0.1099189 | -0.4797 | 0.246 | 0.31 | 1 |
| 846 | Rap1b | 0.1385507 | -0.48527 | 0.29 | 0.346 | 1 |
| 996 | Sec62 | 0.5263775 | -0.48642 | 0.275 | 0.283 | 1 |
| 759 | Cnbp | 0.0583195 | -0.48989 | 0.333 | 0.404 | 1 |
| 986 | Pdcd10 | 0.4908386 | -0.4941 | 0.246 | 0.251 | 1 |
| 737 | Arpc2 | 0.0440832 | -0.49663 | 0.319 | 0.434 | 1 |
| 1060 | Dek | 0.8799354 | -0.49851 | 0.275 | 0.227 | 1 |
| 1007 | Rps16 | 0.5643641 | -0.50084 | 0.884 | 0.696 | 1 |
| 768 | Rpl13a | 0.0653808 | -0.50418 | 0.725 | 0.683 | 1 |
| 1026 | Rtf1 | 0.667314 | -0.50468 | 0.275 | 0.214 | 1 |
| 997 | Pafah1b | 0.5286077 | -0.50513 | 0.333 | 0.327 | 1 |
| 947 | Luc7l3 | 0.3348636 | -0.5071 | 0.29 | 0.311 | 1 |
| 767 | Sh3kbp1 | 0.0647727 | -0.51252 | 0.174 | 0.255 | 1 |
| 641 | Nsd3 | 0.0158177 | -0.51517 | 0.29 | 0.408 | 1 |
| 999 | Tmod3 | 0.5330039 | -0.52234 | 0.29 | 0.284 | 1 |
| 811 | Ctcf | 0.0936825 | -0.52262 | 0.188 | 0.256 | 1 |
| 959 | Hnrnpl | 0.3802945 | -0.52528 | 0.275 | 0.288 | 1 |
| 1061 | Nisch | 0.8852856 | -0.53145 | 0.29 | 0.266 | 1 |
| 979 | Atp2b1 | 0.468349 | -0.53359 | 0.246 | 0.278 | 1 |
| 791 | Rplp2 | 0.0800036 | -0.5362 | 0.754 | 0.685 | 1 |
| 578 | Ly6e | 0.0069834 | -0.53841 | 0.478 | 0.631 | 1 |
| 992 | Cop1 | 0.5158955 | -0.53898 | 0.261 | 0.195 | 1 |
| 504 | Slamf6 | 0.0024268 | -0.54178 | 0.087 | 0.251 | 1 |

| | | | | | | |
|---|---|---|---|---|---|---|
| 901 | Cyld | 0.2128035 | -0.54218 | 0.261 | 0.289 | 1 |
| 1072 | Wac | 0.9739314 | -0.5442 | 0.275 | 0.236 | 1 |
| 866 | Srsf5 | 0.1641193 | -0.54432 | 0.362 | 0.403 | 1 |
| 858 | Morf4l1 | 0.1486582 | -0.54488 | 0.348 | 0.403 | 1 |
| 989 | Top1 | 0.5058543 | -0.54808 | 0.29 | 0.302 | 1 |
| 730 | Tmsb10 | 0.0423418 | -0.54928 | 0.536 | 0.595 | 1 |
| 970 | Wdr26 | 0.4228206 | -0.5527 | 0.261 | 0.266 | 1 |
| 752 | Ptp4a2 | 0.0545878 | -0.55746 | 0.377 | 0.464 | 1 |
| 1059 | Fbl | 0.8776767 | -0.55766 | 0.261 | 0.231 | 1 |
| 869 | Hnrnph1 | 0.1684997 | -0.55904 | 0.333 | 0.38 | 1 |
| 928 | Grcc10 | 0.2740622 | -0.56016 | 0.333 | 0.34 | 1 |
| 904 | Mkln1 | 0.2216659 | -0.56049 | 0.261 | 0.303 | 1 |
| 1003 | Acin1 | 0.5473184 | -0.56568 | 0.362 | 0.35 | 1 |
| 908 | Strbp | 0.2260221 | -0.56687 | 0.261 | 0.172 | 1 |
| 868 | Hectd1 | 0.168138 | -0.5757 | 0.246 | 0.304 | 1 |
| 884 | Ep300 | 0.1892794 | -0.57955 | 0.246 | 0.288 | 1 |
| 976 | Tmed9 | 0.4616538 | -0.58234 | 0.246 | 0.262 | 1 |
| 859 | Hnrnpr | 0.1504887 | -0.5829 | 0.232 | 0.283 | 1 |
| 713 | Sh3bgrl3 | 0.0353502 | -0.58407 | 0.348 | 0.452 | 1 |
| 857 | Psme1 | 0.147272 | -0.58942 | 0.319 | 0.354 | 1 |
| 930 | Leprotl1 | 0.2832207 | -0.592 | 0.232 | 0.254 | 1 |
| 885 | Ppp4r3b | 0.1901128 | -0.59339 | 0.217 | 0.259 | 1 |
| 828 | Fau | 0.1112554 | -0.59961 | 0.841 | 0.72 | 1 |
| 829 | Arl6ip1 | 0.1115891 | -0.6011 | 0.261 | 0.306 | 1 |
| 852 | Rbbp6 | 0.142818 | -0.60362 | 0.304 | 0.352 | 1 |
| 1020 | Csnk2a1 | 0.6285737 | -0.60429 | 0.29 | 0.276 | 1 |
| 667 | Gmfg | 0.0210515 | -0.6054 | 0.203 | 0.312 | 1 |
| 1050 | Capza2 | 0.8263446 | -0.6055 | 0.29 | 0.262 | 1 |
| 898 | Kpna4 | 0.2099451 | -0.6213 | 0.232 | 0.268 | 1 |
| 963 | Picalm | 0.395274 | -0.62228 | 0.246 | 0.264 | 1 |
| 763 | Chd3 | 0.0626294 | -0.62409 | 0.29 | 0.353 | 1 |
| 870 | Hnrnpm | 0.1701106 | -0.626 | 0.261 | 0.302 | 1 |
| 871 | Cacybp | 0.1714429 | -0.63013 | 0.232 | 0.265 | 1 |
| 771 | AW1120 | 0.0691131 | -0.63154 | 0.174 | 0.259 | 1 |
| 932 | Pnrc1 | 0.2922383 | -0.63179 | 0.275 | 0.293 | 1 |
| 867 | Scaf11 | 0.1641546 | -0.63653 | 0.304 | 0.335 | 1 |
| 533 | Ccr7 | 0.0036375 | -0.63808 | 0.101 | 0.261 | 1 |
| 931 | Pbrm1 | 0.2839904 | -0.64093 | 0.261 | 0.282 | 1 |
| 1051 | Msi2 | 0.8265537 | -0.64334 | 0.275 | 0.231 | 1 |
| 824 | Kmt2c | 0.1087135 | -0.64501 | 0.246 | 0.298 | 1 |
| 974 | Pkn1 | 0.4468081 | -0.64838 | 0.261 | 0.26 | 1 |
| 1041 | Syncrip | 0.7743048 | -0.65111 | 0.275 | 0.256 | 1 |
| 550 | Hcst | 0.0049027 | -0.65185 | 0.116 | 0.26 | 1 |

| | | | | | | |
|---|---|---|---|---|---|---|
| 813 | Herc1 | 0.0954697 | -0.6555 | 0.232 | 0.287 | 1 |
| 756 | Ncor1 | 0.0556151 | -0.65566 | 0.391 | 0.442 | 1 |
| 916 | Mier1 | 0.2425589 | -0.65708 | 0.29 | 0.311 | 1 |
| 882 | Ubap2l | 0.186359 | -0.65768 | 0.29 | 0.318 | 1 |
| 1067 | Ppp3ca | 0.941128 | -0.66069 | 0.319 | 0.282 | 1 |
| 681 | Chordc1 | 0.0253867 | -0.66929 | 0.203 | 0.294 | 1 |
| 961 | Jmjd1c | 0.3814994 | -0.66966 | 0.304 | 0.312 | 1 |
| 818 | Tln1 | 0.1016059 | -0.67894 | 0.246 | 0.315 | 1 |
| 1005 | Ddx50 | 0.5531818 | -0.68335 | 0.261 | 0.255 | 1 |
| 965 | Gdi2 | 0.3984456 | -0.68347 | 0.348 | 0.349 | 1 |
| 638 | Rpl9 | 0.0152512 | -0.68358 | 0.623 | 0.625 | 1 |
| 841 | Rock1 | 0.1359391 | -0.68827 | 0.348 | 0.381 | 1 |
| 808 | Arglu1 | 0.0930778 | -0.68998 | 0.304 | 0.355 | 1 |
| 987 | Klhl24 | 0.4961446 | -0.69123 | 0.261 | 0.264 | 1 |
| 878 | Sbno1 | 0.1833162 | -0.69323 | 0.275 | 0.303 | 1 |
| 792 | Ewsr1 | 0.0819502 | -0.69516 | 0.261 | 0.318 | 1 |
| 850 | Ube2d2a | 0.1399106 | -0.69809 | 0.319 | 0.368 | 1 |
| 754 | Mat2a | 0.0547451 | -0.70114 | 0.304 | 0.382 | 1 |
| 540 | Srsf3 | 0.0042199 | -0.70296 | 0.377 | 0.479 | 1 |
| 802 | Emp3 | 0.0884077 | -0.70693 | 0.246 | 0.308 | 1 |
| 418 | Rps21 | 0.0003726 | -0.70843 | 0.739 | 0.767 | 1 |
| 952 | Kmt5b | 0.3393069 | -0.71286 | 0.232 | 0.252 | 1 |
| 879 | Tnrc6c | 0.1847371 | -0.7134 | 0.29 | 0.32 | 1 |
| 777 | Rab14 | 0.0710824 | -0.71443 | 0.304 | 0.373 | 1 |
| 876 | Pten | 0.1802553 | -0.71566 | 0.261 | 0.301 | 1 |
| 428 | Dnaja1 | 0.0004514 | -0.71569 | 0.406 | 0.569 | 1 |
| 687 | Rsrc2 | 0.0267326 | -0.71647 | 0.29 | 0.38 | 1 |
| 922 | Ugcg | 0.2586515 | -0.71693 | 0.246 | 0.271 | 1 |
| 1056 | Mark3 | 0.8703373 | -0.71818 | 0.261 | 0.217 | 1 |
| 489 | Add3 | 0.0018192 | -0.72299 | 0.188 | 0.349 | 1 |
| 953 | Xrn2 | 0.340703 | -0.72366 | 0.232 | 0.25 | 1 |
| 775 | Prpf4b | 0.0705936 | -0.72461 | 0.333 | 0.386 | 1 |
| 548 | Emb | 0.0047556 | -0.72489 | 0.13 | 0.266 | 1 |
| 660 | Hsp90aa | 0.0198537 | -0.72638 | 0.377 | 0.488 | 1 |
| 788 | Tbc1d1 | 0.0764048 | -0.72819 | 0.188 | 0.256 | 1 |
| 517 | Hnrnpa3 | 0.0029301 | -0.72994 | 0.377 | 0.532 | 1 |
| 684 | Pde7a | 0.0261932 | -0.73123 | 0.217 | 0.308 | 1 |
| 784 | Esyt1 | 0.0732469 | -0.73899 | 0.188 | 0.261 | 1 |
| 731 | Rps24 | 0.0425835 | -0.74047 | 0.812 | 0.771 | 1 |
| 523 | Cd47 | 0.0030441 | -0.74491 | 0.333 | 0.499 | 1 |
| 583 | Arid1a | 0.0073059 | -0.74552 | 0.246 | 0.377 | 1 |
| 718 | Sptbn1 | 0.0386658 | -0.74634 | 0.261 | 0.361 | 1 |
| 682 | Ccnd2 | 0.025413 | -0.74724 | 0.217 | 0.327 | 1 |

| | | | | | | |
|---|---|---|---|---|---|---|
| 849 | Cnot1 | 0.139816 | -0.75149 | 0.232 | 0.278 | 1 |
| 962 | Celf1 | 0.3887904 | -0.75167 | 0.29 | 0.297 | 1 |
| 589 | Bptf | 0.007941 | -0.75508 | 0.319 | 0.432 | 1 |
| 917 | Ppp2r5a | 0.2437623 | -0.75672 | 0.246 | 0.268 | 1 |
| 758 | Pum2 | 0.0570843 | -0.76152 | 0.246 | 0.321 | 1 |
| 623 | Hnrnpu | 0.0123345 | -0.76351 | 0.42 | 0.52 | 1 |
| 374 | Myh9 | 0.0001436 | -0.76352 | 0.333 | 0.587 | 1 |
| 1028 | Kdm5a | 0.6806456 | -0.76634 | 0.261 | 0.243 | 1 |
| 613 | Clk1 | 0.010637 | -0.77784 | 0.348 | 0.45 | 1 |
| 848 | Ankrd17 | 0.1396734 | -0.78598 | 0.304 | 0.354 | 1 |
| 425 | Srrm2 | 0.0004311 | -0.78867 | 0.435 | 0.591 | 1 |
| 854 | Tia1 | 0.1440938 | -0.79247 | 0.203 | 0.254 | 1 |
| 392 | Hnrnpa2 | 0.0002219 | -0.79252 | 0.507 | 0.683 | 1 |
| 822 | Top2b | 0.107626 | -0.79349 | 0.275 | 0.321 | 1 |
| 740 | Rps15a | 0.0473076 | -0.79817 | 0.783 | 0.682 | 1 |
| 659 | Zfp638 | 0.0197902 | -0.79853 | 0.232 | 0.332 | 1 |
| 920 | Usp34 | 0.2532717 | -0.80053 | 0.304 | 0.317 | 1 |
| 700 | Nufip2 | 0.0300463 | -0.80296 | 0.203 | 0.297 | 1 |
| 746 | Stk4 | 0.0510494 | -0.80597 | 0.217 | 0.289 | 1 |
| 656 | Eif3e | 0.0192981 | -0.80946 | 0.275 | 0.353 | 1 |
| 668 | Ppp1r18 | 0.0210979 | -0.81078 | 0.159 | 0.255 | 1 |
| 921 | Kdm2a | 0.2554714 | -0.8111 | 0.29 | 0.307 | 1 |
| 430 | Il21r | 0.000454 | -0.81198 | 0.072 | 0.26 | 1 |
| 349 | Cytip | 8.28E-05 | -0.81283 | 0.145 | 0.362 | 1 |
| 821 | Rsbn1l | 0.1064687 | -0.81462 | 0.261 | 0.312 | 1 |
| 863 | Cnot6l | 0.15895 | -0.81754 | 0.261 | 0.296 | 1 |
| 645 | Cyba | 0.0165263 | -0.82086 | 0.261 | 0.377 | 1 |
| 914 | Ddx24 | 0.2407345 | -0.82407 | 0.232 | 0.26 | 1 |
| 522 | Rps13 | 0.0030375 | -0.82479 | 0.609 | 0.652 | 1 |
| 534 | Stk38 | 0.0036584 | -0.82575 | 0.232 | 0.361 | 1 |
| 487 | Tsc22d3 | 0.0017346 | -0.83998 | 0.232 | 0.387 | 1 |
| 909 | Setd2 | 0.2260844 | -0.84084 | 0.246 | 0.273 | 1 |
| 694 | Tacc1 | 0.0282821 | -0.8424 | 0.159 | 0.259 | 1 |
| 938 | Gramd1a | 0.3178094 | -0.84431 | 0.246 | 0.257 | 1 |
| 743 | Zc3hav1 | 0.0494999 | -0.84582 | 0.261 | 0.33 | 1 |
| 729 | Gpbp1 | 0.0423306 | -0.84719 | 0.232 | 0.306 | 1 |
| 617 | Nipbl | 0.011619 | -0.8512 | 0.29 | 0.391 | 1 |
| 902 | Sltm | 0.2151804 | -0.85656 | 0.304 | 0.32 | 1 |
| 799 | 7-Mar | 0.0860163 | -0.85736 | 0.246 | 0.304 | 1 |
| 624 | Tra2b | 0.0124337 | -0.85935 | 0.319 | 0.415 | 1 |
| 689 | Elf1 | 0.0271814 | -0.8625 | 0.246 | 0.334 | 1 |
| 701 | Lbr | 0.0303581 | -0.86329 | 0.188 | 0.275 | 1 |
| 941 | Ubn2 | 0.3210618 | -0.86802 | 0.246 | 0.27 | 1 |

| | | | | | | |
|---|---|---|---|---|---|---|
| 760 | Hsph1 | 0.0592122 | -0.86943 | 0.188 | 0.259 | 1 |
| 929 | Mtdh | 0.2800754 | -0.87045 | 0.29 | 0.303 | 1 |
| 447 | Eif4a2 | 0.0007163 | -0.87819 | 0.304 | 0.475 | 1 |
| 794 | Rb1cc1 | 0.0834345 | -0.88324 | 0.217 | 0.278 | 1 |
| 732 | Grk6 | 0.0427179 | -0.886 | 0.174 | 0.257 | 1 |
| 331 | Prrc2c | 4.80E-05 | -0.88643 | 0.377 | 0.571 | 1 |
| 546 | Actr2 | 0.0046328 | -0.88862 | 0.319 | 0.433 | 1 |
| 503 | Snrnp70 | 0.0024214 | -0.88938 | 0.319 | 0.464 | 1 |
| 618 | Ash1l | 0.0117193 | -0.89224 | 0.333 | 0.432 | 1 |
| 515 | Sfpq | 0.0028374 | -0.89423 | 0.362 | 0.502 | 1 |
| 795 | B4galnt1 | 0.0838893 | -0.89484 | 0.217 | 0.275 | 1 |
| 856 | Crebbp | 0.1470824 | -0.89499 | 0.261 | 0.299 | 1 |
| 552 | Tpr | 0.0049808 | -0.89893 | 0.304 | 0.429 | 1 |
| 651 | Ogt | 0.0174405 | -0.90348 | 0.333 | 0.419 | 1 |
| 629 | Nfatc3 | 0.0140899 | -0.90362 | 0.174 | 0.284 | 1 |
| 691 | Klf13 | 0.0273883 | -0.91017 | 0.29 | 0.367 | 1 |
| 915 | Gls | 0.2413967 | -0.91201 | 0.275 | 0.292 | 1 |
| 609 | Ptpn22 | 0.0104396 | -0.91319 | 0.13 | 0.255 | 1 |
| 773 | Sp3 | 0.069507 | -0.91385 | 0.217 | 0.275 | 1 |
| 728 | Srpk2 | 0.0422094 | -0.91751 | 0.203 | 0.289 | 1 |
| 704 | Pds5a | 0.0309725 | -0.92028 | 0.232 | 0.324 | 1 |
| 839 | Filip1l | 0.1292865 | -0.92168 | 0.246 | 0.294 | 1 |
| 639 | 1-Sep | 0.015257 | -0.9254 | 0.174 | 0.279 | 1 |
| 327 | Tbc1d10 | 4.43E-05 | -0.93207 | 0.043 | 0.271 | 0.92754 |
| 502 | Phip | 0.0023822 | -0.93207 | 0.348 | 0.483 | 1 |
| 564 | Fam49b | 0.0057475 | -0.93317 | 0.246 | 0.377 | 1 |
| 543 | Pik3ip1 | 0.0045905 | -0.9365 | 0.13 | 0.273 | 1 |
| 614 | Kmt2a | 0.0107202 | -0.9421 | 0.319 | 0.418 | 1 |
| 695 | Rictor | 0.0284561 | -0.94376 | 0.174 | 0.262 | 1 |
| 420 | Sp110 | 0.0003863 | -0.94533 | 0.072 | 0.259 | 1 |
| 685 | Utrn | 0.0266379 | -0.94932 | 0.232 | 0.325 | 1 |
| 519 | Evl | 0.0029563 | -0.95159 | 0.13 | 0.282 | 1 |
| 679 | Elmo1 | 0.0243502 | -0.95322 | 0.174 | 0.276 | 1 |
| 714 | Cbl | 0.0358441 | -0.95489 | 0.232 | 0.31 | 1 |
| 762 | Ube3a | 0.0619649 | -0.95509 | 0.246 | 0.308 | 1 |
| 408 | Prpf38b | 0.0003151 | -0.9553 | 0.261 | 0.436 | 1 |
| 405 | Fam169b | 0.0003006 | -0.95687 | 0.058 | 0.254 | 1 |
| 394 | Tcf7 | 0.0002433 | -0.95871 | 0.087 | 0.289 | 1 |
| 471 | Akna | 0.0012349 | -0.96139 | 0.101 | 0.265 | 1 |
| 905 | Tut7 | 0.2218268 | -0.96568 | 0.304 | 0.316 | 1 |
| 304 | Msn | 2.20E-05 | -0.96589 | 0.261 | 0.515 | 0.46045 |
| 529 | Apbb1ip | 0.0034206 | -0.96676 | 0.159 | 0.302 | 1 |
| 889 | Setd5 | 0.1984412 | -0.968 | 0.246 | 0.276 | 1 |

| | | | | | | |
|---|---|---|---|---|---|---|
| 599 | Kmt2e | 0.0089437 | -0.97279 | 0.304 | 0.411 | 1 |
| 837 | AC14909 | 0.1207952 | -0.97512 | 0.246 | 0.293 | 1 |
| 757 | Matr3 | 0.0556187 | -0.9783 | 0.304 | 0.358 | 1 |
| 750 | Zbtb20 | 0.0532558 | -0.98765 | 0.246 | 0.33 | 1 |
| 506 | Ddx6 | 0.0024781 | -0.98794 | 0.333 | 0.461 | 1 |
| 787 | Ago2 | 0.075811 | -0.98936 | 0.217 | 0.274 | 1 |
| 807 | Pnn | 0.0924376 | -0.9899 | 0.246 | 0.299 | 1 |
| 680 | Gabpb2 | 0.0246506 | -0.99096 | 0.159 | 0.252 | 1 |
| 778 | Zfp644 | 0.0713647 | -0.9916 | 0.232 | 0.288 | 1 |
| 569 | Slc38a1 | 0.0062282 | -0.99181 | 0.203 | 0.335 | 1 |
| 708 | Serinc3 | 0.0334024 | -0.99347 | 0.406 | 0.462 | 1 |
| 664 | Tsc22d4 | 0.0205979 | -0.99798 | 0.275 | 0.348 | 1 |
| 711 | Srpk1 | 0.0343382 | -0.9981 | 0.246 | 0.318 | 1 |
| 545 | Arhgef18 | 0.0046131 | -0.99955 | 0.246 | 0.377 | 1 |
| 381 | Skap1 | 0.0001621 | -1.00288 | 0.058 | 0.261 | 1 |
| 620 | Smad7 | 0.0118472 | -1.00342 | 0.246 | 0.363 | 1 |
| 596 | 4932438 | 0.008373 | -1.00421 | 0.232 | 0.344 | 1 |
| 154 | Ddx5 | 2.31E-08 | -1.00736 | 0.493 | 0.769 | 0.00048 |
| 346 | Ablim1 | 7.60E-05 | -1.008 | 0.304 | 0.504 | 1 |
| 357 | Lat | 0.0001004 | -1.00861 | 0.058 | 0.275 | 1 |
| 826 | Zdhhc20 | 0.1106027 | -1.00908 | 0.232 | 0.276 | 1 |
| 810 | Cpsf6 | 0.0936563 | -1.0101 | 0.217 | 0.27 | 1 |
| 688 | Tasor | 0.0271193 | -1.01129 | 0.217 | 0.31 | 1 |
| 625 | Itsn2 | 0.0126114 | -1.01725 | 0.275 | 0.377 | 1 |
| 798 | Man2b1 | 0.0852102 | -1.02368 | 0.188 | 0.25 | 1 |
| 823 | Resf1 | 0.1081086 | -1.02388 | 0.217 | 0.266 | 1 |
| 612 | Ccdc88c | 0.0105933 | -1.02728 | 0.217 | 0.326 | 1 |
| 588 | Phf20l1 | 0.0078797 | -1.0286 | 0.217 | 0.324 | 1 |
| 592 | Saraf | 0.0081136 | -1.02946 | 0.174 | 0.288 | 1 |
| 790 | Irf2 | 0.079017 | -1.0331 | 0.261 | 0.307 | 1 |
| 872 | Camk2d | 0.1745636 | -1.03588 | 0.217 | 0.257 | 1 |
| 477 | Man1a | 0.001359 | -1.03673 | 0.203 | 0.363 | 1 |
| 945 | Bod1l | 0.3295445 | -1.04426 | 0.275 | 0.274 | 1 |
| 256 | Rsrp1 | 3.93E-06 | -1.04598 | 0.377 | 0.596 | 0.08222 |
| 586 | Ptpn6 | 0.0074721 | -1.04599 | 0.145 | 0.269 | 1 |
| 372 | Pik3cd | 0.0001425 | -1.04736 | 0.058 | 0.257 | 1 |
| 587 | Sesn3 | 0.0078641 | -1.05223 | 0.145 | 0.269 | 1 |
| 705 | Creb1 | 0.0313299 | -1.05247 | 0.203 | 0.288 | 1 |
| 329 | Actr3 | 4.60E-05 | -1.05312 | 0.304 | 0.515 | 0.96424 |
| 591 | Srsf11 | 0.00804 | -1.05885 | 0.29 | 0.389 | 1 |
| 361 | Thy1 | 0.000102 | -1.05932 | 0.043 | 0.254 | 1 |
| 497 | Ikbkb | 0.0021498 | -1.06144 | 0.246 | 0.386 | 1 |
| 690 | Safb2 | 0.0272683 | -1.06192 | 0.246 | 0.327 | 1 |

| | | | | | | |
|---|---|---|---|---|---|---|
| 370 | Dgka | 0.0001378 | -1.06462 | 0.13 | 0.348 | 1 |
| 383 | Cxcr4 | 0.0001642 | -1.06666 | 0.145 | 0.369 | 1 |
| 567 | Rps27rt | 0.0060227 | -1.07414 | 0.217 | 0.325 | 1 |
| 393 | Lsp1 | 0.0002248 | -1.08434 | 0.145 | 0.345 | 1 |
| 676 | Phf3 | 0.0235285 | -1.09251 | 0.217 | 0.303 | 1 |
| 363 | Iqgap1 | 0.0001062 | -1.09555 | 0.304 | 0.543 | 1 |
| 525 | Cela1 | 0.0032429 | -1.1024 | 0.406 | 0.619 | 1 |
| 315 | Cd2 | 3.08E-05 | -1.10399 | 0.072 | 0.303 | 0.64442 |
| 313 | Gimap8 | 2.85E-05 | -1.10586 | 0.043 | 0.274 | 0.59667 |
| 739 | Zc3h7a | 0.046863 | -1.11743 | 0.174 | 0.25 | 1 |
| 683 | Cflar | 0.02608 | -1.11908 | 0.159 | 0.25 | 1 |
| 640 | Grk2 | 0.0154722 | -1.12019 | 0.203 | 0.296 | 1 |
| 665 | Foxn3 | 0.0207811 | -1.12829 | 0.246 | 0.331 | 1 |
| 654 | Nktr | 0.0182885 | -1.13028 | 0.29 | 0.372 | 1 |
| 449 | Lnpep | 0.0007349 | -1.13031 | 0.203 | 0.366 | 1 |
| 350 | Ankrd11 | 8.28E-05 | -1.13117 | 0.377 | 0.566 | 1 |
| 774 | Ranbp2 | 0.0704286 | -1.13185 | 0.217 | 0.271 | 1 |
| 633 | Chd2 | 0.0147727 | -1.13207 | 0.261 | 0.355 | 1 |
| 348 | Ptbp3 | 8.18E-05 | -1.13748 | 0.261 | 0.473 | 1 |
| 888 | Dmxl1 | 0.1918672 | -1.13952 | 0.232 | 0.265 | 1 |
| 566 | N4bp2l2 | 0.0059932 | -1.13995 | 0.217 | 0.336 | 1 |
| 611 | Itch | 0.0105246 | -1.14249 | 0.188 | 0.296 | 1 |
| 286 | Crlf3 | 1.16E-05 | -1.14426 | 0.203 | 0.443 | 0.24391 |
| 462 | Nsd1 | 0.0009806 | -1.14749 | 0.217 | 0.381 | 1 |
| 541 | Ptpn18 | 0.0042222 | -1.15807 | 0.275 | 0.38 | 1 |
| 263 | Grap2 | 4.88E-06 | -1.1603 | 0.058 | 0.327 | 0.10221 |
| 275 | Limd2 | 7.40E-06 | -1.16179 | 0.174 | 0.438 | 0.15495 |
| 382 | Fxyd5 | 0.0001628 | -1.16247 | 0.217 | 0.409 | 1 |
| 457 | Rps27 | 0.0008907 | -1.16434 | 0.739 | 0.708 | 1 |
| 179 | Ube2d3 | 1.73E-07 | -1.16757 | 0.449 | 0.699 | 0.00363 |
| 367 | Selplg | 0.0001309 | -1.1723 | 0.101 | 0.308 | 1 |
| 509 | Inpp4b | 0.0025925 | -1.17951 | 0.116 | 0.264 | 1 |
| 528 | Tap2 | 0.0033943 | -1.18174 | 0.116 | 0.251 | 1 |
| 332 | Atrx | 4.85E-05 | -1.18587 | 0.275 | 0.47 | 1 |
| 493 | Bcl2 | 0.0020669 | -1.18593 | 0.174 | 0.33 | 1 |
| 450 | Slfn2 | 0.0007481 | -1.18868 | 0.174 | 0.343 | 1 |
| 604 | Tet3 | 0.00983 | -1.19027 | 0.174 | 0.284 | 1 |
| 532 | Atf7ip | 0.0036283 | -1.19147 | 0.261 | 0.394 | 1 |
| 675 | Tecpr1 | 0.0228389 | -1.19349 | 0.174 | 0.261 | 1 |
| 419 | Gm8369 | 0.0003831 | -1.19474 | 0.087 | 0.276 | 1 |
| 312 | Gm2682 | 2.79E-05 | -1.20424 | 0.043 | 0.271 | 0.58451 |
| 585 | Nfkb1 | 0.0074363 | -1.20709 | 0.188 | 0.302 | 1 |
| 288 | Pnisr | 1.28E-05 | -1.21304 | 0.261 | 0.488 | 0.26743 |

| | | | | | | |
|---|---|---|---|---|---|---|
| 342 | Ifi209 | 6.87E-05 | -1.21744 | 0.087 | 0.308 | 1 |
| 323 | Sf3b1 | 3.82E-05 | -1.22261 | 0.319 | 0.521 | 0.80012 |
| 478 | Itpr2 | 0.0013704 | -1.22301 | 0.159 | 0.31 | 1 |
| 717 | Pan3 | 0.0382685 | -1.22464 | 0.246 | 0.312 | 1 |
| 460 | Smc4 | 0.0009542 | -1.22552 | 0.159 | 0.324 | 1 |
| 195 | Gimap3 | 4.79E-07 | -1.23105 | 0.072 | 0.389 | 0.01002 |
| 222 | Ptprcap | 1.33E-06 | -1.23873 | 0.145 | 0.437 | 0.02784 |
| 661 | Slc44a2 | 0.0199076 | -1.24147 | 0.203 | 0.298 | 1 |
| 442 | Sipa1 | 0.000681 | -1.24322 | 0.087 | 0.256 | 1 |
| 455 | Clec2d | 0.0008482 | -1.24361 | 0.217 | 0.38 | 1 |
| 424 | Itpkb | 0.0004186 | -1.2445 | 0.13 | 0.313 | 1 |
| 387 | Psmb8 | 0.0001854 | -1.24803 | 0.188 | 0.372 | 1 |
| 472 | Ythdc1 | 0.0012585 | -1.24867 | 0.232 | 0.381 | 1 |
| 678 | Tcf12 | 0.0237117 | -1.25603 | 0.203 | 0.287 | 1 |
| 298 | Cd3g | 1.76E-05 | -1.25719 | 0.072 | 0.327 | 0.36828 |
| 423 | Zcchc7 | 0.0004069 | -1.25923 | 0.275 | 0.448 | 1 |
| 322 | Rasal3 | 3.79E-05 | -1.26425 | 0.043 | 0.264 | 0.79327 |
| 429 | Bclaf1 | 0.0004515 | -1.26811 | 0.304 | 0.455 | 1 |
| 328 | Cd79b | 4.48E-05 | -1.27639 | 0.087 | 0.32 | 0.93817 |
| 631 | Tapbp | 0.0143625 | -1.27781 | 0.217 | 0.306 | 1 |
| 570 | Samd9l | 0.0062728 | -1.28094 | 0.246 | 0.352 | 1 |
| 536 | Adgre5 | 0.0037333 | -1.28119 | 0.174 | 0.297 | 1 |
| 467 | Prkd2 | 0.0011924 | -1.28457 | 0.116 | 0.273 | 1 |
| 301 | Rasgrp2 | 1.84E-05 | -1.28528 | 0.116 | 0.368 | 0.38625 |
| 755 | Brwd1 | 0.0552351 | -1.29017 | 0.217 | 0.279 | 1 |
| 269 | Lcp1 | 6.05E-06 | -1.29425 | 0.246 | 0.492 | 0.12671 |
| 725 | Sorl1 | 0.0415796 | -1.29536 | 0.188 | 0.264 | 1 |
| 427 | Krit1 | 0.0004494 | -1.29655 | 0.217 | 0.391 | 1 |
| 765 | Xist | 0.064442 | -1.30247 | 0.333 | 0.401 | 1 |
| 378 | Bcl11b | 0.0001512 | -1.30297 | 0.116 | 0.322 | 1 |
| 321 | Tra2a | 3.53E-05 | -1.3058 | 0.246 | 0.443 | 0.73898 |
| 411 | Tut4 | 0.0003291 | -1.30757 | 0.261 | 0.42 | 1 |
| 330 | Cd69 | 4.77E-05 | -1.3107 | 0.101 | 0.335 | 0.99848 |
| 282 | Cd27 | 9.83E-06 | -1.31661 | 0.029 | 0.264 | 0.20595 |
| 390 | Fus | 0.000204 | -1.32114 | 0.348 | 0.504 | 1 |
| 473 | Rnf167 | 0.0013006 | -1.32476 | 0.116 | 0.266 | 1 |
| 208 | Son | 6.35E-07 | -1.32479 | 0.348 | 0.606 | 0.01329 |
| 454 | Gm2055 | 0.0008082 | -1.32688 | 0.159 | 0.327 | 1 |
| 188 | H2-Q7 | 3.25E-07 | -1.32922 | 0.159 | 0.496 | 0.0068 |
| 396 | Wnk1 | 0.0002566 | -1.32929 | 0.29 | 0.464 | 1 |
| 250 | Rac2 | 3.31E-06 | -1.33293 | 0.188 | 0.452 | 0.06938 |
| 186 | Macf1 | 3.07E-07 | -1.33451 | 0.391 | 0.638 | 0.00643 |
| 283 | Trim12a | 1.03E-05 | -1.3422 | 0.13 | 0.371 | 0.2158 |

| | | | | | | |
|---|---|---|---|---|---|---|
| 486 | Ccnl1 | 0.0017114 | -1.36174 | 0.232 | 0.359 | 1 |
| 339 | Ttc14 | 6.03E-05 | -1.39434 | 0.275 | 0.457 | 1 |
| 470 | Birc6 | 0.0012286 | -1.40501 | 0.261 | 0.389 | 1 |
| 225 | Zfp36l2 | 1.45E-06 | -1.40638 | 0.29 | 0.539 | 0.03036 |
| 520 | Smg1 | 0.0029652 | -1.4067 | 0.217 | 0.343 | 1 |
| 413 | Atp11b | 0.0003356 | -1.4069 | 0.174 | 0.346 | 1 |
| 498 | Zfp292 | 0.0022256 | -1.40847 | 0.232 | 0.349 | 1 |
| 204 | S1pr1 | 5.77E-07 | -1.40856 | 0.072 | 0.376 | 0.01209 |
| 605 | Mdm4 | 0.0098432 | -1.41007 | 0.188 | 0.287 | 1 |
| 242 | Tespa1 | 2.38E-06 | -1.41393 | 0.043 | 0.315 | 0.04992 |
| 151 | B2m | 1.44E-08 | -1.4182 | 0.435 | 0.671 | 0.0003 |
| 210 | Rapgef6 | 7.90E-07 | -1.41878 | 0.232 | 0.503 | 0.01654 |
| 326 | Cd84 | 4.41E-05 | -1.42528 | 0.101 | 0.327 | 0.92301 |
| 244 | Itgb7 | 2.47E-06 | -1.43319 | 0.058 | 0.326 | 0.05163 |
| 379 | Ankrd12 | 0.0001513 | -1.43377 | 0.174 | 0.367 | 1 |
| 373 | 1810026 | 0.0001436 | -1.44252 | 0.145 | 0.336 | 1 |
| 230 | Ms4a6b | 1.98E-06 | -1.44709 | 0.116 | 0.401 | 0.04149 |
| 259 | Srsf7 | 4.31E-06 | -1.45666 | 0.232 | 0.464 | 0.09026 |
| 285 | Apobec3 | 1.16E-05 | -1.45811 | 0.275 | 0.489 | 0.24242 |
| 163 | Sell | 5.43E-08 | -1.45982 | 0.145 | 0.483 | 0.00114 |
| 254 | Cd3d | 3.64E-06 | -1.45996 | 0.072 | 0.35 | 0.07616 |
| 207 | Luc7l2 | 6.30E-07 | -1.46789 | 0.319 | 0.576 | 0.0132 |
| 113 | Satb1 | 2.68E-10 | -1.47196 | 0.159 | 0.564 | 5.60E-06 |
| 214 | Cd3e | 8.45E-07 | -1.47393 | 0.058 | 0.354 | 0.0177 |
| 432 | Pag1 | 0.0004992 | -1.47607 | 0.087 | 0.262 | 1 |
| 444 | Prkcb | 0.0007066 | -1.47755 | 0.087 | 0.251 | 1 |
| 320 | Fyb | 3.51E-05 | -1.48091 | 0.13 | 0.357 | 0.73512 |
| 306 | Traf3ip3 | 2.21E-05 | -1.48379 | 0.087 | 0.317 | 0.46272 |
| 280 | Il2rg | 9.48E-06 | -1.49017 | 0.13 | 0.376 | 0.19858 |
| 307 | Peli1 | 2.30E-05 | -1.52711 | 0.188 | 0.404 | 0.48201 |
| 316 | H2-T23 | 3.24E-05 | -1.52797 | 0.232 | 0.424 | 0.67916 |
| 593 | Slc12a6 | 0.0081935 | -1.53294 | 0.145 | 0.255 | 1 |
| 202 | Lck | 5.55E-07 | -1.5374 | 0.043 | 0.34 | 0.01163 |
| 107 | Rbm39 | 6.80E-11 | -1.54365 | 0.406 | 0.72 | 1.42E-06 |
| 165 | Ltb | 6.94E-08 | -1.54387 | 0.203 | 0.511 | 0.00145 |
| 184 | Arhgdib | 2.84E-07 | -1.54695 | 0.188 | 0.483 | 0.00594 |
| 295 | Tnrc6b | 1.65E-05 | -1.55623 | 0.232 | 0.433 | 0.34514 |
| 173 | Arhgap3 | 1.18E-07 | -1.56586 | 0.116 | 0.436 | 0.00247 |
| 155 | Fubp1 | 2.63E-08 | -1.56628 | 0.275 | 0.585 | 0.00055 |
| 398 | Slc38a2 | 0.0002665 | -1.5698 | 0.232 | 0.409 | 1 |
| 177 | Akap13 | 1.49E-07 | -1.57002 | 0.29 | 0.561 | 0.00311 |
| 187 | Ikzf1 | 3.12E-07 | -1.57015 | 0.101 | 0.405 | 0.00654 |
| 376 | St8sia4 | 0.0001501 | -1.58054 | 0.087 | 0.29 | 1 |

| | | | | | | |
|---|---|---|---|---|---|---|
| 253 | Celf2 | 3.56E-06 | -1.59106 | 0.246 | 0.474 | 0.07459 |
| 235 | Rbm25 | 2.10E-06 | -1.59168 | 0.29 | 0.529 | 0.04405 |
| 422 | Parp14 | 0.0003956 | -1.59746 | 0.087 | 0.259 | 1 |
| 200 | Gimap4 | 5.23E-07 | -1.60718 | 0.072 | 0.364 | 0.01096 |
| 309 | Ikzf3 | 2.45E-05 | -1.60783 | 0.029 | 0.254 | 0.51254 |
| 318 | Ccnd3 | 3.33E-05 | -1.61427 | 0.217 | 0.417 | 0.69807 |
| 205 | Trac | 5.93E-07 | -1.6169 | 0.058 | 0.345 | 0.01241 |
| 90 | H2-D1 | 1.20E-13 | -1.61726 | 0.42 | 0.773 | 2.50E-09 |
| 516 | Zfp318 | 0.0028631 | -1.61745 | 0.116 | 0.261 | 1 |
| 438 | Ifi27l2a | 0.000605 | -1.63003 | 0.145 | 0.324 | 1 |
| 343 | Smchd1 | 7.17E-05 | -1.63524 | 0.232 | 0.409 | 1 |
| 174 | Cd53 | 1.39E-07 | -1.64268 | 0.188 | 0.484 | 0.00291 |
| 180 | Mndal | 1.83E-07 | -1.64468 | 0.159 | 0.471 | 0.00383 |
| 193 | Cmah | 4.15E-07 | -1.64477 | 0.072 | 0.369 | 0.0087 |
| 268 | Ssh2 | 5.97E-06 | -1.64925 | 0.188 | 0.424 | 0.12499 |
| 192 | Mycbp2 | 4.04E-07 | -1.65239 | 0.304 | 0.54 | 0.00846 |
| 215 | Rbm5 | 9.16E-07 | -1.65373 | 0.232 | 0.48 | 0.01919 |
| 196 | Gm2674 | 4.81E-07 | -1.66668 | 0.101 | 0.405 | 0.01007 |
| 531 | Ralgps2 | 0.0034637 | -1.6685 | 0.217 | 0.341 | 1 |
| 203 | Tspan32 | 5.60E-07 | -1.66859 | 0.043 | 0.331 | 0.01172 |
| 245 | Dock8 | 2.47E-06 | -1.67005 | 0.087 | 0.349 | 0.05169 |
| 176 | Lef1 | 1.45E-07 | -1.67334 | 0.058 | 0.381 | 0.00303 |
| 356 | H2-Aa | 0.0001001 | -1.67348 | 0.261 | 0.45 | 1 |
| 237 | Etnk1 | 2.26E-06 | -1.67516 | 0.174 | 0.429 | 0.0474 |
| 100 | Btg1 | 9.89E-12 | -1.68587 | 0.377 | 0.725 | 2.07E-07 |
| 117 | Arhgap1! | 3.30E-10 | -1.68744 | 0.087 | 0.49 | 6.90E-06 |
| 106 | Arhgap4! | 5.33E-11 | -1.69363 | 0.217 | 0.606 | 1.12E-06 |
| 70 | H2-K1 | 5.51E-18 | -1.69413 | 0.522 | 0.855 | 1.15E-13 |
| 389 | R3hdm1 | 0.0001977 | -1.69512 | 0.217 | 0.385 | 1 |
| 227 | Jak1 | 1.74E-06 | -1.6977 | 0.246 | 0.466 | 0.03636 |
| 121 | Laptm5 | 6.21E-10 | -1.69811 | 0.174 | 0.557 | 1.30E-05 |
| 441 | Igkc | 0.0006548 | -1.69935 | 0.536 | 0.614 | 1 |
| 385 | Fchsd2 | 0.0001803 | -1.70034 | 0.203 | 0.395 | 1 |
| 134 | Cd52 | 4.97E-09 | -1.70076 | 0.304 | 0.614 | 0.0001 |
| 147 | Klf2 | 1.38E-08 | -1.72911 | 0.13 | 0.482 | 0.00029 |
| 140 | Stk17b | 8.02E-09 | -1.73605 | 0.232 | 0.544 | 0.00017 |
| 261 | 4930523 | 4.62E-06 | -1.73851 | 0.203 | 0.437 | 0.09674 |
| 169 | 9930111 | 9.61E-08 | -1.75541 | 0.145 | 0.448 | 0.00201 |
| 86 | Mbnl1 | 1.46E-14 | -1.75886 | 0.377 | 0.761 | 3.07E-10 |
| 290 | Btla | 1.36E-05 | -1.75965 | 0.029 | 0.265 | 0.28502 |
| 64 | Malat1 | 1.91E-20 | -1.76609 | 0.623 | 0.968 | 3.99E-16 |
| 247 | Samhd1 | 2.57E-06 | -1.77005 | 0.217 | 0.465 | 0.05386 |
| 158 | Gimap1 | 3.25E-08 | -1.78553 | 0.087 | 0.425 | 0.00068 |

| | | | | | | |
|---|---|---|---|---|---|---|
| 249 | Trbc2 | 2.79E-06 | -1.81839 | 0.13 | 0.4 | 0.05839 |
| 287 | Stat1 | 1.27E-05 | -1.82119 | 0.13 | 0.361 | 0.26688 |
| 770 | Bcl11a | 0.0688311 | -1.83087 | 0.203 | 0.26 | 1 |
| 291 | H2-Ab1 | 1.40E-05 | -1.85082 | 0.217 | 0.452 | 0.29306 |
| 248 | H2-Ob | 2.77E-06 | -1.85906 | 0 | 0.251 | 0.05795 |
| 336 | Trbc1 | 5.78E-05 | -1.88203 | 0.058 | 0.271 | 1 |
| 119 | Sp100 | 4.72E-10 | -1.88335 | 0.145 | 0.524 | 9.88E-06 |
| 185 | Stap1 | 3.02E-07 | -1.89839 | 0.058 | 0.367 | 0.00633 |
| 172 | Ankrd44 | 1.12E-07 | -1.93103 | 0.174 | 0.466 | 0.00234 |
| 118 | Foxp1 | 3.65E-10 | -1.94846 | 0.261 | 0.587 | 7.64E-06 |
| 67 | Ptprc | 3.34E-19 | -1.96599 | 0.29 | 0.837 | 7.00E-15 |
| 150 | Dock10 | 1.43E-08 | -1.97001 | 0.116 | 0.456 | 0.0003 |
| 101 | Arhgef1 | 1.03E-11 | -1.97334 | 0.275 | 0.641 | 2.15E-07 |
| 125 | Dock2 | 1.44E-09 | -1.97526 | 0.145 | 0.502 | 3.02E-05 |
| 92 | Rhoh | 6.90E-13 | -1.98258 | 0.087 | 0.571 | 1.44E-08 |
| 246 | Il7r | 2.53E-06 | -1.98514 | 0.058 | 0.322 | 0.05297 |
| 129 | Gimap6 | 2.48E-09 | -1.98792 | 0.072 | 0.439 | 5.19E-05 |
| 123 | Ripor2 | 1.15E-09 | -1.99893 | 0.101 | 0.478 | 2.42E-05 |
| 217 | Pou2f2 | 1.00E-06 | -2.01388 | 0.058 | 0.344 | 0.02097 |
| 142 | Hbb-bs | 9.84E-09 | -2.03463 | 0.014 | 0.362 | 0.00021 |
| 212 | Bank1 | 8.36E-07 | -2.04309 | 0.029 | 0.321 | 0.0175 |
| 139 | Rcsd1 | 7.96E-09 | -2.12155 | 0.072 | 0.414 | 0.00017 |
| 161 | Itga4 | 4.41E-08 | -2.1457 | 0.058 | 0.375 | 0.00092 |
| 97 | Shisa5 | 2.14E-12 | -2.15119 | 0.261 | 0.627 | 4.47E-08 |
| 274 | Ms4a1 | 7.33E-06 | -2.16064 | 0.058 | 0.31 | 0.15347 |
| 99 | Ets1 | 4.79E-12 | -2.16874 | 0.203 | 0.6 | 1.00E-07 |
| 194 | Git2 | 4.56E-07 | -2.17872 | 0.188 | 0.442 | 0.00956 |
| 276 | H2-Eb1 | 7.59E-06 | -2.20069 | 0.188 | 0.424 | 0.15889 |
| 104 | Coro1a | 1.96E-11 | -2.21979 | 0.188 | 0.564 | 4.10E-07 |
| 228 | Cd74 | 1.75E-06 | -2.23094 | 0.406 | 0.605 | 0.03666 |
| 167 | Malt1 | 8.65E-08 | -2.25338 | 0.159 | 0.454 | 0.00181 |
| 116 | Ms4a4b | 3.24E-10 | -2.25888 | 0.072 | 0.483 | 6.79E-06 |
| 294 | Gm3124 | 1.46E-05 | -2.26713 | 0.029 | 0.26 | 0.30636 |
| 229 | Cd79a | 1.81E-06 | -2.27327 | 0.029 | 0.301 | 0.03796 |
| 102 | BE69200 | 1.39E-11 | -2.31723 | 0.058 | 0.503 | 2.92E-07 |
| 231 | Kcnq1ot | 2.00E-06 | -2.33544 | 0.232 | 0.461 | 0.04197 |
| 87 | Ighm | 1.49E-14 | -2.53151 | 0.188 | 0.675 | 3.13E-10 |
| 206 | Mef2c | 6.24E-07 | -2.56265 | 0.101 | 0.385 | 0.01307 |
| 95 | Ifi203 | 1.45E-12 | -2.64831 | 0.087 | 0.524 | 3.05E-08 |
| 157 | Ighd | 3.14E-08 | -2.70098 | 0.029 | 0.357 | 0.00066 |
| 148 | Ebf1 | 1.38E-08 | -2.75377 | 0.072 | 0.422 | 0.00029 |
| 91 | Cd37 | 3.60E-13 | -2.76554 | 0.087 | 0.543 | 7.53E-09 |

| pathway | pval | padj | log2err | ES | NES | size | LeadingEdge |
|---|---|---|---|---|---|---|---|
| ACEVEDO_ | 7.45E-02 | 1.90E-01 | 0.2572065 | 0.3395 | 1.40902 | 38 | Krt8, Mt.... |
| ACEVEDO_ | 6.67E-01 | 8.33E-01 | 0.0541201 | -0.183 | -0.89433 | 97 | Cd74, Et.... |
| ACEVEDO_ | 5.35E-01 | 7.26E-01 | 0.0670513 | -0.281 | -0.93287 | 19 | Dock8, I.... |
| ACEVEDO_ | 7.35E-02 | 1.90E-01 | 0.2616635 | 0.4554 | 1.45907 | 16 | Mt1, Mt2.... |
| ACEVEDO_ | 7.72E-01 | 9.08E-01 | 0.0713053 | 0.1685 | 0.83906 | 84 | Cystm1, .... |
| ACEVEDO_ | 2.90E-01 | 4.88E-01 | 0.0999277 | -0.27 | -1.11429 | 42 | Malt1, R.... |
| ACEVEDO_ | 3.66E-03 | 2.16E-02 | 0.4317077 | 0.4804 | 1.87975 | 31 | Mt1, Cld.... |
| ACEVEDO_ | 8.35E-01 | 9.44E-01 | 0.0461379 | -0.189 | -0.72156 | 30 | Rbm5, My.... |
| AFFAR_YY | 1.76E-01 | 3.56E-01 | 0.133555 | -0.391 | -1.27435 | 18 | Ighm, It.... |
| AFFAR_YY | 2.54E-03 | 1.63E-02 | 0.4317077 | 0.6116 | 1.91829 | 15 | Clu, Krt.... |
| AGUIRRE_ | 9.88E-01 | 1.00E+00 | 0.0555947 | 0.1223 | 0.4567 | 27 | Nfib, Cl.... |
| AGUIRRE_ | 3.88E-01 | 5.98E-01 | 0.1055209 | 0.2688 | 1.06056 | 33 | Nme2, As.... |
| ALCALAY_ | 1.71E-01 | 3.52E-01 | 0.1357409 | -0.396 | -1.29286 | 18 | H2-Eb1, .... |
| ALCALA_AI | 1.31E-01 | 2.86E-01 | 0.1596467 | -0.385 | -1.33934 | 23 | Cd74, Rh.... |
| ALFANO_M | 3.84E-01 | 5.93E-01 | 0.1059203 | 0.2882 | 1.06443 | 26 | Anxa1, S.... |
| ALONSO_M | 8.86E-03 | 4.25E-02 | 0.3807304 | 0.4295 | 1.73663 | 36 | Tceal9, .... |
| AMIT_EGF_ | 4.96E-02 | 1.45E-01 | 0.3217759 | 0.4638 | 1.59511 | 20 | Mt1, Mt2.... |
| AMUNDSOI | 7.48E-01 | 8.91E-01 | 0.0687943 | 0.2173 | 0.80234 | 26 | Mt1, Mt2.... |
| ANDERSEI | 1.10E-01 | 2.51E-01 | 0.2114002 | 0.4333 | 1.38813 | 16 | Epcam, C.... |
| APPIERTO_ | 3.77E-03 | 2.20E-02 | 0.4317077 | 0.5986 | 1.91775 | 16 | Ifitm3, .... |
| APRELIKO | 4.06E-02 | 1.25E-01 | 0.3217759 | 0.4905 | 1.53847 | 15 | Krt8, Tp.... |
| AZARE_NE | 2.75E-03 | 1.74E-02 | 0.4317077 | 0.552 | 1.93239 | 21 | S100a9, .... |
| BAELDE_D | 4.99E-01 | 6.95E-01 | 0.0937465 | 0.2121 | 0.98796 | 63 | Nupr1, D.... |
| BAE_BRCA | 7.22E-03 | 3.71E-02 | 0.4070179 | 0.5536 | 1.79684 | 17 | Krt19, C.... |
| BASAKI_YE | 4.26E-01 | 6.36E-01 | 0.0782955 | -0.239 | -1.01514 | 46 | Btg1, Ra.... |
| BASAKI_YE | 6.99E-02 | 1.83E-01 | 0.2663507 | 0.4211 | 1.44828 | 20 | Mt1, Mt2.... |
| BASSO_B_ | 4.48E-01 | 6.56E-01 | 0.0758887 | -0.257 | -1.01078 | 34 | Ripor2, .... |
| BASSO_CD | 5.22E-03 | 2.94E-02 | 0.4070179 | -0.526 | -1.74374 | 19 | Mef2c, C.... |
| BENPORAT | 1.45E-01 | 3.12E-01 | 0.1464162 | -0.281 | -1.26601 | 61 | Stat1, M.... |
| BENPORAT | 6.89E-01 | 8.54E-01 | 0.0721798 | 0.2143 | 0.83859 | 31 | Mt1, Mt2.... |
| BENPORAT | 2.12E-01 | 4.05E-01 | 0.1473312 | 0.3297 | 1.21513 | 25 | Krt8, Ep.... |
| BENPORAT | 7.43E-02 | 1.90E-01 | 0.2139279 | -0.45 | -1.49159 | 19 | Ebf1, It.... |
| BENPORAT | 9.95E-01 | 1.00E+00 | 0.059476 | 0.114 | 0.5928 | 114 | Mt2, Fab.... |
| BENPORAT | 6.41E-01 | 8.16E-01 | 0.0776799 | 0.2064 | 0.89411 | 45 | Txn1, Ju.... |
| BENPORAT | 6.31E-01 | 8.09E-01 | 0.0799859 | 0.179 | 0.91457 | 92 | Krt18, S.... |
| BENPORAT | 5.15E-01 | 7.12E-01 | 0.0689567 | -0.284 | -0.94331 | 19 | Fus, Tra.... |
| BENPORAT | 2.54E-02 | 8.92E-02 | 0.3524879 | 0.4411 | 1.59488 | 23 | S100a9, .... |

| | | | | | | | |
|---|---|---|---|---|---|---|---|
| BENPORAT | 9.68E-01 | 1.00E+00 | 0.0369161 | -0.136 | -0.6484 | 84 | Foxp1, B.... |
| BENPORAT | 4.00E-01 | 6.09E-01 | 0.1035763 | 0.3014 | 1.06377 | 22 | Mt1, Mt2.... |
| BERENJEN | 3.19E-02 | 1.07E-01 | 0.3217759 | 0.3165 | 1.43947 | 57 | Mt1, Mt2.... |
| BERENJEN | 1.19E-02 | 5.26E-02 | 0.3807304 | 0.346 | 1.60538 | 62 | Spp1, Tx.... |
| BERTUCCI_ | 2.60E-01 | 4.67E-01 | 0.1071402 | -0.352 | -1.16767 | 19 | H2-Ob, S.... |
| BHATI_G2M | 4.58E-01 | 6.64E-01 | 0.095288 | 0.3185 | 1.03384 | 17 | Mt1, Nup.... |
| BHAT_ESR | 3.79E-01 | 5.87E-01 | 0.1088201 | 0.3425 | 1.07426 | 15 | Tff2, Fo.... |
| BIDUS_ME | 2.19E-01 | 4.14E-01 | 0.1167392 | -0.386 | -1.20806 | 15 | Coro1a, .... |
| BIDUS_ME | 9.88E-01 | 1.00E+00 | 0.0387962 | -0.133 | -0.50222 | 29 | Zfp318, .... |
| BILANGES_ | 4.41E-01 | 6.49E-01 | 0.0972151 | 0.3065 | 1.02837 | 19 | Serpinb6.... |
| BILANGES_ | 3.85E-01 | 5.93E-01 | 0.1039585 | 0.299 | 1.08126 | 23 | Ptms, Ee.... |
| BILBAN_B_ | 5.81E-01 | 7.66E-01 | 0.0619763 | -0.292 | -0.9123 | 15 | Mndal, S.... |
| BILD_HRAS | 3.64E-01 | 5.70E-01 | 0.1083943 | 0.2985 | 1.08721 | 24 | S100a6, .... |
| BLALOCK_ | 3.61E-01 | 5.70E-01 | 0.1167392 | 0.1848 | 1.05237 | 181 | Epcam, T.... |
| BLALOCK_ | 9.01E-01 | 9.71E-01 | 0.0426734 | -0.176 | -0.67 | 30 | Ccnl1, H.... |
| BLALOCK_ | 4.15E-01 | 6.23E-01 | 0.1017139 | 0.2489 | 1.03428 | 39 | Clu, Mt1.... |
| BLALOCK_ | 4.80E-01 | 6.80E-01 | 0.09855 | 0.1769 | 0.98341 | 158 | Reg3b, C.... |
| BLANCO_M | 7.79E-04 | 6.28E-03 | 0.4772708 | -0.579 | -2.00476 | 22 | Malt1, E.... |
| BLANCO_M | 4.93E-03 | 2.81E-02 | 0.4070179 | -0.419 | -1.79474 | 48 | Mef2c, S.... |
| BLANCO_M | 3.41E-03 | 2.07E-02 | 0.4317077 | 0.6012 | 1.92611 | 16 | Spp1, Ct.... |
| BLANCO_M | 6.13E-02 | 1.72E-01 | 0.2377938 | -0.384 | -1.46895 | 31 | Pou2f2, .... |
| BLANCO_M | 3.00E-02 | 1.02E-01 | 0.3524879 | -0.396 | -1.59846 | 37 | Pou2f2, .... |
| BLANCO_M | 3.33E-01 | 5.38E-01 | 0.0919686 | -0.332 | -1.08396 | 18 | Sp100, S.... |
| BLANCO_M | 7.73E-02 | 1.95E-01 | 0.208955 | -0.447 | -1.45877 | 18 | Cd37, Ri.... |
| BLANCO_M | 3.51E-05 | 5.60E-04 | 0.5573322 | -0.677 | -2.16419 | 17 | Ets1, Po.... |
| BLUM_RES | 8.40E-01 | 9.47E-01 | 0.0636424 | 0.1846 | 0.75208 | 37 | Sox4, So.... |
| BLUM_RES | 1.71E-02 | 6.86E-02 | 0.3524879 | 0.4001 | 1.66054 | 38 | Nupr1, C.... |
| BOCHKIS_ | 3.73E-02 | 1.20E-01 | 0.3217759 | 0.4043 | 1.58208 | 31 | Mt1, Mt2.... |
| BONOME_ | 6.17E-01 | 8.00E-01 | 0.0603786 | -0.259 | -0.88675 | 21 | Sp100, C.... |
| BONOME_ | 2.71E-02 | 9.40E-02 | 0.3524879 | -0.43 | -1.59671 | 27 | Mef2c, A.... |
| BOQUEST_ | 6.55E-02 | 1.77E-01 | 0.2765006 | 0.328 | 1.44815 | 47 | Mt1, Krt.... |
| BOQUEST_ | 1.55E-03 | 1.05E-02 | 0.4550599 | -0.562 | -1.9257 | 21 | Cd74, H2.... |
| BORCZUK_ | 1.24E-04 | 1.40E-03 | 0.5188481 | -0.673 | -2.10377 | 15 | Ighm, Cd.... |
| BORCZUK_ | 8.13E-01 | 9.33E-01 | 0.0645031 | 0.1703 | 0.76684 | 50 | S100a11,.... |
| BOSCO_AL | 5.93E-02 | 1.68E-01 | 0.24134 | -0.442 | -1.51553 | 21 | Mef2c, S.... |
| BOUDOUKI | 1.74E-01 | 3.55E-01 | 0.1656567 | 0.3998 | 1.28092 | 16 | S100a6, .... |
| BOYAULT_ | 4.03E-02 | 1.24E-01 | 0.3217759 | 0.4605 | 1.50884 | 18 | Sox9, Cl.... |
| BOYAULT_ | 2.72E-02 | 9.40E-02 | 0.3524879 | 0.4531 | 1.66982 | 25 | Txn1, Ma.... |

| | | | | | | | |
|---|---|---|---|---|---|---|---|
| BOYLAN_M | 3.92E-02 | 1.23E-01 | 0.3217759 | 0.3939 | 1.55212 | 32 | S100a9, .... |
| BOYLAN_M | 1.17E-01 | 2.65E-01 | 0.1656567 | -0.447 | -1.39662 | 15 | Ebf1, Rh.... |
| BOYLAN_M | 1.15E-01 | 2.61E-01 | 0.2065879 | 0.4203 | 1.36436 | 17 | S100a9, .... |
| BROCKE_A | 8.35E-02 | 2.03E-01 | 0.2450418 | 0.4284 | 1.40358 | 18 | Spp1, Mt.... |
| BROWNE_I | 6.76E-01 | 8.41E-01 | 0.0558265 | -0.236 | -0.85696 | 26 | Ripor2, .... |
| BROWNE_I | 7.23E-01 | 8.79E-01 | 0.0525899 | -0.249 | -0.81109 | 18 | Lef1, Mn.... |
| BROWNE_I | 5.51E-01 | 7.40E-01 | 0.0847985 | 0.2926 | 0.93754 | 16 | Anxa2, C.... |
| BROWNE_I | 4.15E-01 | 6.23E-01 | 0.1009906 | 0.2481 | 1.04079 | 40 | Spp1, An.... |
| BROWN_M | 5.61E-01 | 7.49E-01 | 0.0838361 | 0.2842 | 0.92266 | 17 | Sox4, If.... |
| BROWN_M | 1.93E-02 | 7.51E-02 | 0.3524879 | 0.4769 | 1.68323 | 22 | S100a6, .... |
| BRUECKNE | 2.25E-01 | 4.23E-01 | 0.1464162 | 0.3885 | 1.21854 | 15 | Reg3g, C.... |
| BRUINS_U | 2.05E-01 | 3.97E-01 | 0.1238422 | -0.345 | -1.23231 | 25 | Pou2f2, .... |
| BRUINS_U | 7.99E-01 | 9.27E-01 | 0.0687943 | 0.1678 | 0.82185 | 79 | Ctsl, An.... |
| BRUINS_U | 3.92E-01 | 6.01E-01 | 0.1059203 | 0.24 | 1.03951 | 45 | Clps, Cd.... |
| BRUINS_U | 1.24E-01 | 2.78E-01 | 0.197822 | 0.3227 | 1.30774 | 35 | S100a6, .... |
| BRUINS_U | 1.33E-01 | 2.90E-01 | 0.1563124 | -0.404 | -1.34166 | 19 | Arhgap45.... |
| BURTON_A | 9.09E-02 | 2.15E-01 | 0.2377938 | 0.4569 | 1.43294 | 15 | Fabp5, U.... |
| BURTON_A | 1.87E-02 | 7.31E-02 | 0.3524879 | 0.452 | 1.63418 | 23 | S100a8, .... |
| BURTON_A | 2.90E-01 | 4.88E-01 | 0.1250334 | 0.3481 | 1.14032 | 18 | Dstn, Tc.... |
| BUYTAERT | 1.84E-01 | 3.66E-01 | 0.1314576 | -0.307 | -1.24219 | 38 | Coro1a, .... |
| BUYTAERT | 2.64E-01 | 4.70E-01 | 0.104344 | -0.244 | -1.1389 | 77 | Stk17b, .... |
| BYSTRYKH | 9.50E-01 | 9.98E-01 | 0.0559429 | 0.1638 | 0.59228 | 23 | Fabp5, N.... |
| BYSTRYKH | 6.23E-01 | 8.04E-01 | 0.0583453 | -0.197 | -0.90127 | 66 | Cd79a, C.... |
| CAIRO_HE | 7.42E-01 | 8.86E-01 | 0.0517406 | -0.233 | -0.79658 | 21 | 4930523C.... |
| CAIRO_HE | 2.69E-02 | 9.40E-02 | 0.3524879 | 0.2903 | 1.45146 | 87 | Epcam, K.... |
| CAIRO_HE | 8.09E-02 | 1.98E-01 | 0.2489111 | 0.3984 | 1.40609 | 22 | Epcam, S.... |
| CAIRO_LIV | 8.82E-02 | 2.10E-01 | 0.2377938 | 0.443 | 1.41916 | 16 | S100a9, .... |
| CASORELL | 8.25E-01 | 9.39E-01 | 0.0673624 | 0.1622 | 0.80847 | 85 | Ctsl, Cd.... |
| CASORELL | 2.31E-01 | 4.31E-01 | 0.1404062 | 0.3279 | 1.19415 | 24 | Mt1, Krt.... |
| CAVARD_L | 3.98E-04 | 3.71E-03 | 0.4984931 | 0.6664 | 2.08992 | 15 | Reg3b, R.... |
| CHANDRAN | 2.72E-01 | 4.81E-01 | 0.1294429 | 0.2853 | 1.14091 | 34 | Spp1, Nu.... |
| CHARAFE_ | 8.56E-03 | 4.14E-02 | 0.3807304 | 0.5686 | 1.78346 | 15 | S100a8, .... |
| CHARAFE_ | 1.82E-01 | 3.64E-01 | 0.1608014 | 0.2719 | 1.22417 | 50 | Mt1, Mt2.... |
| CHARAFE_ | 6.59E-01 | 8.30E-01 | 0.0753094 | 0.2368 | 0.88462 | 27 | Krt19, P.... |
| CHARAFE_ | 8.20E-01 | 9.38E-01 | 0.046882 | -0.188 | -0.77241 | 41 | Malt1, E.... |
| CHARAFE_ | 6.79E-05 | 8.65E-04 | 0.5384341 | 0.5656 | 2.22878 | 32 | S100a8, .... |
| CHAUHAN_ | 1.26E-01 | 2.80E-01 | 0.1596467 | -0.309 | -1.31012 | 45 | Cd74, Tr.... |
| CHEMNITZ | 3.66E-02 | 1.18E-01 | 0.3217759 | 0.4667 | 1.52898 | 18 | Krt7, Cl.... |

| | | | | | | | |
|---|---|---|---|---|---|---|---|
| CHEN_HO) | 4.63E-01 | 6.67E-01 | 0.0745501 | -0.265 | -0.9902 | 28 | Rbm5, Sm.... |
| CHEN_MET | 2.03E-01 | 3.94E-01 | 0.1193484 | -0.232 | -1.16109 | 113 | Cd37, Cd.... |
| CHIANG_LI | 1.72E-01 | 3.52E-01 | 0.1695706 | 0.4134 | 1.29652 | 15 | Sox4, So.... |
| CHIANG_LI | 1.59E-01 | 3.37E-01 | 0.1737478 | 0.3394 | 1.26785 | 27 | Ybx1, Ps.... |
| CHIARADO | 1.64E-02 | 6.75E-02 | 0.3524879 | 0.4888 | 1.71111 | 21 | Nupr1, D.... |
| CHIARADO | 4.22E-01 | 6.33E-01 | 0.0791317 | -0.307 | -1.0175 | 19 | Mndal, S.... |
| CHIARADO | 4.61E-02 | 1.38E-01 | 0.3217759 | 0.4637 | 1.50501 | 17 | Nupr1, C.... |
| CHIARADO | 5.05E-01 | 7.01E-01 | 0.0897105 | 0.3026 | 0.98211 | 17 | Spp1, Ct.... |
| CHICAS_RI | 4.92E-04 | 4.31E-03 | 0.4772708 | 0.4249 | 1.95978 | 58 | Clu, S10.... |
| CHICAS_RI | 4.16E-02 | 1.27E-01 | 0.3217759 | 0.4671 | 1.51614 | 17 | Clu, Nup.... |
| CHICAS_RI | 3.48E-03 | 2.10E-02 | 0.4317077 | 0.5207 | 1.88264 | 23 | Mt1, Sox.... |
| CHICAS_RI | 7.87E-02 | 1.96E-01 | 0.2065879 | -0.332 | -1.39667 | 44 | Pou2f2, .... |
| CHNG_MUI | 5.95E-01 | 7.81E-01 | 0.0808589 | 0.2768 | 0.90679 | 18 | Klk1, Ee.... |
| CHUNG_BL | 4.91E-02 | 1.45E-01 | 0.3217759 | 0.4522 | 1.48146 | 18 | Mt1, Cts.... |
| CHYLA_CB | 1.08E-01 | 2.47E-01 | 0.2139279 | 0.3606 | 1.34698 | 27 | Krt18, K.... |
| COLINA_TA | 8.74E-02 | 2.08E-01 | 0.195789 | -0.361 | -1.41079 | 33 | Shisa5, .... |
| CONCANN( | 2.11E-01 | 4.04E-01 | 0.1492075 | 0.389 | 1.24617 | 16 | Cd24a, F.... |
| CONCANN( | 1.28E-03 | 9.34E-03 | 0.4550599 | 0.5772 | 2.02058 | 21 | Clu, Spp.... |
| CREIGHTO | 6.48E-01 | 8.21E-01 | 0.0574877 | -0.21 | -0.88597 | 44 | Jak1, Do.... |
| CREIGHTO | 8.50E-02 | 2.05E-01 | 0.24134 | 0.3144 | 1.38828 | 47 | S100a6, .... |
| CREIGHTO | 8.89E-01 | 9.67E-01 | 0.0597318 | 0.1936 | 0.66594 | 20 | Mt2, Gst.... |
| CREIGHTO | 8.79E-01 | 9.63E-01 | 0.0430173 | -0.173 | -0.69733 | 36 | Malt1, F.... |
| CUI_TCF21 | 1.99E-06 | 6.90E-05 | 0.6272567 | -0.45 | -2.14446 | 86 | Ebf1, Me.... |
| CUI_TCF21 | 3.91E-03 | 2.26E-02 | 0.4070179 | 0.47 | 1.87974 | 34 | Clu, Krt.... |
| DACOSTA_ | 3.39E-04 | 3.29E-03 | 0.4984931 | -0.427 | -1.92587 | 61 | Il7r, Mb.... |
| DACOSTA_ | 3.18E-05 | 5.25E-04 | 0.5573322 | -0.395 | -1.95125 | 103 | Malt1, I.... |
| DACOSTA_ | 9.14E-01 | 9.77E-01 | 0.0596037 | 0.159 | 0.66059 | 39 | Ctsl, Rh.... |
| DAIRKEE_( | 6.40E-01 | 8.15E-01 | 0.0768737 | 0.2417 | 0.90278 | 27 | Txn1, Tp.... |
| DANG_BOU | 6.80E-01 | 8.45E-01 | 0.078921 | 0.1634 | 0.90805 | 158 | Mt2, Txn.... |
| DANG_MY( | 8.24E-01 | 9.39E-01 | 0.0642141 | 0.1879 | 0.75172 | 34 | Txn1, Nm.... |
| DANG_RE( | 4.79E-02 | 1.42E-01 | 0.3217759 | 0.36 | 1.54475 | 44 | Mt1, Col.... |
| DARWICHE | 6.27E-02 | 1.73E-01 | 0.2820134 | 0.4289 | 1.4751 | 20 | Spp1, Ct.... |
| DARWICHE | 2.82E-03 | 1.77E-02 | 0.4317077 | 0.5173 | 1.91033 | 26 | S100a9, .... |
| DARWICHE | 8.54E-02 | 2.05E-01 | 0.24134 | 0.4233 | 1.42047 | 19 | Spp1, Mt.... |
| DARWICHE | 1.08E-03 | 8.23E-03 | 0.4550599 | 0.5236 | 1.95586 | 27 | S100a9, .... |
| DARWICHE | 6.18E-02 | 1.72E-01 | 0.3217759 | 0.4103 | 1.5123 | 25 | Mt1, S10.... |
| DARWICHE | 3.24E-03 | 1.99E-02 | 0.4317077 | 0.5448 | 1.90716 | 21 | S100a9, .... |
| DARWICHE | 4.95E-01 | 6.91E-01 | 0.0908241 | 0.3059 | 0.98003 | 16 | App, Asn.... |

| | | | | | | | |
|---|---|---|---|---|---|---|---|
| DARWICHE | 9.00E-05 | 1.10E-03 | 0.5384341 | 0.5638 | 2.17278 | 30 | S100a9, .... |
| DAVICIONI_ | 9.62E-01 | 1.00E+00 | 0.0405756 | -0.157 | -0.54613 | 23 | Ripor2, .... |
| DAVICIONI_ | 7.73E-02 | 1.95E-01 | 0.2529611 | 0.3674 | 1.40729 | 29 | Krt18, C.... |
| DAZARD_R | 1.12E-03 | 8.45E-03 | 0.4550599 | -0.534 | -1.94059 | 26 | Zfp318, .... |
| DAZARD_R | 9.56E-03 | 4.50E-02 | 0.3807304 | 0.3899 | 1.68915 | 45 | S100a9, .... |
| DAZARD_R | 6.73E-01 | 8.39E-01 | 0.0739901 | 0.2541 | 0.85261 | 19 | Anxa2, F.... |
| DAZARD_U | 2.45E-01 | 4.52E-01 | 0.1364904 | 0.3351 | 1.17317 | 21 | Anxa2, T.... |
| DEBIASI_A | 4.58E-01 | 6.64E-01 | 0.0753094 | -0.295 | -0.99504 | 20 | Bcl11a, .... |
| DEBIASI_A | 4.94E-01 | 6.91E-01 | 0.070961 | -0.269 | -0.97771 | 26 | Sp100, S.... |
| DELACROI | 9.98E-03 | 4.67E-02 | 0.3807304 | 0.3905 | 1.67678 | 43 | Nupr1, S.... |
| DELACROI | 5.21E-01 | 7.17E-01 | 0.0891647 | 0.2359 | 0.95388 | 36 | Spp1, Cd.... |
| DELYS_TH | 2.36E-02 | 8.65E-02 | 0.3524879 | 0.5245 | 1.64503 | 15 | Mt1, Mt2.... |
| DELYS_TH | 1.73E-05 | 3.18E-04 | 0.5756103 | 0.5224 | 2.26284 | 45 | Spp1, Tr.... |
| DER_IFN_A | 2.20E-02 | 8.31E-02 | 0.3524879 | -0.494 | -1.58053 | 17 | Stat1, B.... |
| DER_IFN_B | 1.54E-02 | 6.51E-02 | 0.3807304 | -0.477 | -1.6791 | 24 | Stat1, B.... |
| DER_IFN_G | 2.38E-02 | 8.68E-02 | 0.3524879 | -0.521 | -1.62938 | 16 | Stat1, B.... |
| DESERT_S | 2.12E-05 | 3.62E-04 | 0.5756103 | 0.5507 | 2.31042 | 40 | S100a6, .... |
| DEURIG_T_ | 3.78E-04 | 3.59E-03 | 0.4984931 | -0.465 | -1.96041 | 44 | Malt1, G.... |
| DEURIG_T_ | 1.90E-03 | 1.27E-02 | 0.4550599 | 0.5633 | 1.98816 | 22 | Krt18, C.... |
| DE_YY1_TA | 7.37E-01 | 8.86E-01 | 0.052057 | -0.233 | -0.79963 | 21 | Itga4, C.... |

| pathway | pval | padj | log2err | ES | NES | size | LeadingEdge |
|---|---|---|---|---|---|---|---|
| AAACCAC_MIF | 0.6096 | 0.871 | 0.061436 | -0.287 | -0.8922 | 15 | Bcl11a, .... |
| AAAGACA_MIF | 0.5947 | 0.859 | 0.064214 | -0.26 | -0.9079 | 23 | Bcl11a, .... |
| AAAGGAT_MIF | 0.7907 | 0.976 | 0.063642 | 0.227 | 0.75553 | 17 | Sox4, Er.... |
| AAAGGGA_MIF | 0.4031 | 0.696 | 0.082434 | -0.276 | -1.0315 | 28 | Mbnl1, T.... |
| AAANWWTGC_ | 0.1311 | 0.373 | 0.160801 | -0.418 | -1.3391 | 17 | Ebf1, Me.... |
| AAAYRNCTG_U | 0.7624 | 0.961 | 0.06407 | 0.215 | 0.78373 | 24 | Col1a2, .... |
| AAAYWAACM_ | 0.0938 | 0.306 | 0.193813 | -0.4 | -1.3998 | 23 | Mef2c, S.... |
| AACATTC_MIR | 0.9037 | 1 | 0.056648 | 0.178 | 0.65182 | 25 | Anxa2, C.... |
| AACTGGA_MIF | 0.6775 | 0.905 | 0.05653 | -0.228 | -0.8697 | 30 | Ssh2, Ce.... |
| AACTTT_UNKN | 0.6372 | 0.885 | 0.055479 | -0.178 | -0.9219 | 136 | Ebf1, Me.... |
| AAGCAAT_MIR | 0.2191 | 0.496 | 0.120433 | -0.331 | -1.2071 | 26 | Satb1, A.... |
| AAGCACA_MIF | 0.4394 | 0.733 | 0.078296 | -0.259 | -1.0097 | 33 | Mbnl1, S.... |
| AAGCACT_MIF | 0.1664 | 0.421 | 0.1396 | -0.335 | -1.2758 | 30 | Mef2c, R.... |
| AAGCCAT_MIF | 0.0062 | 0.08 | 0.407018 | -0.457 | -1.7835 | 33 | Mef2c, B.... |
| AAGWWRNYG | 0.8554 | 1 | 0.04707 | -0.223 | -0.7023 | 16 | Ube2d3, .... |
| AATGTGA_MIR | 0.0023 | 0.045 | 0.431708 | -0.419 | -1.7868 | 49 | Mef2c, B.... |
| ACACTAC_MIR | 0.4486 | 0.745 | 0.076675 | -0.324 | -1.0081 | 15 | Smg1, Bc.... |
| ACACTGG_MIF | 0.075 | 0.265 | 0.216543 | -0.417 | -1.4157 | 21 | Ets1, Ce.... |
| ACATTCC_MIR | 0.6378 | 0.885 | 0.05986 | -0.216 | -0.8863 | 42 | Ets1, Fo.... |
| ACAWNRNSRC | 0.5342 | 0.812 | 0.067834 | -0.304 | -0.9469 | 15 | Ddx5, Pr.... |
| ACCAAAG_MIF | 0.019 | 0.134 | 0.352488 | -0.394 | -1.6017 | 41 | Foxp1, B.... |
| ACCATTT_MIR | 0.2751 | 0.567 | 0.105125 | -0.294 | -1.1447 | 32 | Foxp1, M.... |
| ACTAYRNNNC | 0.9603 | 1 | 0.040823 | -0.153 | -0.5875 | 31 | R3hdm1, .... |
| ACTGAAA_MIR | 0.1265 | 0.364 | 0.16318 | -0.36 | -1.3269 | 27 | Mef2c, F.... |
| ACTGCCT_MIF | 0.1149 | 0.345 | 0.170932 | -0.365 | -1.3621 | 28 | Ets1, Fo.... |
| ACTGTAG_MIF | 0.8897 | 1 | 0.044954 | -0.199 | -0.6664 | 20 | Ets1, Foxp1 |
| ACTGTGA_MIF | 0.8652 | 1 | 0.060119 | 0.189 | 0.73485 | 31 | Nedd4, M.... |
| ACTTTAT_MIR | 0.4706 | 0.763 | 0.074739 | -0.254 | -0.9855 | 32 | Btg1, Ss.... |
| ADA2_TARGET | 0.9929 | 1 | 0.053898 | 0.116 | 0.53 | 54 | Anxa2, S.... |
| AEBP2_TARGE | 1 | 1 | 0.057488 | 0.096 | 0.46938 | 80 | Cldn7, P.... |
| AFP1_Q6 | 0.2084 | 0.483 | 0.124434 | -0.366 | -1.2259 | 20 | Foxp1, A.... |
| AGCACTT_MIR | 0.0666 | 0.248 | 0.227987 | -0.357 | -1.4385 | 37 | Mef2c, B.... |
| AGCATTA_MIR | 0.1501 | 0.401 | 0.149208 | -0.384 | -1.3059 | 21 | Ets1, Ss.... |
| AHRR_TARGE | 0.9728 | 1 | 0.036685 | -0.134 | -0.6784 | 109 | Ripor2, .... |
| AHR_Q5 | 0.9243 | 1 | 0.055595 | 0.172 | 0.63052 | 25 | Dstn, Sp.... |
| ALKBH3_TARG | 0.5815 | 0.849 | 0.063785 | -0.239 | -0.9108 | 30 | Jak1, Rb.... |
| ALPHACP1_01 | 0.9026 | 1 | 0.057246 | 0.183 | 0.65381 | 22 | Cdh1, Co.... |
| AML1_01 | 0.2042 | 0.476 | 0.12564 | -0.366 | -1.2439 | 21 | Tspan32,.... |
| AML1_Q6 | 0.2042 | 0.476 | 0.12564 | -0.366 | -1.2439 | 21 | Tspan32,.... |
| AML_Q6 | 0.3053 | 0.605 | 0.098212 | -0.296 | -1.1272 | 30 | Il7r, Ar.... |
| AP1FJ_Q2 | 0.3388 | 0.649 | 0.111463 | 0.295 | 1.10407 | 27 | Dstn, Tc.... |
| AP1_01 | 0.8729 | 1 | 0.059476 | 0.183 | 0.68599 | 27 | Krt8, Ta.... |

| | | | | | | | |
|---|---|---|---|---|---|---|---|
| AP1_C | 0.8857 | 1 | 0.043976 | -0.205 | -0.669 | 18 | Laptm5, Ssh2 |
| AP1_Q2 | 0.4518 | 0.749 | 0.09226 | 0.276 | 1.01205 | 25 | Dstn, Tx.... |
| AP1_Q2_01 | 0.4005 | 0.695 | 0.099233 | 0.286 | 1.03468 | 23 | Krt8, Co.... |
| AP1_Q4 | 0.3561 | 0.662 | 0.108394 | 0.274 | 1.07456 | 32 | Krt19, D.... |
| AP1_Q4_01 | 0.8028 | 0.976 | 0.0628 | 0.211 | 0.73617 | 20 | Krt8, Ga.... |
| AP1_Q6 | 0.2347 | 0.508 | 0.140406 | 0.31 | 1.1702 | 29 | Dstn, Ta.... |
| AP1_Q6_01 | 0.9361 | 1 | 0.041993 | -0.162 | -0.6248 | 31 | Git2, La.... |
| AP2ALPHA_01 | 0.7007 | 0.922 | 0.069448 | 0.231 | 0.82337 | 22 | Sox9, Cl.... |
| AP2GAMMA_01 | 0.8121 | 0.979 | 0.062249 | 0.211 | 0.75369 | 22 | Sox9, Cl.... |
| AP2REP_01 | 0.7483 | 0.952 | 0.051846 | -0.251 | -0.7807 | 15 | Cd3d, Cd.... |
| AP2_Q3 | 0.8642 | 1 | 0.046602 | -0.222 | -0.6978 | 16 | Smg1, Bc.... |
| AP2_Q6 | 0.1913 | 0.461 | 0.130106 | -0.339 | -1.2343 | 26 | Wnk1, Ra.... |
| AP2_Q6_01 | 0.2643 | 0.552 | 0.107972 | -0.319 | -1.1609 | 26 | Wnk1, Ra.... |
| AP3_Q6 | 0.5381 | 0.815 | 0.06912 | -0.271 | -0.9481 | 23 | Mbnl1, T.... |
| AP4_01 | 0.6776 | 0.905 | 0.057006 | -0.258 | -0.8538 | 19 | Foxp1, B.... |
| AP4_Q5 | 0.056 | 0.237 | 0.252961 | -0.45 | -1.5069 | 20 | Arhgap45.... |
| AP4_Q6 | 0.0226 | 0.142 | 0.352488 | -0.486 | -1.5857 | 18 | Lck, Bcl.... |
| AREB6_01 | 0.0473 | 0.219 | 0.321776 | 0.482 | 1.61584 | 18 | S100a9, .... |
| AREB6_02 | 0.2588 | 0.543 | 0.130106 | 0.348 | 1.19674 | 19 | Sox4, Cy.... |
| AREB6_03 | 0.16 | 0.417 | 0.169571 | 0.377 | 1.29796 | 19 | Krt8, Ep.... |
| AREB6_04 | 0.1604 | 0.417 | 0.142057 | -0.393 | -1.2833 | 18 | Il7r, Bc.... |
| ARHGAP35_TA | 0.7619 | 0.961 | 0.052376 | -0.247 | -0.7756 | 16 | Rbm39, L.... |
| ARID5B_TARG | 0.0404 | 0.204 | 0.287857 | -0.296 | -1.3964 | 77 | Itga4, R.... |
| ARNT2_TARGE | 0.2922 | 0.59 | 0.122108 | 0.244 | 1.10977 | 54 | S100a6, .... |
| ARNT_01 | 0.4683 | 0.762 | 0.07455 | -0.256 | -0.9766 | 30 | Mbnl1, L.... |
| ARNT_02 | 0.9189 | 1 | 0.057731 | 0.168 | 0.63098 | 28 | Gapdh, A.... |
| AR_Q6 | 0.0901 | 0.3 | 0.197822 | -0.395 | -1.4288 | 25 | Mef2c, M.... |
| ASH1L_TARGE | 1 | 1 | 0.057006 | 0.088 | 0.44591 | 102 | Atpif1, .... |
| ATAAGCT_MIR | 0.0595 | 0.241 | 0.245042 | -0.446 | -1.4935 | 20 | Mbnl1, F.... |
| ATACCTC_MIR | 0.116 | 0.345 | 0.169571 | -0.419 | -1.3683 | 18 | 4930523C.... |
| ATACTGT_MIR | 0.8619 | 1 | 0.046882 | -0.201 | -0.7038 | 23 | Ets1, Ce.... |
| ATAGGAA_MIR | 0.116 | 0.345 | 0.169571 | -0.419 | -1.3678 | 18 | Bcl11a, .... |
| ATATGCA_MIR | 0.0002 | 0.021 | 0.518848 | -0.633 | -2.0937 | 19 | Mef2c, B.... |
| ATCATGA_MIR | 0.0619 | 0.243 | 0.24134 | -0.417 | -1.4584 | 23 | Tnrc6b, .... |
| ATF1_Q6 | 0.6353 | 0.885 | 0.075118 | 0.246 | 0.84712 | 19 | Col1a1, .... |
| ATF3_Q6 | 0.9506 | 1 | 0.055479 | 0.163 | 0.6113 | 27 | Cystm1, .... |
| ATF4_Q2 | 0.9792 | 1 | 0.040003 | -0.134 | -0.5215 | 32 | Slc38a2,.... |
| ATF5_TARGET | 0.1831 | 0.448 | 0.131458 | -0.288 | -1.2254 | 48 | Mef2c, S.... |
| ATF6_TARGET | 0.6455 | 0.89 | 0.08042 | 0.184 | 0.9111 | 91 | Mt1, Krt.... |
| ATF_01 | 0.1486 | 0.399 | 0.176694 | 0.322 | 1.278 | 33 | Cystm1, .... |
| ATF_B | 0.7981 | 0.976 | 0.064794 | 0.227 | 0.7616 | 18 | Cystm1, .... |
| ATGAAGG_MIF | 0.9459 | 1 | 0.05571 | 0.17 | 0.58393 | 19 | Marcks, .... |
| ATGCAGT_MIF | 0.0788 | 0.271 | 0.2114 | -0.426 | -1.4282 | 20 | Bcl11a, .... |

| | | | | | | | |
|---|---|---|---|---|---|---|---|
| ATGCTGC_MIF | 0.2346 | 0.508 | 0.116739 | -0.385 | -1.2111 | 16 | Bcl11a, .... |
| ATGCTGG_MIF | 0.0873 | 0.293 | 0.197822 | -0.462 | -1.437 | 15 | Ets1, My.... |
| ATGGYGGA_U | 0.6007 | 0.866 | 0.061976 | -0.276 | -0.8996 | 18 | Cd3e, Bc.... |
| ATGTACA_MIR | 0.0177 | 0.13 | 0.352488 | -0.397 | -1.6144 | 41 | Mef2c, B.... |
| ATGTAGC_MIF | 0.6724 | 0.903 | 0.056648 | -0.258 | -0.8433 | 18 | Ets1, Tc.... |
| ATGTTAA_MIR | 0.0714 | 0.26 | 0.21925 | -0.349 | -1.4075 | 38 | Foxp1, B.... |
| ATGTTTC_MIR | 0.1527 | 0.403 | 0.146416 | -0.34 | -1.2699 | 28 | Celf2, R.... |
| ATTACAT_MIR | 0.0529 | 0.231 | 0.261664 | -0.486 | -1.5265 | 16 | Mef2c, M.... |
| ATTCTTT_MIR | 0.1119 | 0.339 | 0.173748 | -0.325 | -1.3197 | 40 | Mbnl1, T.... |
| ATXN7L3_TAR | 0.6776 | 0.905 | 0.071828 | 0.204 | 0.86574 | 42 | S100a6, .... |
| AUTS2_TARGE | 0.9108 | 1 | 0.042931 | -0.17 | -0.6825 | 36 | Smchd1, .... |
| BACH1_01 | 0.089 | 0.298 | 0.231127 | 0.376 | 1.39736 | 26 | Krt8, Kr.... |
| BACH2_01 | 0.0624 | 0.243 | 0.321776 | 0.465 | 1.62597 | 20 | Tagln2, .... |
| BANP_TARGET | 0.4894 | 0.774 | 0.069613 | -0.206 | -0.9836 | 80 | R3hdm1, .... |
| BARHL1_TARG | 0.8601 | 1 | 0.064794 | 0.156 | 0.78239 | 96 | Dstn, An.... |
| BARX1_TARGE | 1 | 1 | 0.059476 | 0.088 | 0.45684 | 114 | Krt8, S1.... |
| BARX2_TARGE | 1 | 1 | 0.03479 | -0.085 | -0.4309 | 112 | Mef2c, R.... |
| BDP1_TARGET | 0.9737 | 1 | 0.055135 | 0.127 | 0.58241 | 57 | Krt19, K.... |
| BPTF_TARGET | 0.9445 | 1 | 0.04174 | -0.151 | -0.6521 | 53 | Mbnl1, P.... |
| BRCA2_TARGE | 0.981 | 1 | 0.035693 | -0.127 | -0.6442 | 112 | Mbnl1, F.... |
| BRN2_01 | 0.6469 | 0.891 | 0.059604 | -0.271 | -0.8694 | 17 | Bcl11a, .... |
| CACBINDINGP | 0.9859 | 1 | 0.053567 | 0.129 | 0.47783 | 26 | Pbx1, Pt.... |
| CACCAGC_MIF | 0.2769 | 0.57 | 0.10592 | -0.366 | -1.1503 | 16 | Bcl11a, .... |
| CACCCBINDIN | 0.648 | 0.891 | 0.059604 | -0.259 | -0.8686 | 20 | Bcl11a, .... |
| CACGTG_MYC | 0.6535 | 0.891 | 0.054232 | -0.178 | -0.9015 | 110 | Mbnl1, F.... |
| CACTGCC_MIF | 0.075 | 0.265 | 0.216543 | -0.417 | -1.4161 | 21 | Foxp1, L.... |
| CACTGTG_MIF | 0.7856 | 0.974 | 0.04969 | -0.205 | -0.7805 | 30 | Bcl11a, .... |
| CACTTTG_MIR | 0.2815 | 0.577 | 0.104344 | -0.358 | -1.1474 | 17 | Ets1, Ma.... |
| CAGCACT_MIF | 0.2329 | 0.508 | 0.11524 | -0.394 | -1.2241 | 15 | Mbnl1, L.... |
| CAGCAGG_MII | 0.9266 | 1 | 0.041908 | -0.189 | -0.6156 | 18 | Tap2, Sr.... |
| CAGCTG_AP4_ | 0.5244 | 0.806 | 0.066133 | -0.194 | -0.9759 | 106 | Mef2c, E.... |
| CAGCTTT_MIR | 0.1701 | 0.425 | 0.137251 | -0.313 | -1.2629 | 38 | Tnrc6b, .... |
| CAGGTA_AREI | 0.0722 | 0.26 | 0.21925 | -0.33 | -1.3787 | 45 | Cd37, Cd.... |
| CAGTATT_MIR | 0.2218 | 0.498 | 0.118288 | -0.297 | -1.1946 | 37 | Git2, Et.... |
| CAGTGTT_MIR | 0.1514 | 0.401 | 0.146416 | -0.32 | -1.2881 | 38 | Foxp1, M.... |
| CART1_01 | 0.5556 | 0.83 | 0.067362 | -0.296 | -0.931 | 16 | Mef2c, P.... |
| CASP8AP2_TA | 0.9823 | 1 | 0.040823 | -0.149 | -0.5376 | 25 | Stat1, C.... |
| CATGTAA_MIR | 0.3224 | 0.628 | 0.095603 | -0.335 | -1.11 | 19 | Mef2c, T.... |
| CATTGTYY_SC | 0.8451 | 0.994 | 0.046508 | -0.185 | -0.7522 | 40 | Coro1a, .... |
| CATTTCA_MIR | 0.0875 | 0.293 | 0.197822 | -0.365 | -1.391 | 30 | Mef2c, M.... |
| CBFA2T2_TAR | 0.9906 | 1 | 0.03509 | -0.125 | -0.6449 | 129 | Mef2c, G.... |
| CBX5_TARGET | 0.1748 | 0.431 | 0.160801 | 0.292 | 1.24503 | 43 | Krt8, Kr.... |
| CBX7_TARGET | 0.3126 | 0.616 | 0.095603 | -0.244 | -1.0904 | 59 | Shisa5, .... |

| | | | | | | | |
|---|---|---|---|---|---|---|---|
| CC2D1A_TARG | 0.9206 | 1 | 0.039841 | -0.152 | -0.7172 | 76 | Cd53, Sm.... |
| CCAWYNNGAA | 0.0759 | 0.267 | 0.248911 | 0.437 | 1.41919 | 16 | Sox4, Ct.... |
| CCCNNGGGAR | 0.5583 | 0.833 | 0.067207 | -0.254 | -0.9175 | 25 | Fchsd2, .... |
| CCTGCTG_MIR | 0.5271 | 0.808 | 0.069613 | -0.286 | -0.9575 | 20 | Bcl11a, .... |
| CDC5L_TARGE | 0.6719 | 0.903 | 0.057731 | -0.253 | -0.8591 | 21 | Arhgef1,.... |
| CDC5_01 | 0.0355 | 0.189 | 0.321776 | -0.465 | -1.5578 | 20 | Foxp1, M.... |
| CDC73_TARGE | 0.8537 | 1 | 0.046046 | -0.184 | -0.7468 | 40 | Mef2c, M.... |
| CDPCR3HD_01 | 0.0619 | 0.243 | 0.24134 | -0.417 | -1.457 | 23 | Fchsd2, .... |
| CDX2_Q5 | 0.9091 | 1 | 0.043888 | -0.201 | -0.6451 | 17 | Foxp1, Mbnl1 |
| CEBPA_01 | 0.3816 | 0.674 | 0.087052 | -0.291 | -1.0529 | 25 | Mbnl1, B.... |
| CEBPB_01 | 0.4471 | 0.744 | 0.094356 | 0.276 | 1.03444 | 27 | S100a8, .... |
| CEBPB_02 | 0.8232 | 0.981 | 0.048022 | -0.203 | -0.749 | 27 | Foxp1, B.... |
| CEBPDELTA_C | 0.534 | 0.812 | 0.084075 | 0.257 | 0.95462 | 26 | S100a9, .... |
| CEBPGAMMA_ | 0.047 | 0.219 | 0.276501 | -0.414 | -1.5089 | 26 | Mef2c, S.... |
| CEBPZ_TARGE | 1 | 1 | 0.057853 | 0.111 | 0.5613 | 101 | Atpif1, .... |
| CEBP_01 | 0.8003 | 0.976 | 0.048603 | -0.229 | -0.7463 | 18 | Foxp1, R.... |
| CEBP_C | 0.7741 | 0.964 | 0.051322 | -0.232 | -0.7785 | 20 | Foxp1, L.... |
| CEBP_Q2 | 0.1611 | 0.417 | 0.170932 | 0.382 | 1.28002 | 18 | Clu, S10.... |
| CEBP_Q2_01 | 0.5188 | 0.803 | 0.06912 | -0.294 | -0.9613 | 18 | Foxp1, M.... |
| CEBP_Q3 | 0.3678 | 0.667 | 0.088626 | -0.312 | -1.0736 | 22 | Bcl11a, .... |
| CETS1P54_01 | 0.1629 | 0.419 | 0.142057 | -0.346 | -1.2765 | 27 | Mef2c, I.... |
| CGTSACG_PAX | 0.7054 | 0.926 | 0.055135 | -0.253 | -0.8367 | 19 | Bcl11a, .... |
| CHAF1B_TARG | 0.4685 | 0.762 | 0.071479 | -0.201 | -0.987 | 93 | Dock2, A.... |
| CHOP_01 | 0.5744 | 0.846 | 0.078921 | 0.249 | 0.90314 | 23 | S100a9, .... |
| CIC_TARGET_( | 0.9609 | 1 | 0.040085 | -0.155 | -0.6231 | 38 | Smchd1, .... |
| CIITA_TARGET | 0.1048 | 0.325 | 0.172324 | -0.243 | -1.2502 | 128 | Mef2c, C.... |
| CIZ_01 | 0.6222 | 0.879 | 0.061036 | -0.265 | -0.8779 | 19 | Bcl11a, .... |
| CMYB_01 | 0.8104 | 0.979 | 0.048897 | -0.205 | -0.746 | 26 | Arhgap45.... |
| COREBINDING | 0.3693 | 0.667 | 0.088895 | -0.295 | -1.0651 | 25 | Il7r, Ar.... |
| COUP_01 | 0.9395 | 1 | 0.055479 | 0.184 | 0.6112 | 17 | Gapdh, P.... |
| COUP_DR1_Q6 | 0.965 | 1 | 0.041154 | -0.167 | -0.5342 | 17 | Shisa5 |
| CP2_01 | 0.8231 | 0.981 | 0.048506 | -0.219 | -0.7347 | 20 | Arhgap45.... |
| CP2_02 | 0.9862 | 1 | 0.05259 | 0.137 | 0.501 | 25 | Actn1, C.... |
| CREB3L4_TAR | 1 | 1 | 0.059476 | 0.094 | 0.50401 | 135 | Krt8, Ep.... |
| CREB3_TARGE | 0.5339 | 0.812 | 0.068633 | -0.259 | -0.9436 | 26 | Mef2c, B.... |
| CREBP1CJUN_ | 0.9039 | 1 | 0.05653 | 0.182 | 0.65846 | 23 | Cystm1, .... |
| CREBP1_Q2 | 0.4963 | 0.778 | 0.090543 | 0.263 | 0.98996 | 29 | Cystm1, .... |
| CREB_01 | 0.8077 | 0.977 | 0.061303 | 0.207 | 0.75485 | 24 | Cystm1, .... |
| CREB_02 | 0.4872 | 0.774 | 0.088626 | 0.273 | 0.97339 | 22 | Cldn7, T.... |
| CREB_Q2 | 0.2363 | 0.51 | 0.138022 | 0.306 | 1.17093 | 30 | Cystm1, .... |
| CREB_Q2_01 | 0.3753 | 0.67 | 0.103197 | 0.292 | 1.05691 | 23 | Cystm1, .... |
| CREB_Q3 | 0.7951 | 0.976 | 0.050092 | -0.22 | -0.7588 | 22 | Ebf1, Bc.... |
| CREB_Q4 | 0.8729 | 1 | 0.061303 | 0.185 | 0.69637 | 29 | Cystm1, .... |

| pathway | pval | padj | log2err | ES | NES | size | LeadingEdge |
|---|---|---|---|---|---|---|---|
| GCM_ACTG | 3.80E-01 | 5.57E-01 | 0.10396 | 0.24395 | 1.0483 | 45 | Slc25a3,.... |
| GCM_APEX | 7.34E-01 | 8.36E-01 | 0.06831 | 0.1976 | 0.8372 | 42 | Psmd8, S.... |
| GCM_CBFB | 9.88E-01 | 9.99E-01 | 0.05536 | 0.13103 | 0.4914 | 28 | Atxn10, .... |
| GCM_CSNK | 2.33E-01 | 4.08E-01 | 0.1396 | 0.2821 | 1.1703 | 37 | Ybx1, Ps.... |
| GCM_DDX5 | 5.22E-01 | 6.78E-01 | 0.06831 | -0.2777 | -0.972 | 23 | Fus, Bcl.... |
| GCM_DFFA | 8.86E-02 | 2.00E-01 | 0.19579 | -0.4557 | -1.422 | 15 | Rbm25, S.... |
| GCM_HDAC | 9.72E-01 | 9.98E-01 | 0.04058 | -0.1627 | -0.528 | 17 | Arhgef1 |
| GCM_MLL | 1.01E-02 | 3.11E-02 | 0.38073 | -0.4935 | -1.688 | 21 | Rbm25, A.... |
| GCM_MYST | 6.81E-02 | 1.63E-01 | 0.22497 | -0.468 | -1.461 | 15 | Rbm25, K.... |
| GCM_NF2 | 5.64E-01 | 7.12E-01 | 0.06538 | -0.28 | -0.93 | 19 | Luc7l2, .... |
| GCM_NPM1 | 1.88E-01 | 3.46E-01 | 0.15631 | 0.28424 | 1.2105 | 44 | Hint1, S.... |
| GCM_PFN1 | 9.66E-01 | 9.97E-01 | 0.03952 | -0.151 | -0.529 | 23 | Arhgef1,.... |
| GCM_PPP1 | 5.31E-01 | 6.86E-01 | 0.08554 | 0.27281 | 0.9536 | 21 | Slc25a3,.... |
| GCM_PSME | 9.30E-01 | 9.77E-01 | 0.05749 | 0.16185 | 0.6652 | 36 | Slc25a3,.... |
| GCM_RAB1 | 1.92E-01 | 3.49E-01 | 0.12879 | -0.3964 | -1.267 | 16 | Luc7l2, .... |
| GCM_TPT1 | 1.55E-01 | 2.95E-01 | 0.1752 | 0.33174 | 1.2497 | 27 | Slc25a3,.... |
| GCM_UBE2 | 1.63E-01 | 3.04E-01 | 0.14123 | -0.4101 | -1.311 | 16 | Zfp292, .... |
| GNF2_APEX | 9.95E-01 | 9.99E-01 | 0.03816 | -0.113 | -0.433 | 31 | Srsf7, F.... |
| GNF2_CASI | 7.39E-01 | 8.37E-01 | 0.05291 | -0.2421 | -0.793 | 18 | Samhd1, .... |
| GNF2_CD48 | 1.59E-06 | 1.59E-05 | 0.64355 | -0.7193 | -2.357 | 18 | Coro1a, .... |
| GNF2_CD53 | 4.46E-09 | 1.71E-07 | 0.76146 | -0.6841 | -2.618 | 31 | Coro1a, .... |
| GNF2_DAP3 | 6.11E-01 | 7.44E-01 | 0.07628 | 0.20381 | 0.9197 | 52 | Ifrd1, S.... |
| GNF2_DDX | 6.37E-04 | 2.93E-03 | 0.47727 | -0.5445 | -1.978 | 26 | Itga4, P.... |
| GNF2_DEK | 4.39E-01 | 6.12E-01 | 0.0783 | -0.3031 | -1.019 | 20 | Smc4, Cp.... |
| GNF2_DENI | 6.61E-01 | 7.78E-01 | 0.0581 | -0.257 | -0.864 | 20 | Fus, Cps.... |
| GNF2_EIF3 | 2.55E-01 | 4.32E-01 | 0.13146 | 0.2501 | 1.1331 | 54 | Eprs, Cd.... |
| GNF2_FBL | 5.74E-01 | 7.22E-01 | 0.08064 | 0.2058 | 0.9324 | 54 | Hint1, E.... |
| GNF2_GLTS | 3.10E-01 | 4.92E-01 | 0.09689 | -0.3647 | -1.138 | 15 | Rps27, A.... |
| GNF2_HDA | 2.44E-02 | 6.76E-02 | 0.35249 | -0.386 | -1.536 | 38 | Rbm39, S.... |
| GNF2_HLA_ | 9.79E-06 | 8.34E-05 | 0.59333 | -0.6564 | -2.263 | 22 | Coro1a, .... |
| GNF2_INPF | 5.80E-04 | 2.72E-03 | 0.47727 | -0.5889 | -1.98 | 20 | Coro1a, .... |
| GNF2_KPNI | 1.77E-01 | 3.28E-01 | 0.135 | -0.3832 | -1.272 | 19 | Srsf7, C.... |
| GNF2_MYD | 4.10E-03 | 1.43E-02 | 0.40702 | -0.5587 | -1.855 | 19 | Coro1a, .... |
| GNF2_NPM | 7.08E-01 | 8.19E-01 | 0.05313 | -0.23 | -0.828 | 25 | Cpsf6, E.... |
| GNF2_PTPI | 1.82E-08 | 4.19E-07 | 0.73376 | -0.7288 | -2.552 | 23 | Coro1a, .... |
| GNF2_PTPI | 1.38E-07 | 2.11E-06 | 0.69013 | -0.6893 | -2.504 | 26 | Coro1a, .... |
| GNF2_RAN | 1.48E-01 | 2.85E-01 | 0.17822 | 0.40349 | 1.3747 | 19 | Nme1, Se.... |
| GNF2_RAP | 2.62E-04 | 1.44E-03 | 0.49849 | -0.6463 | -2.065 | 16 | Ptprc, M.... |
| GNF2_RBBI | 1.33E-01 | 2.67E-01 | 0.15631 | -0.359 | -1.336 | 28 | Rbm25, R.... |
| GNF2_SELL | 9.45E-06 | 8.34E-05 | 0.59333 | -0.7089 | -2.301 | 17 | Coro1a, .... |
| GNF2_SERI | 6.44E-09 | 2.10E-07 | 0.76146 | 0.83644 | 2.6342 | 15 | Clps, Pn.... |
| GNF2_SPIN | 1.79E-09 | 8.24E-08 | 0.78819 | 0.84691 | 2.7361 | 16 | Reg1, Cl.... |

| | | | | | | | |
|---|---|---|---|---|---|---|---|
| GNF2_ST13 | 2.98E-01 | 4.78E-01 | 0.12211 | 0.30019 | 1.1309 | 27 | Snrpe, E.... |
| GNF2_STA1 | 7.83E-07 | 9.01E-06 | 0.65944 | -0.6153 | -2.367 | 33 | Coro1a, .... |
| GNF2_TDG | 2.62E-01 | 4.34E-01 | 0.10755 | -0.3684 | -1.177 | 16 | Cpsf6, P.... |
| GNF2_TPT1 | 5.80E-01 | 7.26E-01 | 0.06378 | -0.2848 | -0.91 | 16 | Rps27, R.... |
| GNF2_UBE2 | 4.30E-01 | 6.03E-01 | 0.09855 | 0.2883 | 1.0197 | 22 | Eprs, Se.... |
| GNF2_VAV1 | 2.00E-05 | 1.60E-04 | 0.57561 | -0.6781 | -2.251 | 19 | Coro1a, .... |
| GNF2_XRC | 7.73E-01 | 8.63E-01 | 0.06613 | 0.22338 | 0.7808 | 21 | Ifrd1, S.... |
| MODULE_1 | 9.41E-05 | 6.18E-04 | 0.53843 | 0.50954 | 2.1719 | 43 | Nupr1, K.... |
| MODULE_1 | 2.05E-04 | 1.24E-03 | 0.51885 | 0.53417 | 2.0674 | 31 | Spp1, Nu.... |
| MODULE_1 | 4.99E-03 | 1.69E-02 | 0.40702 | 0.53026 | 1.8755 | 22 | Sox4, An.... |
| MODULE_1 | 5.73E-03 | 1.91E-02 | 0.40702 | 0.44077 | 1.7634 | 34 | Spp1, Nu.... |
| MODULE_1 | 5.78E-02 | 1.45E-01 | 0.32178 | 0.24868 | 1.3257 | 123 | Fabp5, P.... |
| MODULE_1 | 4.46E-01 | 6.12E-01 | 0.09592 | 0.32283 | 1.0167 | 15 | Eif3b, E.... |
| MODULE_1 | 8.47E-02 | 1.95E-01 | 0.24134 | 0.37395 | 1.4087 | 27 | S100a9, .... |
| MODULE_1 | 2.52E-03 | 9.68E-03 | 0.43171 | 0.41072 | 1.836 | 51 | Spp1, Kr.... |
| MODULE_1 | 3.81E-06 | 3.50E-05 | 0.62726 | -0.6561 | -2.297 | 23 | Cd74, Ba.... |
| MODULE_1 | 1.24E-09 | 7.14E-08 | 0.78819 | 0.69301 | 2.8148 | 35 | Reg1, Sp.... |
| MODULE_1 | 3.20E-01 | 5.04E-01 | 0.09375 | -0.313 | -1.127 | 25 | Ets1, Sp.... |
| MODULE_1 | 5.60E-01 | 7.11E-01 | 0.08266 | 0.26507 | 0.9266 | 21 | Dap, Nme.... |
| MODULE_1 | 2.40E-01 | 4.19E-01 | 0.13802 | 0.33871 | 1.198 | 22 | Reg1, Dm.... |
| MODULE_1 | 1.93E-01 | 3.49E-01 | 0.12879 | -0.298 | -1.21 | 42 | Itga4, I.... |
| MODULE_1 | 2.05E-04 | 1.24E-03 | 0.51885 | 0.53417 | 2.0674 | 31 | Spp1, Nu.... |
| MODULE_1 | 1.98E-02 | 5.54E-02 | 0.35249 | -0.516 | -1.611 | 15 | Ighm, Rc.... |
| MODULE_1 | 4.07E-01 | 5.89E-01 | 0.10135 | 0.33607 | 1.0584 | 15 | Marcks, .... |
| MODULE_1 | 2.62E-01 | 4.34E-01 | 0.12879 | 0.35169 | 1.174 | 17 | Eif3b, U.... |
| MODULE_1 | 1.25E-03 | 5.40E-03 | 0.45506 | -0.3959 | -1.789 | 67 | Cd37, Ig.... |
| MODULE_1 | 8.64E-02 | 1.97E-01 | 0.24891 | 0.24086 | 1.2744 | 116 | Fabp5, P.... |
| MODULE_1 | 8.31E-07 | 9.10E-06 | 0.65944 | 0.56469 | 2.4265 | 45 | Gpx4, Co.... |
| MODULE_1 | 6.85E-01 | 7.96E-01 | 0.07307 | 0.22493 | 0.8435 | 28 | Rab11a, .... |
| MODULE_1 | 2.97E-01 | 4.78E-01 | 0.10063 | -0.2635 | -1.113 | 50 | Coro1a, .... |
| MODULE_1 | 3.24E-02 | 8.57E-02 | 0.32178 | 0.47309 | 1.5793 | 17 | Reg1, Sp.... |
| MODULE_1 | 9.69E-02 | 2.14E-01 | 0.22206 | 0.29499 | 1.3365 | 54 | Krt18, T.... |
| MODULE_1 | 3.10E-05 | 2.38E-04 | 0.55733 | -0.618 | -2.225 | 25 | Ighm, Cd.... |
| MODULE_1 | 3.35E-03 | 1.22E-02 | 0.43171 | -0.5035 | -1.829 | 26 | Ptprc, C.... |
| MODULE_1 | 3.28E-01 | 5.13E-01 | 0.09375 | -0.2788 | -1.095 | 36 | Itga4, I.... |
| MODULE_1 | 1.60E-02 | 4.66E-02 | 0.35249 | 0.38787 | 1.6176 | 38 | Spp1, Kr.... |
| MODULE_1 | 3.54E-06 | 3.39E-05 | 0.62726 | 0.73665 | 2.4868 | 18 | Clu, Krt.... |
| MODULE_1 | 3.68E-01 | 5.46E-01 | 0.08705 | -0.3474 | -1.085 | 15 | Peli1, A.... |
| MODULE_1 | 2.18E-01 | 3.89E-01 | 0.11882 | -0.3342 | -1.214 | 26 | Rbm5, Ce.... |
| MODULE_1 | 2.70E-04 | 1.44E-03 | 0.49849 | -0.5977 | -2.045 | 21 | Ighm, Rc.... |
| MODULE_1 | 9.11E-02 | 2.03E-01 | 0.23113 | 0.34258 | 1.3705 | 34 | Nupr1, C.... |
| MODULE_1 | 4.66E-01 | 6.27E-01 | 0.09168 | 0.29639 | 0.9894 | 17 | Marcks, .... |
| MODULE_1 | 3.63E-01 | 5.43E-01 | 0.10925 | 0.30069 | 1.0771 | 23 | Sox4, So.... |

| | | | | | | | |
|---|---|---|---|---|---|---|---|
| MODULE_1 | 6.19E-01 | 7.46E-01 | 0.07768 | 0.20722 | 0.906 | 48 | Krt19, S.... |
| MODULE_2 | 1.67E-03 | 6.73E-03 | 0.45506 | 0.42627 | 1.8317 | 45 | Spp1, Nu.... |
| MODULE_2 | 2.02E-01 | 3.63E-01 | 0.12503 | -0.3909 | -1.249 | 16 | Mbnl1, S.... |
| MODULE_2 | 5.24E-05 | 3.65E-04 | 0.55733 | -0.6321 | -2.162 | 21 | Ighm, Cd.... |
| MODULE_2 | 1.17E-07 | 2.11E-06 | 0.70498 | 0.63132 | 2.6192 | 37 | Nupr1, T.... |
| MODULE_2 | 1.65E-03 | 6.73E-03 | 0.45506 | 0.56989 | 1.9746 | 20 | Ndufc1, .... |
| MODULE_2 | 4.84E-02 | 1.22E-01 | 0.27129 | -0.4579 | -1.54 | 20 | Cd79a, H.... |
| MODULE_2 | 7.25E-03 | 2.32E-02 | 0.40702 | 0.48847 | 1.8284 | 26 | Krt18, N.... |
| MODULE_2 | 2.67E-04 | 1.44E-03 | 0.49849 | -0.5776 | -2.08 | 25 | Ighm, Rc.... |
| MODULE_2 | 1.59E-01 | 3.00E-01 | 0.14376 | -0.3922 | -1.285 | 18 | Mbnl1, S.... |
| MODULE_2 | 2.80E-07 | 3.39E-06 | 0.67496 | 0.68641 | 2.5741 | 28 | S100a9, .... |
| MODULE_2 | 9.19E-01 | 9.72E-01 | 0.04208 | -0.1709 | -0.628 | 27 | Lck, Tap.... |
| MODULE_2 | 7.14E-01 | 8.21E-01 | 0.07028 | 0.20235 | 0.8439 | 38 | Sox4, So.... |
| MODULE_2 | 4.48E-01 | 6.12E-01 | 0.07687 | -0.2639 | -1.016 | 33 | Itga4, A.... |
| MODULE_2 | 1.13E-01 | 2.39E-01 | 0.17232 | -0.4113 | -1.365 | 19 | Ankrd44,.... |
| MODULE_2 | 1.32E-07 | 2.11E-06 | 0.69013 | -0.7712 | -2.503 | 17 | Cd79a, H.... |
| MODULE_2 | 2.66E-01 | 4.36E-01 | 0.12944 | 0.33313 | 1.1644 | 21 | Ybx1, Nd.... |
| MODULE_2 | 1.25E-01 | 2.57E-01 | 0.19579 | 0.43411 | 1.3671 | 15 | Psmd8, P.... |
| MODULE_2 | 1.29E-07 | 2.11E-06 | 0.69013 | -0.6603 | -2.486 | 30 | Cd37, Ig.... |
| MODULE_3 | 6.00E-02 | 1.49E-01 | 0.28201 | 0.30955 | 1.4182 | 57 | Krt18, K.... |
| MODULE_3 | 2.33E-04 | 1.37E-03 | 0.51885 | -0.5923 | -2.042 | 22 | Ighm, Rc.... |
| MODULE_3 | 4.22E-01 | 5.95E-01 | 0.0802 | -0.2773 | -1.044 | 30 | Itga4, A.... |
| MODULE_3 | 9.89E-01 | 9.99E-01 | 0.03676 | -0.1247 | -0.586 | 81 | Rhoh, R3.... |
| MODULE_3 | 8.31E-02 | 1.93E-01 | 0.24134 | 0.36462 | 1.4104 | 30 | Krt8, Kr.... |
| MODULE_3 | 1.27E-01 | 2.58E-01 | 0.19381 | 0.42743 | 1.3809 | 16 | Pnliprp1.... |
| MODULE_3 | 8.08E-09 | 2.10E-07 | 0.74774 | 0.7482 | 2.7344 | 25 | Clu, Reg.... |
| MODULE_3 | 2.80E-08 | 5.86E-07 | 0.73376 | -0.6934 | -2.549 | 27 | Cd37, Ig.... |
| MODULE_3 | 4.64E-01 | 6.27E-01 | 0.09315 | 0.30956 | 1.0001 | 16 | Marcks, .... |
| MODULE_3 | 2.02E-05 | 1.60E-04 | 0.57561 | -0.6436 | -2.219 | 22 | Cd37, Ig.... |
| MODULE_3 | 1.46E-02 | 4.31E-02 | 0.38073 | 0.53887 | 1.6971 | 15 | Ybx1, Nd.... |
| MODULE_3 | 2.28E-01 | 4.03E-01 | 0.11623 | -0.2934 | -1.179 | 40 | Igkc, Ar.... |
| MODULE_3 | 4.22E-01 | 5.95E-01 | 0.0802 | -0.2773 | -1.044 | 30 | Itga4, A.... |
| MODULE_3 | 1.76E-07 | 2.53E-06 | 0.69013 | 0.66547 | 2.5741 | 30 | S100a9, .... |
| MODULE_4 | 2.59E-01 | 4.34E-01 | 0.13011 | 0.34035 | 1.1792 | 20 | Reg1, Dm.... |
| MODULE_4 | 1.96E-03 | 7.63E-03 | 0.43171 | 0.5628 | 1.95 | 20 | Cox5b, U.... |
| MODULE_4 | 8.20E-09 | 2.10E-07 | 0.74774 | -0.7019 | -2.613 | 28 | Cd37, Ig.... |
| MODULE_4 | 3.21E-05 | 2.39E-04 | 0.55733 | -0.4272 | -2.004 | 80 | Cd37, Ig.... |
| MODULE_4 | 2.67E-07 | 3.39E-06 | 0.67496 | -0.508 | -2.324 | 70 | Cd37, Ig.... |
| MODULE_4 | 4.22E-01 | 5.95E-01 | 0.0802 | -0.2773 | -1.044 | 30 | Itga4, A.... |
| MODULE_4 | 4.38E-04 | 2.14E-03 | 0.49849 | -0.4576 | -1.906 | 46 | Cd79a, H.... |
| MODULE_4 | 6.78E-04 | 3.06E-03 | 0.47727 | 0.6108 | 2.0389 | 17 | Col1a2, .... |
| MODULE_5 | 2.68E-04 | 1.44E-03 | 0.49849 | 0.42618 | 1.9867 | 61 | S100a9, .... |
| MODULE_5 | 1.91E-07 | 2.58E-06 | 0.69013 | 0.56846 | 2.5411 | 51 | Spp1, Kr.... |

| | | | | | | | |
|---|---|---|---|---|---|---|---|
| MODULE_5 | 2.58E-03 | 9.71E-03 | 0.43171 | -0.4192 | -1.736 | 45 | Cd37, Ig.... |
| MODULE_5 | 5.91E-01 | 7.34E-01 | 0.06239 | -0.2488 | -0.904 | 26 | Arhgap45.... |
| MODULE_5 | 6.63E-01 | 7.78E-01 | 0.07418 | 0.26705 | 0.841 | 15 | Cdh1, Pr.... |
| MODULE_5 | 4.21E-13 | 9.67E-11 | 0.9326 | 0.65048 | 2.9888 | 59 | Reg1, S1.... |
| MODULE_6 | 2.32E-10 | 2.67E-08 | 0.82666 | 0.62683 | 2.849 | 53 | S100a9, .... |
| MODULE_6 | 1.76E-02 | 5.06E-02 | 0.35249 | -0.4112 | -1.639 | 39 | Cd37, Ig.... |
| MODULE_6 | 1.26E-06 | 1.32E-05 | 0.64355 | 0.59459 | 2.4797 | 38 | Ndufc1, .... |
| MODULE_6 | 4.16E-04 | 2.13E-03 | 0.49849 | -0.5003 | -1.965 | 36 | Cd37, Cd.... |
| MODULE_6 | 4.27E-04 | 2.13E-03 | 0.49849 | 0.52736 | 2.0735 | 32 | Spp1, Nu.... |
| MODULE_6 | 1.16E-01 | 2.42E-01 | 0.17093 | -0.3107 | -1.299 | 47 | Itga4, I.... |
| MODULE_7 | 6.34E-01 | 7.60E-01 | 0.07707 | 0.22823 | 0.8833 | 31 | Col1a2, .... |
| MODULE_7 | 1.40E-03 | 5.94E-03 | 0.45506 | -0.4175 | -1.763 | 50 | Cd79a, H.... |
| MODULE_8 | 4.55E-05 | 3.27E-04 | 0.55733 | 0.46388 | 2.1163 | 56 | Krt18, N.... |
| MODULE_8 | 1.02E-01 | 2.24E-01 | 0.22799 | 0.23454 | 1.2494 | 122 | Prdx1, D.... |
| MODULE_8 | 8.32E-05 | 5.63E-04 | 0.53843 | -0.4089 | -1.924 | 82 | Cd37, Ig.... |
| MODULE_8 | 3.56E-10 | 2.73E-08 | 0.81404 | 0.58964 | 2.7807 | 63 | Reg1, S1.... |
| MODULE_9 | 5.52E-01 | 7.06E-01 | 0.08313 | 0.27431 | 0.9346 | 19 | Psmd8, P.... |
| MODULE_9 | 1.07E-04 | 6.84E-04 | 0.53843 | 0.55487 | 2.1816 | 32 | Prdx1, G.... |
| MODULE_9 | 3.24E-02 | 8.57E-02 | 0.32178 | 0.47309 | 1.5793 | 17 | Reg1, Sp.... |
| MODULE_9 | 2.45E-01 | 4.23E-01 | 0.11192 | -0.3766 | -1.204 | 16 | Peli1, W.... |
| MODULE_9 | 9.67E-01 | 9.97E-01 | 0.05502 | 0.14249 | 0.672 | 63 | Krt19, S.... |
| MORF_AAT | 3.60E-01 | 5.43E-01 | 0.10839 | 0.25782 | 1.0775 | 39 | Atxn10, .... |
| MORF_ACP | 4.48E-01 | 6.12E-01 | 0.09689 | 0.21246 | 1.0012 | 65 | Ybx1, At.... |
| MORF_ACT | 4.50E-01 | 6.12E-01 | 0.09375 | 0.21499 | 0.9878 | 59 | Tagln2, .... |
| MORF_ANP | 1.54E-01 | 2.95E-01 | 0.17822 | 0.26278 | 1.2443 | 67 | Ybx1, At.... |
| MORF_AP2 | 5.16E-04 | 2.47E-03 | 0.47727 | 0.39713 | 1.884 | 69 | Sec61b, .... |
| MORF_AP3 | 7.86E-02 | 1.84E-01 | 0.24891 | 0.33092 | 1.383 | 39 | Ndufc1, .... |
| MORF_ATC | 1.12E-03 | 4.96E-03 | 0.45506 | 0.50454 | 1.9516 | 30 | Sec61g, .... |
| MORF_ATR | 3.15E-02 | 8.57E-02 | 0.32178 | -0.4802 | -1.574 | 18 | Zfp292, .... |
| MORF_BAG | 5.50E-01 | 7.06E-01 | 0.08336 | 0.27444 | 0.935 | 19 | Vcp, Rab.... |
| MORF_BEC | 9.10E-01 | 9.72E-01 | 0.04225 | -0.1774 | -0.621 | 23 | Arhgef1,.... |
| MORF_BMI | 9.95E-01 | 9.99E-01 | 0.03856 | -0.1356 | -0.423 | 15 | Atp11b, .... |
| MORF_BUB | 3.63E-01 | 5.43E-01 | 0.10839 | 0.22184 | 1.036 | 62 | Ndufc1, .... |
| MORF_CCN | 2.98E-01 | 4.78E-01 | 0.12268 | 0.28836 | 1.116 | 31 | Ybx1, Hi.... |
| MORF_CDC | 3.75E-01 | 5.53E-01 | 0.08554 | -0.2867 | -1.067 | 28 | Atp11b, .... |
| MORF_CDK | 7.79E-01 | 8.65E-01 | 0.04979 | -0.2438 | -0.761 | 15 | Fus, Hnr.... |
| MORF_CSN | 1.87E-03 | 7.43E-03 | 0.45506 | 0.36321 | 1.7761 | 77 | Dap, Gpx.... |
| MORF_CTB | 1.13E-01 | 2.39E-01 | 0.20429 | 0.28918 | 1.3102 | 54 | Cox5b, H.... |
| MORF_CUL | 4.80E-01 | 6.38E-01 | 0.09168 | 0.31191 | 0.9823 | 15 | Ywhaq, E.... |
| MORF_DAP | 3.19E-02 | 8.57E-02 | 0.32178 | 0.47839 | 1.615 | 18 | Dap, Gpx.... |
| MORF_DAP | 9.85E-03 | 3.06E-02 | 0.38073 | 0.34551 | 1.5875 | 59 | Sec61b, .... |
| MORF_DDE | 3.50E-01 | 5.36E-01 | 0.11146 | 0.2757 | 1.084 | 32 | Glud1, Y.... |
| MORF_DEK | 7.87E-01 | 8.70E-01 | 0.06879 | 0.1694 | 0.8404 | 84 | Sec61b, .... |

| pathway | pval | padj | log2err | ES | NES | size | LeadingEdge |
|---|---|---|---|---|---|---|---|
| GOBP_ACTIN_FILA | 8.34E-01 | 9.58E-01 | 0.0456 | -0.17041 | -0.7936 | 81 | Coro1a, .... |
| GOBP_ACTIN_FILA | 5.66E-01 | 8.12E-01 | 0.0806 | 0.28099 | 0.93462 | 17 | Marcks, .... |
| GOBP_ACTIN_FILA | 9.41E-01 | 9.90E-01 | 0.0393 | -0.15278 | -0.66159 | 55 | Coro1a, .... |
| GOBP_ACTIN_FILA | 4.21E-01 | 7.02E-01 | 0.0804 | -0.29134 | -1.0337 | 24 | Coro1a, .... |
| GOBP_ACTIN_POL | 8.01E-01 | 9.48E-01 | 0.0489 | -0.2108 | -0.76542 | 26 | Coro1a, .... |
| GOBP_ACTIVATION | 8.88E-07 | 4.31E-05 | 0.6594 | -0.56134 | -2.32218 | 46 | Mef2c, I.... |
| GOBP_ACTOMYOS | 9.34E-01 | 9.89E-01 | 0.0547 | 0.170754 | 0.59165 | 19 | Krt19, T.... |
| GOBP_ADAPTIVE_ | 7.17E-07 | 4.11E-05 | 0.6594 | -0.56345 | -2.33093 | 46 | Mef2c, I.... |
| GOBP_ADAPTIVE_ | 7.48E-06 | 1.90E-04 | 0.6105 | -0.598 | -2.2534 | 30 | Mef2c, I.... |
| GOBP_ADP_METAI | 2.06E-01 | 4.82E-01 | 0.1464 | 0.396375 | 1.26169 | 15 | Nupr1, G.... |
| GOBP_AEROBIC_F | 5.61E-06 | 1.54E-04 | 0.6105 | 0.5516 | 2.35584 | 44 | Nupr1, N.... |
| GOBP_AGING | 3.18E-01 | 6.13E-01 | 0.1167 | 0.289993 | 1.09013 | 28 | Fos, Ybx.... |
| GOBP_ALCOHOL_I | 4.59E-01 | 7.34E-01 | 0.092 | 0.308509 | 1.00038 | 16 | Cel, App.... |
| GOBP_ALPHA_BET | 1.09E-03 | 1.08E-02 | 0.4551 | -0.56022 | -1.9325 | 22 | Malt1, H.... |
| GOBP_ALTERNATI | 5.87E-02 | 2.26E-01 | 0.2489 | -0.444 | -1.46096 | 19 | Mbnl1, R.... |
| GOBP_AMEBOIDAL | 1.17E-01 | 3.48E-01 | 0.1669 | -0.3253 | -1.32145 | 40 | Mef2c, E.... |
| GOBP_AMIDE_BIO | 9.82E-01 | 9.96E-01 | 0.0585 | 0.122903 | 0.65179 | 121 | Sox4, Rp.... |
| GOBP_AMIDE_TRA | 1.49E-01 | 4.00E-01 | 0.1752 | 0.394734 | 1.31295 | 17 | S100a8, .... |
| GOBP_ANATOMICA | 9.87E-02 | 3.21E-01 | 0.2311 | 0.249709 | 1.26308 | 96 | Krt19, S.... |
| GOBP_ANATOMICA | 2.67E-02 | 1.22E-01 | 0.3525 | 0.446289 | 1.6007 | 23 | Spp1, Tf.... |
| GOBP_ANIMAL_OF | 2.16E-01 | 4.99E-01 | 0.1473 | 0.258951 | 1.17002 | 53 | Sox4, Co.... |
| GOBP_ANION_TRA | 9.16E-01 | 9.83E-01 | 0.0568 | 0.18024 | 0.59951 | 17 | Anxa1, C.... |
| GOBP_ANTIGEN_P | 2.70E-04 | 3.42E-03 | 0.4985 | -0.61385 | -2.04966 | 20 | Cd74, H2.... |
| GOBP_ANTIGEN_P | 1.55E-04 | 2.22E-03 | 0.5188 | -0.65985 | -2.10487 | 16 | Cd74, H2.... |
| GOBP_ANTIGEN_R | 1.24E-11 | 7.02E-09 | 0.8753 | -0.70976 | -2.81352 | 36 | Mef2c, I.... |
| GOBP_ANTIMICRO | 1.43E-05 | 3.19E-04 | 0.5933 | 0.708142 | 2.3554 | 17 | Reg3b, R.... |
| GOBP_APOPTOTIC | 1.20E-02 | 6.50E-02 | 0.3807 | 0.326268 | 1.56683 | 73 | Clu, S10.... |
| GOBP_ATP_METAE | 1.13E-05 | 2.64E-04 | 0.5933 | 0.467376 | 2.16097 | 61 | Nupr1, A.... |
| GOBP_ATP_SYNTH | 3.66E-04 | 4.38E-03 | 0.4985 | 0.533648 | 2.04974 | 31 | Ndufc1, .... |
| GOBP_AXON_DEVI | 3.13E-01 | 6.07E-01 | 0.1215 | 0.272333 | 1.09397 | 36 | S100a6, .... |
| GOBP_BEHAVIOR | 9.47E-01 | 9.90E-01 | 0.0399 | -0.16881 | -0.6271 | 29 | Mef2c, B.... |
| GOBP_BIOLOGICA | 1.77E-03 | 1.55E-02 | 0.4551 | 0.296419 | 1.63386 | 140 | Reg3b, S.... |
| GOBP_BIOLOGICA | 6.35E-01 | 8.53E-01 | 0.0613 | -0.26762 | -0.88061 | 19 | Cd74, H2.... |
| GOBP_BIOLOGICA | 3.72E-02 | 1.59E-01 | 0.3218 | 0.244893 | 1.37086 | 149 | Reg3b, C.... |
| GOBP_BIOLOGICA | 1.41E-01 | 3.92E-01 | 0.1882 | 0.336407 | 1.30441 | 32 | Reg3g, C.... |
| GOBP_BLOOD_VES | 6.63E-01 | 8.74E-01 | 0.0552 | -0.20058 | -0.88654 | 62 | Ets1, Sp.... |
| GOBP_BONE_DEVI | 4.55E-01 | 7.29E-01 | 0.0771 | -0.31893 | -1.01736 | 16 | Mef2c, P.... |
| GOBP_B_CELL_AC | 7.27E-09 | 8.83E-07 | 0.7615 | -0.63945 | -2.60589 | 41 | Mef2c, I.... |
| GOBP_B_CELL_DII | 9.09E-04 | 9.30E-03 | 0.4773 | -0.58818 | -1.96394 | 20 | Cd79a, M.... |
| GOBP_B_CELL_ME | 4.19E-04 | 4.88E-03 | 0.4985 | -0.60471 | -2.01913 | 20 | Ighm, Cd.... |
| GOBP_B_CELL_PR | 8.61E-07 | 4.31E-05 | 0.6594 | -0.77148 | -2.40848 | 15 | Mef2c, C.... |
| GOBP_B_CELL_RE | 9.48E-09 | 1.01E-06 | 0.7477 | -0.76396 | -2.55087 | 20 | Mef2c, I.... |

| | | | | | | | |
|---|---|---|---|---|---|---|---|
| GOBP_CALCIUM_I( | 3.89E-01 | 6.77E-01 | 0.0865 | -0.31947 | -1.0667 | 20 | Coro1a, .... |
| GOBP_CALCIUM_I( | 6.77E-02 | 2.47E-01 | 0.2221 | -0.37238 | -1.43033 | 32 | Coro1a, .... |
| GOBP_CALCIUM_I( | 1.20E-02 | 6.50E-02 | 0.3807 | -0.54298 | -1.69512 | 15 | Coro1a, .... |
| GOBP_CALCIUM_M | 4.01E-01 | 6.87E-01 | 0.0841 | -0.3409 | -1.06425 | 15 | Ptprc, C.... |
| GOBP_CAMERA_T | 3.18E-01 | 6.13E-01 | 0.1148 | 0.345101 | 1.11903 | 16 | Sox9, Ju.... |
| GOBP_CANONICAL | 3.14E-01 | 6.08E-01 | 0.1152 | 0.311662 | 1.09924 | 21 | Sox4, So.... |
| GOBP_CARBOHYD | 3.00E-01 | 5.95E-01 | 0.1183 | 0.358918 | 1.14246 | 15 | Nupr1, G.... |
| GOBP_CARBOHYD | 9.65E-01 | 9.94E-01 | 0.0574 | 0.132931 | 0.57826 | 46 | Nme2, Vc.... |
| GOBP_CARBOHYD | 6.36E-01 | 8.53E-01 | 0.0781 | 0.185535 | 0.88686 | 71 | Nupr1, C.... |
| GOBP_CARBOHYD | 6.65E-01 | 8.75E-01 | 0.0755 | 0.216441 | 0.86194 | 34 | Nupr1, F.... |
| GOBP_CATION_TR | 8.64E-01 | 9.70E-01 | 0.0641 | 0.170054 | 0.78719 | 62 | Anxa2, N.... |
| GOBP_CATION_TR | 6.20E-01 | 8.44E-01 | 0.0591 | -0.19338 | -0.912 | 87 | Mef2c, C.... |
| GOBP_CELLULAR_ | 8.14E-01 | 9.51E-01 | 0.0672 | 0.15621 | 0.86306 | 142 | Clu, Sox.... |
| GOBP_CELLULAR_ | 1.31E-01 | 3.75E-01 | 0.1938 | 0.293223 | 1.26729 | 45 | Try5, Pr.... |
| GOBP_CELLULAR_ | 2.40E-01 | 5.34E-01 | 0.1404 | 0.247629 | 1.1377 | 57 | S100a6, .... |
| GOBP_CELLULAR_ | 7.20E-02 | 2.57E-01 | 0.2664 | 0.292821 | 1.38012 | 68 | S100a9, .... |
| GOBP_CELLULAR_ | 9.25E-02 | 3.09E-01 | 0.2344 | 0.312228 | 1.36279 | 48 | S100a9, .... |
| GOBP_CELLULAR_ | 8.21E-01 | 9.53E-01 | 0.0466 | -0.18562 | -0.75645 | 41 | Cd74, Et.... |
| GOBP_CELLULAR_ | 1.00E+00 | 1.00E+00 | 0.0592 | 0.078238 | 0.44232 | 165 | S100a11,.... |
| GOBP_CELLULAR_ | 4.44E-01 | 7.16E-01 | 0.0989 | 0.181191 | 1.00803 | 144 | Clu, Krt.... |
| GOBP_CELLULAR_ | 1.26E-01 | 3.64E-01 | 0.1596 | -0.30364 | -1.28002 | 49 | Samhd1, .... |
| GOBP_CELLULAR_ | 9.04E-06 | 2.19E-04 | 0.5933 | 0.693503 | 2.40292 | 19 | S100a9, .... |
| GOBP_CELLULAR_ | 9.63E-01 | 9.94E-01 | 0.054 | 0.167311 | 0.59011 | 21 | Spink1, .... |
| GOBP_CELLULAR_ | 2.46E-01 | 5.40E-01 | 0.1396 | 0.237384 | 1.14859 | 79 | Clu, Nup.... |
| GOBP_CELLULAR_ | 5.61E-06 | 1.54E-04 | 0.6105 | 0.5516 | 2.35584 | 44 | Nupr1, N.... |
| GOBP_CELLULAR_ | 4.02E-01 | 6.87E-01 | 0.104 | 0.27304 | 1.03203 | 29 | Nedd4, S.... |
| GOBP_CELLULAR_ | 2.06E-01 | 4.82E-01 | 0.1256 | -0.4031 | -1.25842 | 15 | Mef2c, M.... |
| GOBP_CELLULAR_ | 4.86E-03 | 3.34E-02 | 0.407 | 0.422844 | 1.71075 | 37 | Anxa1, P.... |
| GOBP_CELLULAR_ | 5.16E-01 | 7.72E-01 | 0.0674 | -0.22237 | -0.97346 | 58 | Cd74, Sh.... |
| GOBP_CELLULAR_ | 3.39E-01 | 6.28E-01 | 0.1162 | 0.208374 | 1.06503 | 100 | Ctsl, An.... |
| GOBP_CELLULAR_ | 7.24E-01 | 9.05E-01 | 0.0705 | 0.225404 | 0.81677 | 25 | Sox9, Co.... |
| GOBP_CELLULAR_ | 7.36E-01 | 9.11E-01 | 0.0538 | -0.23786 | -0.81061 | 21 | Itga4, P.... |
| GOBP_CELLULAR_ | 8.67E-01 | 9.70E-01 | 0.0648 | 0.16866 | 0.77055 | 60 | Ctsl, An.... |
| GOBP_CELLULAR_ | 1.47E-01 | 3.97E-01 | 0.1737 | 0.377017 | 1.33273 | 20 | Mt1, Mt2.... |
| GOBP_CELLULAR_ | 2.59E-01 | 5.56E-01 | 0.111 | -0.35192 | -1.17506 | 20 | Stat1, C.... |
| GOBP_CELLULAR_ | 2.47E-01 | 5.40E-01 | 0.1128 | -0.37739 | -1.20383 | 16 | Sp100, S.... |
| GOBP_CELLULAR_ | 9.54E-02 | 3.15E-01 | 0.228 | 0.306743 | 1.37594 | 52 | Clu, Spp.... |
| GOBP_CELLULAR_ | 2.06E-01 | 4.82E-01 | 0.1256 | -0.4031 | -1.25842 | 15 | Mef2c, M.... |
| GOBP_CELLULAR_ | 7.29E-01 | 9.07E-01 | 0.0523 | -0.19749 | -0.84153 | 53 | Mef2c, I.... |
| GOBP_CELLULAR_ | 8.56E-02 | 2.90E-01 | 0.2413 | 0.311965 | 1.39936 | 52 | Clu, Spp.... |
| GOBP_CELLULAR_ | 1.90E-02 | 9.53E-02 | 0.3525 | 0.294765 | 1.47808 | 93 | Clu, Spp.... |
| GOBP_CELLULAR_ | 9.70E-01 | 9.96E-01 | 0.0539 | 0.166608 | 0.54025 | 16 | P4hb, Lm.... |
| GOBP_CELLULAR_ | 2.19E-01 | 5.04E-01 | 0.1167 | -0.30914 | -1.17856 | 31 | Itga4, S.... |

| | | | | | | | |
|---|---|---|---|---|---|---|---|
| GOBP_CELLULAR_ | 4.81E-01 | 7.50E-01 | 0.0733 | -0.27309 | -0.99159 | 26 | Stat1, C.... |
| GOBP_CELLULAR_ | 3.22E-04 | 4.00E-03 | 0.4985 | 0.657542 | 2.093 | 15 | Anxa1, P.... |
| GOBP_CELLULAR_ | 6.29E-01 | 8.50E-01 | 0.0793 | 0.234779 | 0.89215 | 30 | Anxa1, N.... |
| GOBP_CELLULAR_ | 1.68E-01 | 4.33E-01 | 0.1723 | 0.206309 | 1.16043 | 152 | Clu, Spp.... |
| GOBP_CELLULAR_ | 3.71E-05 | 6.78E-04 | 0.5573 | 0.657 | 2.31725 | 21 | S100a9, .... |
| GOBP_CELL_ACTI\ | 5.59E-09 | 7.72E-07 | 0.7615 | -0.45804 | -2.31325 | 126 | Mef2c, I.... |
| GOBP_CELL_ACTI\ | 1.52E-04 | 2.19E-03 | 0.5188 | -0.5615 | -2.07176 | 28 | Malt1, C.... |
| GOBP_CELL_CELL_ | 1.73E-02 | 8.90E-02 | 0.3525 | -0.30223 | -1.46526 | 96 | Malt1, C.... |
| GOBP_CELL_CELL_ | 2.86E-02 | 1.29E-01 | 0.3525 | 0.496143 | 1.60881 | 16 | Cdh1, Cl.... |
| GOBP_CELL_CELL_ | 4.68E-02 | 1.90E-01 | 0.3218 | 0.291167 | 1.40882 | 79 | S100a9, .... |
| GOBP_CELL_CELL_ | 3.01E-01 | 5.95E-01 | 0.1244 | 0.291853 | 1.10903 | 30 | Sox4, So.... |
| GOBP_CELL_CHEM | 1.33E-01 | 3.78E-01 | 0.1574 | -0.38056 | -1.33854 | 23 | Cd74, Co.... |
| GOBP_CELL_CYCL | 8.06E-01 | 9.49E-01 | 0.0461 | -0.16698 | -0.83188 | 119 | Ptprc, B.... |
| GOBP_CELL_CYCL | 9.32E-01 | 9.89E-01 | 0.0423 | -0.1784 | -0.61538 | 22 | Ccnd3, B.... |
| GOBP_CELL_CYCL | 8.68E-01 | 9.70E-01 | 0.0617 | 0.178153 | 0.71507 | 35 | Anxa1, R.... |
| GOBP_CELL_CYCL | 5.50E-01 | 8.01E-01 | 0.0657 | -0.20742 | -0.956 | 77 | Ptprc, C.... |
| GOBP_CELL_DIVIS | 3.06E-01 | 6.01E-01 | 0.0969 | -0.27831 | -1.11909 | 39 | Lef1, Cc.... |
| GOBP_CELL_GRO\ | 7.27E-02 | 2.58E-01 | 0.2664 | 0.318103 | 1.38573 | 47 | S100a9, .... |
| GOBP_CELL_JUNC | 1.09E-01 | 3.37E-01 | 0.2139 | 0.357753 | 1.35223 | 29 | Cdh1, Cl.... |
| GOBP_CELL_JUNC | 1.08E-01 | 3.36E-01 | 0.2165 | 0.299893 | 1.3064 | 47 | Nedd4, C.... |
| GOBP_CELL_KILLI | 7.78E-03 | 4.74E-02 | 0.407 | -0.5228 | -1.74563 | 20 | Coro1a, .... |
| GOBP_CELL_MATF | 8.80E-01 | 9.73E-01 | 0.0437 | -0.19133 | -0.68984 | 25 | Itga4, C.... |
| GOBP_CELL_MIGR | 7.75E-02 | 2.72E-01 | 0.2043 | -0.25569 | -1.2881 | 124 | Mef2c, C.... |
| GOBP_CELL_MORI | 2.51E-01 | 5.44E-01 | 0.1373 | 0.228574 | 1.1303 | 87 | Clu, S10.... |
| GOBP_CELL_MORI | 3.94E-01 | 6.81E-01 | 0.1084 | 0.22528 | 1.02923 | 60 | S100a6, .... |
| GOBP_CELL_MORI | 7.00E-01 | 8.98E-01 | 0.0727 | 0.202817 | 0.84245 | 40 | S100a6, .... |
| GOBP_CELL_PART | 6.68E-01 | 8.78E-01 | 0.0759 | 0.201111 | 0.88387 | 49 | S100a6, .... |
| GOBP_CELL_PROJ | 1.50E-01 | 4.02E-01 | 0.1464 | -0.33297 | -1.28208 | 33 | Ripor2, .... |
| GOBP_CELL_PROJ | 7.99E-01 | 9.47E-01 | 0.0471 | -0.16955 | -0.82618 | 100 | Mef2c, C.... |
| GOBP_CELL_RECC | 3.63E-03 | 2.71E-02 | 0.4317 | -0.52961 | -1.8269 | 22 | Ighm, Co.... |
| GOBP_CELL_SUBS | 8.89E-01 | 9.76E-01 | 0.0434 | -0.18579 | -0.71536 | 33 | Coro1a, .... |
| GOBP_CELL_SURF | 4.77E-01 | 7.47E-01 | 0.0947 | 0.252726 | 0.97994 | 32 | Sox4, So.... |
| GOBP_CENTRAL_N | 2.56E-03 | 2.07E-02 | 0.4317 | 0.359001 | 1.72402 | 73 | Clu, S10.... |
| GOBP_CHEMICAL_ | 2.29E-01 | 5.18E-01 | 0.1421 | 0.238311 | 1.15517 | 76 | S100a9, .... |
| GOBP_CHROMATII | 9.29E-01 | 9.89E-01 | 0.0425 | -0.1828 | -0.63056 | 22 | Smchd1, .... |
| GOBP_CHROMATII | 5.84E-02 | 2.26E-01 | 0.2413 | -0.30609 | -1.37149 | 68 | Klf2, Sm.... |
| GOBP_CHROMATII | 9.67E-01 | 9.94E-01 | 0.0401 | -0.15094 | -0.55694 | 28 | Satb1, A.... |
| GOBP_CHROMOSC | 1.08E-01 | 3.36E-01 | 0.1709 | -0.26368 | -1.27026 | 92 | Sp100, K.... |
| GOBP_CHROMOSC | 4.38E-01 | 7.12E-01 | 0.0781 | -0.28161 | -1.02256 | 26 | Smc4, At.... |
| GOBP_CIRCADIAN_ | 3.68E-02 | 1.58E-01 | 0.3218 | -0.47169 | -1.52031 | 18 | Mycbp2, .... |
| GOBP_CIRCULATC | 3.75E-01 | 6.64E-01 | 0.1115 | 0.206736 | 1.04345 | 95 | Sox4, Kl.... |
| GOBP_CIRCULATC | 3.87E-01 | 6.74E-01 | 0.0824 | -0.27413 | -1.0451 | 31 | Stat1, K.... |
| GOBP_COGNITION | 5.60E-01 | 8.08E-01 | 0.0674 | -0.28341 | -0.93256 | 19 | Mef2c, B.... |

| | | | | | | | |
|---|---|---|---|---|---|---|---|
| GOBP_CYTOKINE_ | 9.46E-02 | 3.13E-01 | 0.1882 | -0.35911 | -1.38274 | 33 | Cd74, Il.... |
| GOBP_CYTOKINE_ | 5.82E-03 | 3.88E-02 | 0.407 | -0.36929 | -1.65247 | 67 | Malt1, C.... |
| GOBP_CYTOPLASI | 3.24E-01 | 6.15E-01 | 0.1172 | 0.242149 | 1.07795 | 51 | Ybx1, Yb.... |
| GOBP_CYTOSKELI | 6.94E-01 | 8.96E-01 | 0.0751 | 0.168997 | 0.90685 | 127 | S100a9, .... |
| GOBP_CYTOSOLIC | 2.42E-02 | 1.14E-01 | 0.3525 | -0.49145 | -1.58402 | 18 | Coro1a, .... |
| GOBP_DEFENSE_I | 1.97E-01 | 4.73E-01 | 0.1221 | -0.22984 | -1.16439 | 133 | Mef2c, I.... |
| GOBP_DEFENSE_I | 9.95E-02 | 3.21E-01 | 0.2193 | 0.38422 | 1.39562 | 24 | S100a9, .... |
| GOBP_DEFENSE_I | 1.38E-01 | 3.84E-01 | 0.1502 | -0.25727 | -1.24355 | 95 | Ighm, Ma.... |
| GOBP_DEFENSE_I | 3.26E-01 | 6.16E-01 | 0.0947 | -0.30366 | -1.11104 | 27 | Ptprc, S.... |
| GOBP_DENDRITE_ | 4.03E-01 | 6.87E-01 | 0.0848 | -0.32622 | -1.05144 | 18 | Mef2c, D.... |
| GOBP_DEPHOSPH | 3.45E-02 | 1.50E-01 | 0.3218 | -0.39697 | -1.52479 | 32 | Mef2c, P.... |
| GOBP_DETOXIFIC/ | 1.38E-06 | 5.47E-05 | 0.6436 | 0.686176 | 2.49242 | 24 | S100a9, .... |
| GOBP_DEVELOPM | 5.10E-01 | 7.70E-01 | 0.0858 | 0.27131 | 0.95692 | 21 | Spp1, So.... |
| GOBP_DEVELOPM | 6.90E-01 | 8.95E-01 | 0.0749 | 0.189445 | 0.86504 | 56 | Spp1, An.... |
| GOBP_DEVELOPM | 3.56E-01 | 6.44E-01 | 0.1071 | 0.303916 | 1.07192 | 21 | Spp1, So.... |
| GOBP_DEVELOPM | 5.91E-01 | 8.26E-01 | 0.0648 | -0.27635 | -0.90934 | 19 | Bcl11a, .... |
| GOBP_DEVELOPM | 9.23E-02 | 3.09E-01 | 0.2344 | 0.28149 | 1.32715 | 67 | Spp1, Nu.... |
| GOBP_DEVELOPM | 5.13E-01 | 7.71E-01 | 0.0858 | 0.295766 | 0.95906 | 16 | Nupr1, S.... |
| GOBP_DIVALENT_I | 9.76E-02 | 3.19E-01 | 0.225 | 0.358815 | 1.42022 | 33 | S100a9, .... |
| GOBP_DNA_CONF | 8.61E-01 | 9.69E-01 | 0.0437 | -0.19181 | -0.73677 | 32 | Smchd1, .... |
| GOBP_DNA_METAI | 5.90E-03 | 3.90E-02 | 0.407 | -0.36749 | -1.62426 | 62 | Il7r, Pt.... |
| GOBP_DNA_PACK/ | 5.34E-01 | 7.88E-01 | 0.0685 | -0.27478 | -0.94786 | 22 | Smchd1, .... |
| GOBP_DNA_RECO | 6.98E-03 | 4.34E-02 | 0.407 | -0.53001 | -1.76971 | 20 | Il7r, Pt.... |
| GOBP_DNA_REPAI | 1.22E-01 | 3.58E-01 | 0.1657 | -0.35841 | -1.32243 | 28 | Samhd1, .... |
| GOBP_DNA_REPLI | 6.03E-01 | 8.38E-01 | 0.0771 | 0.276415 | 0.89631 | 16 | S100a11,.... |
| GOBP_DOUBLE_ST | 3.04E-01 | 6.00E-01 | 0.1006 | -0.34659 | -1.14044 | 19 | Samhd1, .... |
| GOBP_ELECTRON_ | 3.42E-04 | 4.19E-03 | 0.4985 | 0.505539 | 2.02913 | 35 | Cyb5a, G.... |
| GOBP_EMBRYONI( | 2.97E-01 | 5.91E-01 | 0.1233 | 0.277735 | 1.11477 | 35 | Sox4, So.... |
| GOBP_EMBRYONI( | 4.16E-01 | 6.99E-01 | 0.0996 | 0.27472 | 1.03272 | 28 | Krt19, S.... |
| GOBP_EMBRYONI( | 3.86E-01 | 6.73E-01 | 0.1021 | 0.332256 | 1.05759 | 15 | Krt19, S.... |
| GOBP_EMBRYO_D | 6.09E-01 | 8.39E-01 | 0.0806 | 0.187814 | 0.91688 | 82 | Krt19, S.... |
| GOBP_EMBRYO_D | 1.70E-01 | 4.36E-01 | 0.1709 | 0.259394 | 1.19934 | 61 | Krt19, S.... |
| GOBP_ENDOCYTO | 1.66E-01 | 4.31E-01 | 0.1696 | 0.273807 | 1.21713 | 50 | Clu, Dmb.... |
| GOBP_ENDOMEME | 2.70E-01 | 5.66E-01 | 0.1315 | 0.279903 | 1.13244 | 37 | Clu, S10.... |
| GOBP_ENDOTHELI | 1.27E-01 | 3.65E-01 | 0.1644 | -0.39149 | -1.33421 | 21 | Mef2c, E.... |
| GOBP_ENDOTHELI | 3.49E-01 | 6.40E-01 | 0.0928 | -0.34431 | -1.10975 | 18 | Foxp1, B.... |
| GOBP_ENERGY_D | 2.73E-05 | 5.39E-04 | 0.5756 | 0.502404 | 2.19285 | 48 | Nupr1, N.... |
| GOBP_ENTRY_INT | 5.99E-01 | 8.34E-01 | 0.0644 | -0.27746 | -0.89429 | 18 | Cd74, H2.... |
| GOBP_ENZYME_LI | 3.73E-01 | 6.62E-01 | 0.1093 | 0.244172 | 1.06367 | 47 | Nedd4, C.... |
| GOBP_EPIDERMAL | 1.48E-03 | 1.33E-02 | 0.4551 | 0.61498 | 1.99415 | 16 | Reg3b, R.... |
| GOBP_EPIDERMIS_ | 1.35E-03 | 1.26E-02 | 0.4551 | 0.566956 | 2.02299 | 22 | Reg3b, R.... |
| GOBP_EPITHELIAL | 3.91E-01 | 6.78E-01 | 0.1028 | 0.285531 | 1.03714 | 24 | Sox9, Sp.... |
| GOBP_EPITHELIAL | 5.89E-04 | 6.34E-03 | 0.4773 | 0.422374 | 1.93485 | 58 | Reg3b, K.... |

e

| pathway | pval | padj | log2err | ES | NES | size | LeadingEdge |
|---|---|---|---|---|---|---|---|
| CAMP_UP.V1 | 8.59E-02 | 3.15E-01 | 0.2 | -0.394 | -1.4207 | 25 | Il7r, Pt.... |
| CAMP_UP.V1 | 3.71E-01 | 4.88E-01 | 0.107 | 0.2771 | 1.0771 | 32 | Mt2, Ifi.... |
| CYCLIN_D1_I | 2.05E-01 | 3.62E-01 | 0.125 | -0.402 | -1.2731 | 16 | Cd37, Ms.... |
| CYCLIN_D1_I | 9.67E-01 | 9.67E-01 | 0.04 | -0.164 | -0.5263 | 17 | Cd52, Tc.... |
| CYCLIN_D1_U | 3.94E-01 | 4.88E-01 | 0.104 | 0.3192 | 1.03292 | 18 | Mt1, Ned.... |
| CYCLIN_D1_U | 4.64E-01 | 5.49E-01 | 0.076 | -0.325 | -1.0054 | 15 | Cd52, Se.... |
| E2F1_UP.V1_ | 1.11E-01 | 3.15E-01 | 0.175 | -0.397 | -1.3557 | 21 | Ebf1, Kl.... |
| ERBB2_UP.V | 1.58E-01 | 3.15E-01 | 0.171 | 0.4322 | 1.33716 | 15 | S100a6, .... |
| ESC_V6.5_UF | 4.32E-06 | 5.62E-05 | 0.611 | 0.7183 | 2.39129 | 19 | S100a6, .... |
| GCNP_SHH_U | 6.33E-01 | 7.15E-01 | 0.06 | -0.27 | -0.8668 | 17 | Arhgef1,.... |
| GCNP_SHH_U | 7.30E-01 | 7.91E-01 | 0.054 | -0.23 | -0.8078 | 23 | Ptprc, P.... |
| HOXA9_DN.V | 1.44E-01 | 3.15E-01 | 0.18 | 0.4425 | 1.36888 | 15 | S100a9, .... |
| MTOR_UP.N4 | 4.79E-04 | 4.15E-03 | 0.498 | -0.599 | -1.9884 | 19 | Gimap6, .... |
| NFE2L2.V2 | 1.28E-01 | 3.15E-01 | 0.192 | 0.3623 | 1.34145 | 27 | Spp1, Nu.... |
| P53_DN.V1_U | 2.63E-11 | 6.83E-10 | 0.863 | 0.828 | 2.88561 | 22 | Krt18, T.... |
| PGF_UP.V1_U | 1.56E-01 | 3.15E-01 | 0.144 | -0.396 | -1.298 | 18 | 4930523C.... |
| PTEN_DN.V1_ | 2.74E-01 | 4.19E-01 | 0.105 | -0.369 | -1.1849 | 17 | H2-Eb1, .... |
| RB_P107_DN | 1.08E-03 | 6.99E-03 | 0.455 | 0.6308 | 1.97931 | 16 | S100a9, .... |
| RB_P130_DN | 3.93E-01 | 4.88E-01 | 0.102 | 0.3444 | 1.06545 | 15 | Clu, Eif.... |
| RPS14_DN.V | 3.04E-02 | 1.32E-01 | 0.352 | 0.5019 | 1.57475 | 16 | Spp1, Ma.... |
| SIRNA_EIF4G | 8.92E-01 | 9.27E-01 | 0.043 | -0.202 | -0.662 | 18 | H2-D1, Z.... |
| SNF5_DN.V1_ | 2.97E-03 | 1.54E-02 | 0.432 | -0.533 | -1.839 | 22 | Ripor2, .... |
| STK33_NOMO | 1.12E-01 | 3.15E-01 | 0.172 | -0.418 | -1.3712 | 18 | Il7r, St.... |
| STK33_SKM_ | 2.09E-01 | 3.62E-01 | 0.146 | 0.3783 | 1.2593 | 19 | Anxa1, E.... |
| STK33_UP | 2.68E-01 | 4.19E-01 | 0.128 | 0.352 | 1.18994 | 20 | Anxa1, J.... |
| TBK1.DF_DN | 3.31E-01 | 4.77E-01 | 0.093 | -0.29 | -1.0791 | 29 | Etnk1, T.... |

| pathway | pval | padj | log2err | ES | NES | size | LeadingEdge |
|---|---|---|---|---|---|---|---|
| HALLMARK_AD | 1.11E-02 | 3.44E-02 | 0.3807304 | 0.461 | 1.7145 | 26 | Gpx4, Uq.... |
| HALLMARK_ALl | 4.29E-10 | 1.20E-08 | 0.81403584 | -0.68 | -2.7185 | 37 | Cd79a, C.... |
| HALLMARK_AN | 1.62E-02 | 3.87E-02 | 0.35248786 | 0.521 | 1.7627 | 17 | Krt8, Kr.... |
| HALLMARK_AP | 3.26E-01 | 4.15E-01 | 0.11524 | 0.302 | 1.09 | 22 | Cdh1, Cl.... |
| HALLMARK_AP | 9.55E-03 | 3.34E-02 | 0.3807304 | 0.439 | 1.78 | 34 | Clu, Krt.... |
| HALLMARK_CO | 5.25E-03 | 2.45E-02 | 0.40701792 | 0.541 | 1.9205 | 20 | Clu, S10.... |
| HALLMARK_E2I | 9.14E-01 | 9.14E-01 | 0.04353782 | -0.19 | -0.6022 | 17 | Smc4, Pn.... |
| HALLMARK_EP | 7.82E-07 | 7.30E-06 | 0.6594444 | 0.742 | 2.601 | 19 | Spp1, Tr.... |
| HALLMARK_ES | 8.21E-03 | 3.28E-02 | 0.3807304 | 0.558 | 1.8483 | 16 | Krt8, Kr.... |
| HALLMARK_ES | 6.69E-04 | 3.75E-03 | 0.47727082 | 0.607 | 2.0983 | 18 | S100a9, .... |
| HALLMARK_G2 | 6.00E-01 | 6.47E-01 | 0.0618406 | -0.26 | -0.9203 | 24 | Ythdc1, .... |
| HALLMARK_GL | 4.51E-02 | 8.30E-02 | 0.32177592 | 0.442 | 1.5957 | 22 | Sox9, Tx.... |
| HALLMARK_HY | 2.70E-02 | 5.80E-02 | 0.35248786 | 0.447 | 1.6593 | 25 | Mt1, Mt2.... |
| HALLMARK_IL2 | 6.56E-02 | 1.08E-01 | 0.26635066 | 0.441 | 1.5455 | 19 | Spp1, If.... |
| HALLMARK_INT | 1.42E-01 | 2.10E-01 | 0.15419097 | -0.4 | -1.3164 | 18 | Cd74, Pa.... |
| HALLMARK_INT | 4.93E-06 | 3.45E-05 | 0.61052688 | -0.6 | -2.3047 | 31 | Cd74, H2.... |
| HALLMARK_KR | 1.66E-02 | 3.87E-02 | 0.35248786 | -0.51 | -1.6442 | 17 | Cd37, Et.... |
| HALLMARK_MIT | 4.14E-01 | 5.04E-01 | 0.08312913 | -0.32 | -1.0501 | 19 | Dock2, S.... |
| HALLMARK_MT | 2.18E-01 | 3.05E-01 | 0.14290115 | 0.287 | 1.1851 | 36 | Nupr1, P.... |
| HALLMARK_MY | 4.37E-01 | 5.10E-01 | 0.09754492 | 0.222 | 1.0269 | 59 | Eif3b, H.... |
| HALLMARK_OX | 2.24E-09 | 3.13E-08 | 0.77493903 | 0.592 | 2.6958 | 57 | Cyb5a, G.... |
| HALLMARK_P5 | 1.62E-02 | 3.87E-02 | 0.35248786 | 0.464 | 1.7052 | 23 | Nupr1, R.... |
| HALLMARK_PI3 | 4.75E-02 | 8.30E-02 | 0.2765006 | -0.44 | -1.4454 | 18 | Lck, Il2.... |
| HALLMARK_PR | 9.07E-02 | 1.41E-01 | 0.22496609 | 0.442 | 1.4491 | 15 | Krt18, C.... |
| HALLMARK_TN | 5.17E-01 | 5.79E-01 | 0.08756971 | 0.261 | 0.9649 | 24 | Pmepa1, .... |
| HALLMARK_UN | 6.57E-01 | 6.81E-01 | 0.07165274 | 0.254 | 0.8409 | 16 | Asns, Ee.... |
| HALLMARK_UV | 4.30E-02 | 8.30E-02 | 0.32177592 | 0.433 | 1.5489 | 21 | Mt1, Anx.... |
| HALLMARK_UV | 2.46E-01 | 3.29E-01 | 0.13574094 | 0.319 | 1.1795 | 24 | Epcam, C.... |
