## Supplementary Table 5 for "CREB drives acinar cells to ductal reprogramming and promotes pancreatic cancer progression in preclinical models of alcoholic pancreatitis"

**Supplementary Table 5.** Mouse Primers used in the study (Qiagen Catalogue#330001).

| Primers | Company | Gene Globe-ID |
|---|---|---|
| Amy2b | Qiagen | PPM66197A-200 |
| Ptf1a | Qiagen | PPM29825A-200 |
| Alb | Qiagen | PPM60179A-200 |
| Bhlh1a15 | Qiagen | PPM41737A-200 |
| Krt19 | Qiagen | PPM02968A-200 |
| Sox9 | Qiagen | PPM05134D-200 |
| Aldh1a3 | Qiagen | PPM31491A-200 |
| Spp1 | Qiagen | PPM03648C-200 |
| Tspan8 | Qiagen | PPM26276B-200 |
| Hes1 | Qiagen | PPM05647A-200 |
| Prss1 | Qiagen | PPM67018A-200 |
| Cpa1 | Qiagen | PPM26764A-200 |
